# Induced alanine auxotrophy as a therapeutic strategy against *Mycobacterium tuberculosis*

**DOI:** 10.64898/2026.06.12.731178

**Authors:** Menna-Allah W. Shalaby, Nagabhushana C. Beeralingappa, Annadka Shrinidhi, Gaelle Guiewi Makafe, Evan Nece, Aditya Patwardhan, Timothy Low-Beer, Atsuo Kuki, Felix Sheinerman, Brian Weinrick, Daniel P. Flaherty, Michaelle Chojnacki

**Affiliations:** Borch Department of Medicinal Chemistry and Molecular Pharmacology, College of Pharmacy, Purdue University, West Lafayette, Indiana 47907, United States; Trudeau Institute, Saranac Lake, NY, USA; Purdue Institute for Drug Discovery, West Lafayette, IN 47907, United States; Purdue Institute of Inflammation, Immunology and Infectious Disease, West Lafayette, IN 47907, United States

## Abstract

New antitubercular agents acting through previously unexploited mechanisms are urgently needed. Using a drug-repurposing platform, we identified TI-374, a hydroxamic acid containing compound that inhibits *Mycobacterium tuberculosis* (*Mtb*) with sub-micromolar potency. Systems analysis, resistance mapping, supplementation assays, and biochemical studies showed that TI-374 inhibits two PLP-dependent aminotransferases, AlaA and HisC1. However, its activity is driven primarily by irreversible inhibition of AlaA, whereas HisC1 inhibition is only partially reversible, revealing differential reversibility between the two targets. Optimization yielded TI-801, a low-nanomolar AlaA inhibitor. Both compounds remained active against intracellular *Mtb* in a macrophage infection model, where alanine supplementation did not rescue growth, indicating that host-derived alanine is unlikely to bypass AlaA inhibition. Genetic deletion of *alaA* attenuated *Mtb* survival in a murine infection model. Together, these findings support AlaA as a host-relevant metabolic vulnerability in *Mtb* and TI-801 as a mechanistic chemical probe for its validation as an antitubercular target.

## Introduction

The latest World Health Organization (WHO) global report estimates that 10.7 million people were diagnosed with tuberculosis (TB) and 1.23 million people succumbed to the disease in 2024.^1^ TB, caused by the bacterium *Mycobacterium tuberculosis (Mtb*), while briefly eclipsed by COVID-19, is once again the leading cause of death from an infectious agent worldwide.^1^ While TB mortality rates declined for much of the 21st century, the COVID-19 pandemic disrupted essential TB services, leading to increased transmission, delays in diagnosis, and worsening treatment outcomes.^2^ Emerging challenges—including antimicrobial resistance, healthcare access disparities, and ongoing geopolitical conflicts—threaten to further stall progress in TB control.^1,2^

A particularly alarming consequence has been the rise in multidrug-resistant TB (MDR-TB), which accounted for an estimated 400,000 infections, resulting in approximately 150,000 deaths in 2023.^1^ The growing incidence of MDR- and extensively drug resistant (XDR)-TB are partly attributable to the misuse of current antitubercular agents, and a direct consequence of the lengthy and complex nature of the treatment: a multidrug combination that typically must be taken for six months or longer. Owing to side effects and length, patients often do not adhere to therapy, with a consequent rise in MDR strains and cases of infection relapse.

In a recent WHO report describing the antibiotic pipeline, since 2017, 12 antibiotics have been approved, and 10 of these belong to existing classes with established mechanisms of antimicrobial resistance.^3^ While new drugs for the treatment of MDR TB – bedaquiline, delamanid and pretomanid – were introduced in the last 12 years, the burden of MDR/rifampicin-resistant TB as a share of all TB cases does not appear to be lessening in recent years and resistance to the newest drugs is emerging.^4–6^ In recent years, intensified research efforts have fueled the TB drug pipeline,^7–9^ however drugs that act via novel targets are still lacking.10

The discovery of therapeutics that act with novel mechanisms targeting previously unexploited pathways is one potential solution to overcome the emergence of resistant strains. While DNA replication, protein synthesis and cell wall biosynthesis have served as traditional targets for antibiotic development, essential metabolic pathways critical for cell growth and survival, particularly those involved in amino acid biosynthesis, have received comparatively little attention. A growing body of evidence indicates that auxotrophy for some amino acids renders *Mtb* less virulent and incapable of sustaining infection in the host lung environment.^8–13^ Therefore, amino acid biosynthetic pathways may represent a new strategy to target *Mtb* not only affecting bacterial survival directly but may also interfere with the ability of *Mtb* to adapt to the nutrient limitations encountered in host tissues.^11,12,14–19^ Furthermore, metabolic pathways are often highly conserved within a pathogen and essential under *in vivo* conditions. In recent years, increased attention to the cellular responses triggered by amino acid starvation has revealed vulnerabilities in *Mtb*’s stress response systems, further paving the way for rational, mechanism-based drug discovery.^10^ As such, the integration of metabolic targeting strategies with insights into nutrient sensing and host-pathogen interactions could provide a rich avenue for the development of durable, next-generation therapeutics.

While *Mtb* can import select amino acids such as aspartate, glutamate, asparagine, glycine and glutamine from the cytoplasm of infected macrophages,^20^ it has evolved to minimize reliance on the host by maintaining robust biosynthetic pathways for most other amino acids.^21^ This metabolic autonomy allows *Mtb* to survive and replicate in the nutrient-poor, hostile environment of host macrophages, where access to nutrients is limited by host defense mechanisms. Disruption of individual amino acid biosynthesis pathways, including those for leucine, methionine, arginine or histidine, has been shown to attenuate virulence and reduce bacterial burden in the lungs *in vivo*.^12,22–24^ Moreover, several enzymes involved in amino acid biosynthesis have emerged as promising drug targets in *Mtb,* offering a strategy to develop therapeutics with novel mechanisms of action. These pathways are often essential for *Mtb* survival in nutrient-limited host environments. To this end, inhibitors have been reported against the shikimate pathway, responsible for the synthesis of aromatic amino acids with antitubercular efficacy *in vitro* and *in vivo*.^25^ Similarly, the histidine biosynthetic pathway, essential under host-imposed nutrient limitation, has also emerged as a compelling target.^13,22,26^ Finally, TrpAB was genetically validated as essential for Mtb survival *in vivo* in a mouse infection model, and its allosteric inhibitor BRD4592 demonstrated on-target efficacy in a zebrafish infection model, although the inhibitor was not assessed in mice due to poor metabolic stability.^11^ Collectively, these developments underscore the therapeutic potential of targeting amino acid biosynthesis pathways in *Mtb*, which remain untapped by existing TB treatments.

Through a computational chemogenomics approach^27^ our team has identified a bioactive hit, PF-04859989 (TI-374), that appears to target the amino acid biosynthesis enzyme *Mtb* alanine aminotransferase, AlaA, encoded by *Rv0337c*. Although initially misannotated as an aspartate aminotransferase AspC,^28^ AlaA was identified as essential in an *in vitro* transposon mutagenesis screen.^29^ As it is essential under standard *in vitro* conditions, *Rv0337c* was absent from several *in vivo* transposon insertion screens,^30,31^ leaving its relevance during infection unclear. While the present work was underway, Edoo et al. recently independently reported that the related hydroxamic acid BVL3572S targets the same PLP-dependent aminotransferases AlaA and HisC in Mtb, corroborating AlaA as a vulnerable target from a distinct chemical starting point;^32^ here, we focus on the covalent mechanism, *in vivo* relevance, and AlaA-selective irreversibility of this engagement using TI-374 and TI-801.

TI-374 was originally in development for the treatment of cognitive impairment associated with schizophrenia^33–35^ and repurposing the scaffold represents a strategic opportunity to accelerate antibiotic development.^36^ Herein, we further elucidate the potency and underlying mechanism of its antitubercular activity. A combination of systems-level, biochemical, and microbiological evidence supports AlaA as a molecular target of TI-374, positioning this enzyme as a novel vulnerability in *Mtb*. To that end, our team has optimized the scaffold to produce a potent analog with low nanomolar activity against the pathogen. The studies reported herein contribute to the goal by identifying small molecule inhibitors with a novel mechanisms of action.

## Results

### TI-374 identification and activity against *Mtb*

TI-374 was identified as an anti-mycobacterial hit from the computational Trudeau Chemogenomic Discovery Platform (TCDP) at Trudeau Institute with a minimum inhibitory concentration (MIC) against *Mycobacterium abscessus* (*Mab*) of 35 µM.^27^ TI-374 (**Figure 1A**, reported previously as PF-04859989)^35^ was part of a curated virtual drug library of molecules, termed the Trudeau Virtual Drug Collection, that had reached advanced stages of preclinical and clinical development with known protein targets.The approach that identified the hit rested on a hypothesis that molecules inhibiting a known protein superfamily in humans could potentially cross-react with the same superfamily of enzymes available in mycobacteria. TI-374 was previously developed as an inhibitor against the human pyridoxal phosphate (PLP)-dependent enzyme kynurenine aminotransferase II (KAT II). Thus, having already demonstrated activity against *Mab*, we tested TI-374 for antitubercular activity. Susceptibility assays revealed that TI-374 had sub-micromolar potency (MIC = 0.54 μM) against *Mtb* H37Rv (**Figure 1B**). As MDR-TB, defined as *Mtb* resistant to standard of care drugs isoniazid (INH) and rifampicin (RIF), has dramatically increased in recent years, new drugs should be active against MDR strains. To this end, we confirmed that TI-374 retained activity against MDR-TB strain mc^2^8251, resistant to INH and RIF, at an MIC value of 0.31 μM. Further, pathogens in the *Mtb* complex (MTBC) are monomorphic with limited genetic variation between strains. However, several genotypic and phenotypic factors separate strains of the different MTBC lineages (L).^37^ While *Mtb* H37Rv is representative of European lineage (L4) and the most commonly used *Mtb* strain, we also evaluated activity against HN878 (L2, Beijing, hypervirulent) and TMC120 (L1, S. Indian).^37–40^ Against these strains TI-374 maintained comparable potencies at 1.88 and 1.25 μM, respectively (**Figure 1B**). To further characterize *in vitro* activity of TI-374 and to determine whether activity was bacteriostatic or bactericidal against *Mtb* H37Rv, time-kill analyses were performed at various multiples of the MIC. TI-374 displays bacteriostatic activity against *Mtb* H37Rv at 10× the MIC. At 100× the MIC, TI-374 was observed to be bactericidal, reducing viability to only 10 Log_10_ CFU/mL in 28 days (**Figure 1C**).

**Figure 1.**
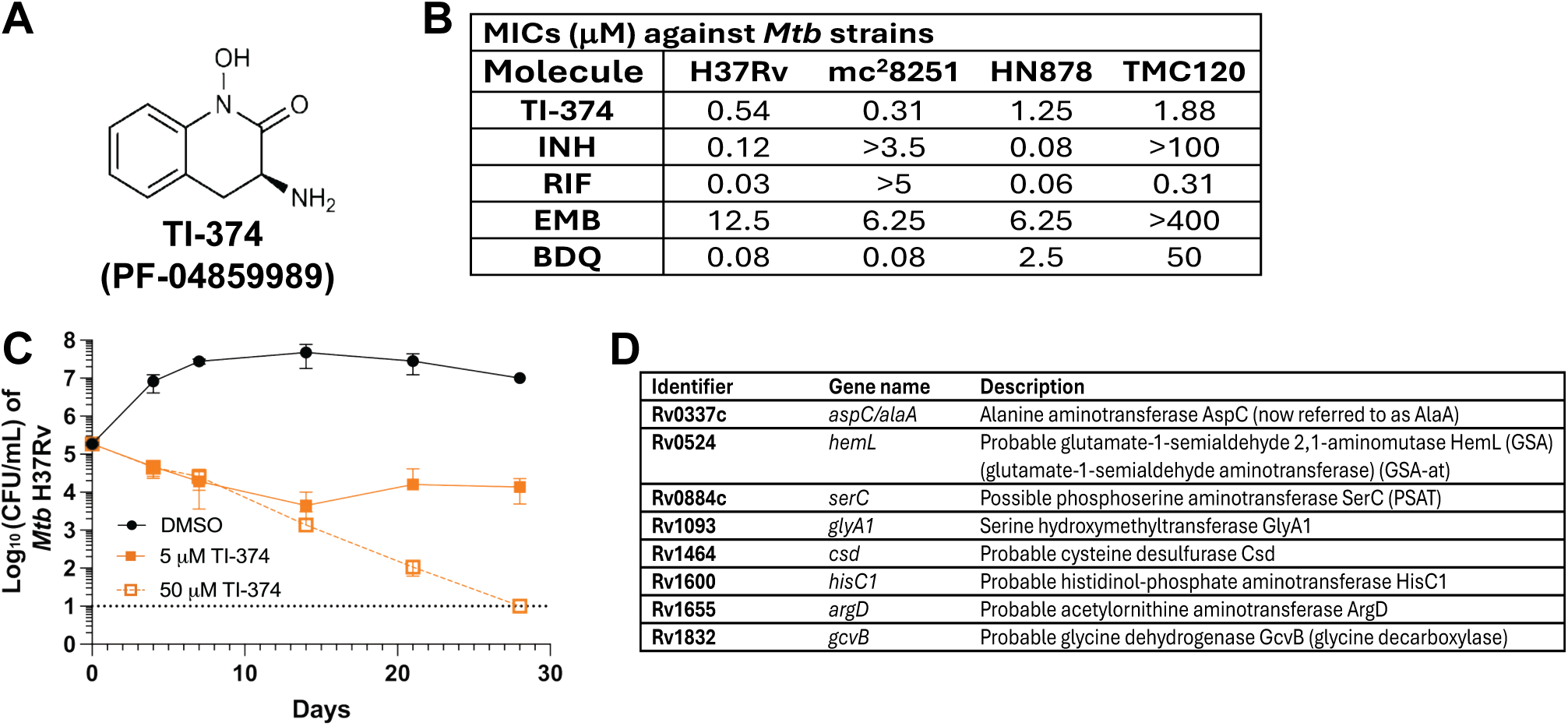
(**A**) Structure of TI-374 (PF-04859989). (**B**) MICs for TI-374 and control antibiotics versus *Mtb* strains presenting diverse lineages. INH = isoniazid, RIF = rifampin, EMB = ethambutol, BDQ = bedaquiline. The dashed line represents the limit of detection. (**C**) Time-kill curve for TI-374 against *Mtb* H37Rv strain at both 10x MIC (5 µM) and 100x MIC (50 µM). Data represented as mean ± SD of three biological replicates. (**D**) Protein products of essential H37Rv *Mtb* genes, containing enzymatic domains, which are members of the superfamily of PLP-dependent transferases

With TI-374 exhibiting potent anti-tubercular activity we began to narrow down a list of theoretical intracellular targets based on premise for cross-reactivity to PLP-dependent transferases. Structural predictions for *Mtb* proteins were conducted using Hidden Markov Models built based on structure-based sequence alignments of proteins with experimentally determined 3D structure.^41^ The structural domains belonging to the superfamily of PLP-dependent transferases were identified within protein products of 37 *Mtb* genes (**Table S1**). Eight out of the thirty-seven genes are annotated as essential for growth based on saturating transposon mutagenesis,^29^ thus representing possible targets for TI-374 (**Figure 1D**).

### Systems analyses indicate TI-374 treated *Mtb* modulates amino acid metabolism

Transcriptomic profiling of TI-374–treated Mtb (RNA-seq, two biological replicates) revealed a response dominated by a general amino acid starvation signature rather than a single-pathway readout. Of the 88 genes changed ≥2-fold, 49 reached significance (−log_10_ FDR > 1.3), with strong induction of *whiB7* and *lat* and coordinate repression of translation- and stress-associated genes (**Figure 2A**). This profile closely resembles the previously described *Mtb* nutrient-starvation response,^42^ an independent concordance that lends confidence to the overall signature despite the limited replicate number. The most statistically robust target-relevant signal was the strong induction of *whiB7*, the transcriptional regulator that coordinates the adaptive response to alanine starvation in mycobacteria reported by Poulton et al.,^43^ independently implicating perturbation of alanine metabolism. Both *alaA* (2.7-fold) and *hisC1* (2.3-fold) were themselves among the induced genes but did not reach the significance threshold, we therefore interpret the transcriptomic data as hypothesis generating, with target identification resting on the orthogonal metabolomic, genetic, supplementation, and resistance analyses described below.

**Figure 2.**
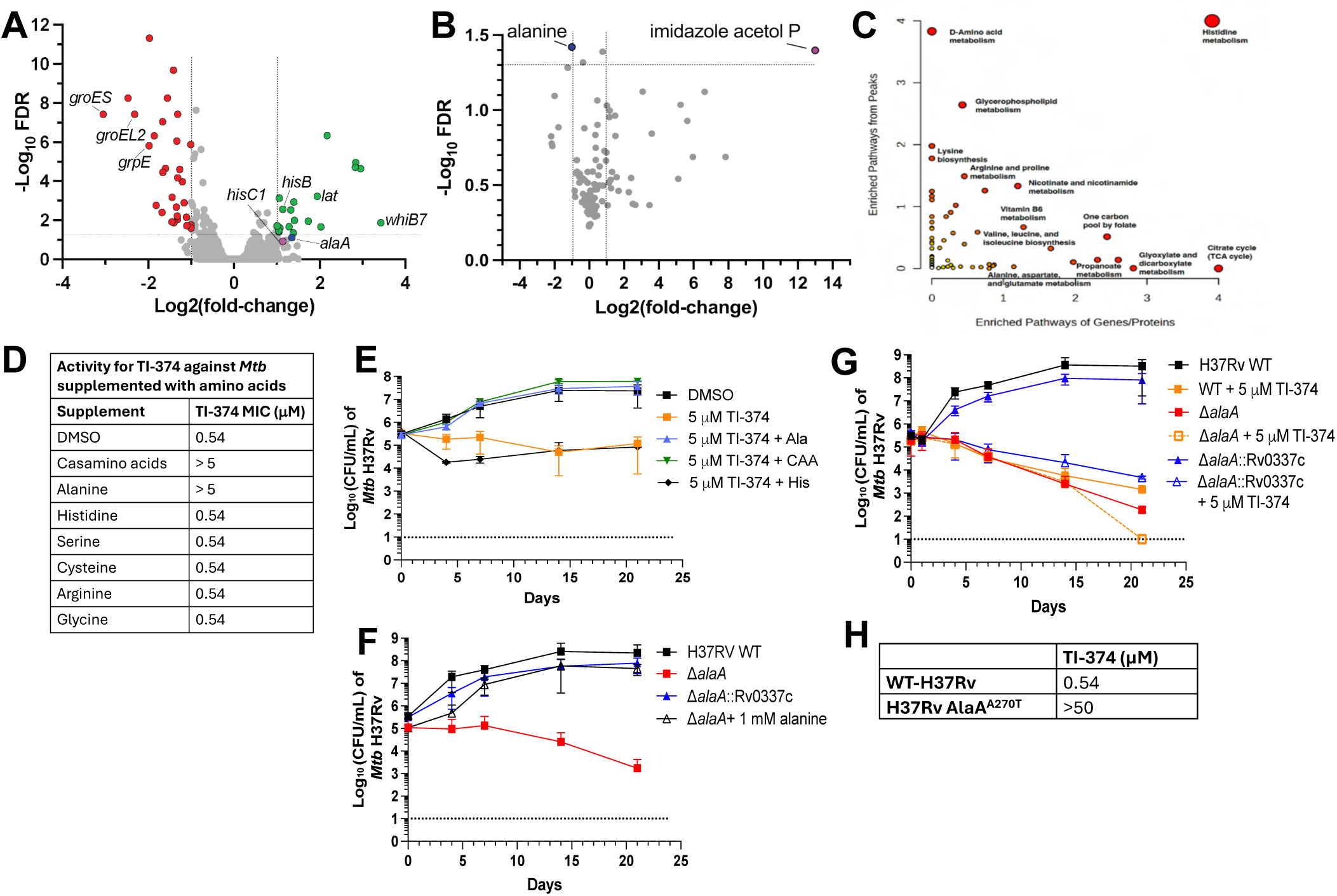
TI-374 activity is linked to alanine biosynthesis and AlaA function. Transcriptomic and metabolomic data suggest that TI-374 affects amino acid biosynthesis. (**A**) RNAseq analysis indicated group of upregulated and downregulated genes in *Mtb* H37Rv dosed at 5 μM (10x MIC) for 4 hours compared to vehicle treated control. Red = significantly down-regulated genes, green = significantly upregulated genes, blue = alaA gene, magenta = hisC1 gene. Statistical significance depicted in FDR = false discovery rate p < 0.05 (n = 2 biological replicates for each group). (**B**) Metabolomic evaluation of *Mtb* H37Rv dosed at 5 μM (10x MIC) for 4 hours. Change in metabolite profile of treated cells compared to vehicle control treated cells. Only metabolites that were confidently identified in all samples by mass spectrometry are plotted. Statistically significance depicted by FDR p < 0.05 (n = 3 biological replicates per group). (**C**) Joint Pathway Analysis in MetaboAnalyst (Version 6.0) using the Fisher’s p-value integration and significant peaks cutoff of 0.1. Scatterplot depicts the scaled enrichment of metabolic pathways based on metabolites (y-axis) and transcripts (x-axis). Selected enriched pathways are labeled. The size and color of data points is proportional to significance of merged GSEA and mummichog values. (**D**) MIC of TI-374 against H37Rv in the presence of individual amino acid supplements. Casamino acids were supplemented to 2 mg/mL, whereas alanine, histidine, arginine and serine were supplemented to 5 mM. Glycine was supplemented to 0.59 mM, cysteine to 0.1 mM and hemin to 0.03 mM. (**E**) Time-kill analysis of H37Rv treated with 5 µM TI-374 in the presence or absence of alanine (Ala), histidine (His) or casamino acids (CAA) supplemented to 5 mM. (**F**) Growth of WT, *ΔalaA*, and complemented (*ΔalaA*::*Rv0337c*) strains in 7H9 medium. (**G**) Time-kill analysis of WT, *ΔalaA*, and complemented strains treated with TI-374. (**H**) MIC of TI-374 against WT H37Rv and a resistant mutant harboring an AlaA Alanine to threonine substitution at position 270 (A270T). All growth curve data represented as mean ± SD of three biological replicates.

To examine the effect of TI-374 on the abundance of alanine, histidine, and other metabolites that may be relevant to its antitubercular activity, we conducted untargeted metabolomic profiling by LC-MS. As with the RNA sequencing analysis, *Mtb* H37Rv was treated for 4 hours at 10-fold the MIC of TI-374 in triplicate before being processed for LC-MS. The analysis detected more than 3,000 accurate mass retention time pair ions between the positive and negative modes. From these, 99 of these metabolites were identified with high confidence based on standards and database matches, and these are plotted as differentially identified metabolites compared to DMSO treated controls (**Figure 2B**). Among these 99 metabolites only two exhibited both a differential fold-change > 2 and Log_10_ FDR > 1.3, alanine and imidazole acetol phosphate. Alanine is the product of the AlaA while imidazole acetol phosphate is a substrate of HisC1. Joint Pathway Analysis of the metabolomic data with the transcriptomic data using MetaboAnalyst 6.0^44^ revealed amino acid metabolism as the most significantly perturbed pathways (**Figure 2C**). Specifically, histidine metabolism was substantially altered in both the transcriptome and the metabolome. Interestingly, despite being one of the hypothesized pathways targeted by TI-374, alanine metabolism exhibited low significance in this result. This may simply be a consequence of limitations of metabolic pathway annotation and/or associated statistical tests. Alanine is produced in a single step by AlaA from pyruvate accounting for only two metabolites and one enzyme among more than twenty annotated in the alanine, aspartate, and glutamate metabolic pathway. Thus, changes specifically to alanine metabolism may not be readily identified among the rest of the related aspartate and glutamate signals. Conversely, apparent inhibition of HisC1 perturbs expression of four genes and six metabolites, a much larger percentage of the annotated products in the similarly complex histidine biosynthetic pathway providing a greater signal.

Indeed, the effect of TI-374 on alanine levels is clearer when accounting for the statistical significance of the “D-amino acid metabolism” pathway. Our LC-MS method is unable to distinguish enantiomers, so the significance of that pathway may be driven by perturbation of the levels of L-alanine, L-lysine, L-glutamate, L-glutamine, L-arginine, L-proline, and L-ornithine. The effect of TI-374 on alanine biosynthesis is further supported by marked depletion of alanine, ∼2 to 3-fold below control samples, coupled with a slight increase in its substrate pyruvate levels. Taken together, the data indicate alanine biosynthesis, and specifically AlaA, may be a direct target of TI-374.

The inhibition of HisC1 activity is specifically supported by the substantial accumulation of its substrate imidazole-acetol phosphate, which was not detected in the control samples but had accumulated to significantly high levels in the TI-374 treated samples, with observed intensities similar to several high-abundance citrate cycle intermediates. Overall, these results are consistent with TI-374 interfering with amino acid biosynthesis to chemically induce an auxotrophy.

### TI-374 inhibits AlaA as intracellular mechanism of action

Based on the systems-level data indicating modulation of amino acid biosynthetic pathways, we evaluated the effect of amino acid supplementation on the activity of TI-374 against *Mtb*. Supplementation with casamino acids significantly reduced susceptibility to TI-374, shifting the MIC from 0.54 μM to > 5 μM (**Figure 2D**). This observation prompted exploration into specific amino acid(s) that could account for this effect. Considering the chemogenomics target list and the systems data, we supplemented media individually with alanine, histidine and the amino acids associated with the other potential targets from **Figure 1**. Alanine supplementation alone was sufficient to abrogate TI-374 activity, shifting the MIC > 5 μM (**Figure 2D**). While *hisC1* was also induced upon TI-374 treatment, histidine alone did not affect the MIC of TI-374, nor did any other amino acid supplemented. Additionally, TI-374 contains a hydroxamate moiety that is known to chelate metals, particularly iron and zinc.^45,46^ To rule out metal chelation as a contributing factor for antimicrobial mechanism of action we tested the activity of TI-374 in the presence of iron, zinc, magnesium, calcium and copper. The MIC for TI-374 did not shift in the presence of any metal (**Table S2**).

To validate this phenotype under dynamic conditions, a checkerboard assay with alanine supplementation demonstrated the strain could be rescued with as low as 20 μM alanine (**Figure S1**). Time-kill assays were performed against *Mtb* H37Rv treated with TI-374 at 10× MIC in the presence or absence of alanine, casamino acids or histidine (**Figure 2E**). The results corroborate the MIC data and demonstrate that supplementation with alanine or casamino acids can restore *Mtb* growth to levels comparable to DMSO-treated control. The specificity of this rescue suggests that TI-374 activity is linked to disruption of alanine biosynthesis, and potentially targets AlaA, one of the computationally predicted targets from **Figure 1D**.

To define the physiological role of AlaA, and directly interrogate the contribution of alanine biosynthesis to growth and TI-374 susceptibility, we generated an Δ*alaA* mutant in H37Rv and compared growth of WT, Δ*alaA*, and the complemented strain under standard *in vitro* conditions (**Figure 2F**). The auxotrophic Δ*alaA* strain exhibited a pronounced and progressive growth defect in 7H9 medium, consistent with impaired endogenous alanine biosynthesis. The defect was fully rescued by supplementation with exogenous alanine, restoring growth to near WT levels. In contrast, WT and complemented strains grew robustly irrespective of alanine availability. Together, these data establish AlaA as a key contributor to alanine biosynthesis and demonstrate that this metabolic requirement can be bypassed by exogenous alanine.

To determine whether pharmacological inhibition of AlaA phenocopies genetic disruption, we evaluated the activity of TI-374 across WT, Δ*alaA,* and complemented strains (**Figure 2G**). Treatment of WT with TI-374 resulted in a pronounced growth defect, closely recapitulating the Δ*alaA* phenotype. Notably, the Δ*alaA* strain exhibited markedly enhanced susceptibility to TI-374, with viability declining toward the limit of detection over time. In contrast, the complemented strain displayed partial restoration of growth and reduced sensitivity to treatment. While the heightened sensitivity of the Δ*alaA* mutant suggests that TI-374 exerts additional effects beyond AlaA inhibition alone, supplementation with alanine, but not other amino acids, alleviated TI-374 activity, functionally linking drug susceptibility to disruption of alanine metabolism. Together, these data identify AlaA as a key determinant of TI-374 sensitivity and support alanine biosynthesis as a primary pathway-level vulnerability targeted by the compound.

Given the strong genetic and pharmacologic evidence linking AlaA to TI-374 susceptibility, we next sought independent evidence of target engagement by selecting for resistant mutants under TI-374 pressure. To do so, 10^9^ *Mtb* H37Rv bacterial cells were plated on 7H10 agar containing 40× MIC TI-374 (20 µM) and plates were incubated until the appearance of colonies. The choice to use a high MIC multiple was based on prior literature suggesting this would increase the probability of selection of mechanism-relevant mutations only.^47^ Colonies arose at a frequency of ∼2x10^-8^, a value comparable to the spontaneous resistance frequencies reported for frontline TB drugs such as rifampicin.^48,49^ Isolation and subsequent susceptibility testing confirmed the resistant isolate had acquired reduced susceptibility to TI-374 with MIC value 50 μM, a 100-fold shift in potency compared to wild-type *Mtb* H37Rv (**Figure 2H**). The DNA was extracted and whole genome sequencing data was collected and analyzed. The resulting sequencing analysis identified a mutation in the *Rv0337c/aspC(alaA)* gene which resulted in Ala270Thr substitution within the protein, supporting the intracellular target for TI-374 is AlaA.

### In vitro inhibition support AlaA as target

To directly evaluate the inhibitory activity of TI-374 against AlaA, enzymatic assays were performed. Recombinant AlaA was expressed in *Escherichia coli*, purified, and prepared for biochemical analysis. Enzymatic activity was monitored using a lactate dehydrogenase-coupled continuous assay, in which the reduction of pyruvate (the AlaA reverse reaction product) to lactate results in stoichiometric consumption of NADH tracked at absorbance 340 nm, as previously described.^28^ TI-374 was shown to inhibit recombinant AlaA in a dose-dependent manner, with an IC_50_ value of 23 ± 8 nM, consistent with its potent antibacterial activity (**Figure 3A**). Histidine biosynthesis was also identified in the systems analysis as a potential pathway that was perturbed by TI-374 treatment, yet histidine supplementation did not rescue inhibition in *Mtb*. For completeness, we evaluated TI-374 for activity against HisC1 to determine if the compound can inhibit the enzyme. Our data indicate that TI-374 exhibits an *Mtb* HisC1 IC_50_ value of 619 ± 74 nM, approximately 25-fold weaker than against AlaA (**Figure 3A**).

**Figure 3.**
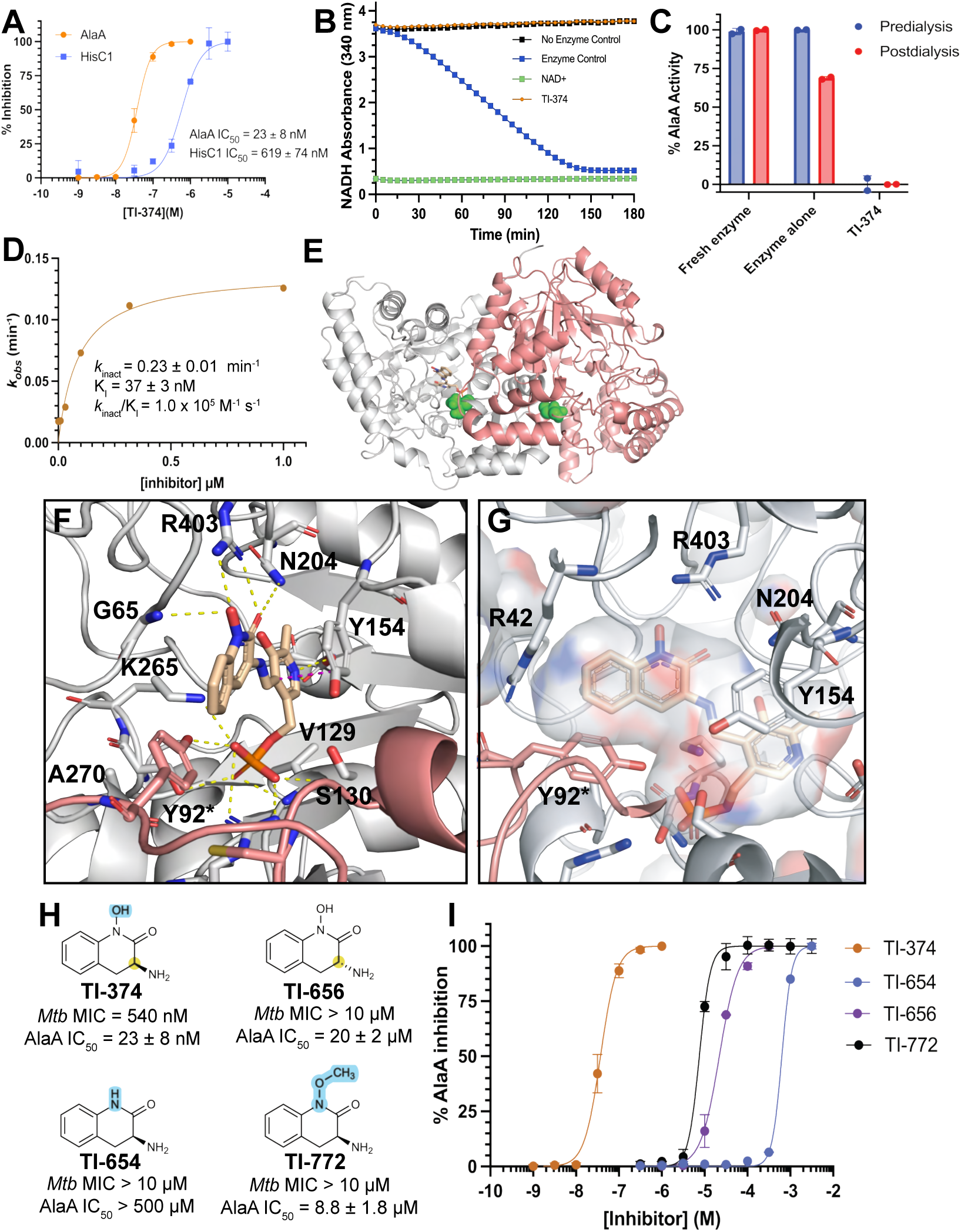
Biochemical and predicted structural analysis of TI-374 scaffold against *Mtb* AlaA. (**A**) Inhibition of *Mtb* AlaA and HisC1 with TI-734 plotted as normalized percent inhibition is each assay represented as mean ± SD, n = 9. (**B**) *Mtb* AlaA continuous assay run in jump-dilution format with TI-374, represented as mean ± SD, n = 3. (**C**) Dialysis study with percent *Mtb* AlaA activity with and without TI-374 pre- and post-dialysis, presented as mean ± SD, n = 2. (**D**) Kinetics of covalent inhibition for TI-374 against *Mtb* AlaA from progress curves performed in n = 3. (**E**) Homology model of *Mtb* AlaA depicting dimeric quaternary structure consisting of two monomeric units (Chain A = gray cartoon; Chain B = salmon cartoon). Ala270 highlighted by green spheres. (**F** and **G**) Predicted binding pose for TI-374 (gold sticks, represented as the PLP-adduct) to *Mtb* AlaA. Residues relevant to ligand binding site shown as sticks and labeled. Residues with * are from Chain B. Predicted hydrogen bonds depicted by yellow dashes. Ligand binding site surface depicted in **G**. (**H**) Analogs designed to probe predicted binding interactions with corresponding activity against *Mtb* AlaA and MICs with values reported as mean ± SD, n = 9. Regions highlighted that were probed by the corresponding modifications. (**I**). Representative dose-response curves for analogs against *Mtb* AlaA represented as mean ± SD.

Based on prior reports for TI-374 inhibition of KAT II,^34^ we hypothesized the molecule acts as a mechanism-based inhibitor via covalent modification of the PLP cofactor within the AlaA active site, leading to irreversible enzyme inactivation. To evaluate reversibility, a jump dilution assay was performed^50^ in which TI-374 was preincubated with AlaA at 10× IC_50_ for 1 hour, followed by rapid 100-fold dilution into assay buffer containing substrates (L-alanine and α-ketoglutarate), PLP, and a coupling system consisting of LDH and NADH, resulting in a final inhibitor concentration of 0.1× IC_50_. Enzymatic activity was monitored over 180 minutes and compared to an enzyme-only control. Under these conditions, reversible inhibitors are expected to dissociate upon dilution, leading to recovery of enzymatic activity, whereas irreversible inhibitors maintain sustained suppression of activity.^50^ Consistent with irreversible inhibition, TI-374 showed no recovery of activity following dilution, as evidenced by a flat progress curve with no detectable product formation (**Figure 3B**). To further confirm irreversibility, an orthogonal dialysis experiment was conducted.^51^ AlaA (1 μM) was preincubated with TI-374 (100 μM) or vehicle control for 1 hour at room temperature, then transferred to a dialysis cassette and dialyzed against 2000 mL of PBS containing 10% glycerol and 150 μM PLP. The buffer was changed after 6 hours, and PLP was included to preserve enzyme stability during dialysis. After 18 hours, the dialyzed samples were retested for activity and compared to freshly prepared enzyme controls. The enzyme preincubated with TI-374 showed complete loss of activity, and following dialysis, retained only ∼5% activity, indicating no recovery. In contrast, the control enzyme alone samples maintained 70% activity throughout the experiment (**Figure 3C**). Together, these results demonstrate that TI-374 inhibition of AlaA is irreversible.

To gain deeper insight into the mechanism of inhibition beyond biochemical potency, which does not account for time-dependent inactivation, we characterized the kinetic inhibition of AlaA by TI-374 using progress curve analysis. Enzymatic progress curves were monitored at varying inhibitor concentrations and fitted to an exponential decay equation (**Figure S2**) to extract observed inactivation rate constants (*k*_obs_) at each concentration. The *k*_obs_ values were then plotted against inhibitor concentration and fitted to a hyperbolic equation corrected for substrate competition (L-alanine) to determine the maximum inactivation rate constant (*k*_inact_) and the apparent dissociation constant (*K*_i_), following a previously reported approach.^52^ As shown in **Figure 3D**, TI-374 demonstrated potent time-dependent inhibition of AlaA, with a *K*_i_ of 37 ± 3 nM and a *k*_inact_/*K*_i_ ratio of 1.0 × 10^5^ M^-1^s^-1^. The high *k*_inact_/*K*_i_ value reflects efficient enzyme inactivation, representing the second-order rate constant that collectively describes initial enzyme–inhibitor complex formation and subsequent enzyme inactivation. Taken together, these kinetic parameters establish TI-374 as a highly potent mechanism-based inhibitor of AlaA, providing quantitative evidence for its time-dependent, irreversible mode of inactivation.

### Structural hypothesis for TI-374 binding to AlaA

Since the crystal structure of *Mtb* AlaA has not yet been resolved, we generated a high-quality atomic model using established homology modeling techniques, starting from a known crystal structure of closely related pyridoxal phosphate (PLP)-dependent transaminases. Based on a BLAST search, *E. coli* AlaA (*Ec*AlaA, UniProt ID: P0A959; PDB: 4CVQ),^53^ which shares 59.16% sequence identity with *Mtb* AlaA (UniProt ID: P9WQ91; **Figure S3**), was selected as the modeling template. Generating a model from the homodimeric structure was necessary to preserve the features of the active site. The model passed quality metrics when aligned with *Ec*Ala producing an RMSD of 0.263 Å. Model validation was performed using a Ramachandran plot to assess backbone dihedral angles (phi/psi) and revealed all residues were located within favored or allowed regions, with none in disallowed regions, indicating good stereochemical quality and overall model stability (**Figure S3**). Active site residues and residues engaged in binding of PLP cofactor in *Ec*AlaA are fully conserved in *Mtb* AlaA (**Figure S4**). Analysis of the *Mtb* AlaA model suggests both monomers contribute to the catalytic site, consistent with what is observed in *Ec*AlaA.

We performed molecular docking using Glide (Schrodinger, LLC). The crystal structure of TI-374 bound to human KAT II (PDB: 3UE8)^35^ depicts the molecule as a PLP adduct in the active site. We sought to evaluate the binding interactions of the molecule within *Mtb* AlaA active site in complex with PLP. Therefore, we generated the PLP-TI-374 adduct to be used for docking analysis. The top three poses in terms of binding score overlapped almost identically within the *Mtb* AlaA binding site and made the same interactions with the protein. Pose 1 was selected for describing interactions and used to illustrate the predicted binding mode (**Figure 3F**). Importantly, the same interactions are observed to the PLP-portion of the molecule in *Mtb* AlaA as were observed in *Ec*AlaA and with the TI-374 adduct in KAT II. Three critical residues were shown to engage the TI-374 hydroxamate moiety, those being the backbone amide of Gly65 and side chains of Asn204 and Arg403 (**Figure 3F**) with the latter two side chains also shown to be critical to TI-374 binding in human KAT II (Asn202 and Arg399, **Figure S3**). The binding site is flanked by Arg42 from chain A (hidden in **Figure 3F**; shown in **Figure 3G** and **Figure S3**) and Tyr92* from chain B. While *Ec*AlaA structure does not have TI-374 bound it does have an acetate molecule within the active site that overlays with the hydroxamate from TI-374 (**Figures S4**).^54^ *Ec*AlaA also has identical corresponding residues Gly41, Asn179 and Arg378 that are shown to engage the acetate in the same manner that they engage the hydroxamate from TI-374 in the *Mtb* AlaA model.

Next, we used the homology model to gain insight into the contributions of the Ala270Thr mutation that inferred reduced susceptibility to TI-374. The binding pocket for TI-374 is a narrow tunnel formed partially by Tyr92* from chain B of *Mtb* AlaA on the bottom half wall (**Figure 3F** and **G**). Ala270 is fully buried within the Mtb AlaA structure and is packed against Tyr92* (**Figure 3F** and **S5**). The Ala270Thr mutation was modeled using the mutagenesis wizard in PyMol (V3.1.8; Schrödinger, LLC). Two possible rotamers of Thr270 were obtained (Figure SX-3), both predicted to generate steric clashes with surrounding residues via the methyl group — rotamer 1 with Pro67 and rotamer 2 with Tyr92* — suggesting this substitution could disrupt the binding pocket geometry and prevent TI-374 engagement with PLP.

### Initial SAR supports proposed TI-374 binding mode

To explore the binding mode targeted analogs of TI-374 wsgs were designed: one lacking the *N*-hydroxyl group (TI-654) and one bearing alkylation of the hydroxyl group (TI-772). Both modifications resulted in diminished activity, with MIC values >10 µM. Enzymatic assays further confirmed markedly reduced potency, with IC_50_ values of 630 µM (TI-654) and 7.5 µM (TI-772), collectively supporting the importance of an intact hydroxamate for maintaining key interactions with Asn204 and Arg403 (**Figure 3H** and **I**).

Next, the role of the hydroxamic acid moiety was examined. Docking analysis of TI-374 revealed that the hydroxamate engages in key interactions with residues Asn204 and Arg403, both essential for activity. To probe the importance of these interactions, two analogs were designed: one lacking the *N*-hydroxyl group (TI-654) and one bearing alkylation of the hydroxyl group (TI-772). Both modifications resulted in diminished activity, with MIC values >10 μM. Enzymatic assays further confirmed markedly reduced potency, with IC_50_ values of 630 μM (TI-654) and 7.5 μM (TI-772), collectively supporting the importance of an intact hydroxamate for maintaining key interactions with Asn204 and Arg403 (**Figure 3H** and **I**).

### TI-801 exhibits improved activity against AlaA and *Mtb*

Early design of novel analogs to improve activity yielded TI-801 with a *Mtb* H37Rv MIC of 93 nM, representing an approximately 6-fold improvement over TI-374, and a *Mtb* AlaA IC_50_ of 10.1 ± 0.3 nM, and 2-fold improvement in biochemical potency (**Figures 4A** and **B**). TI-801 retains the core hydroxamic acid scaffold and the *S*-configured amine of TI-374 while incorporating a chlorine substituent at the 6-position of the aromatic ring. TI-801 was evaluated using a jump-dilution assay and, consistent with the parent compound, no recovery of enzymatic activity was observed following rapid dilution (**Figure 4C**) suggesting TI-801 functions as an irreversible inhibitor of *Mtb* AlaA. When evaluated for HisC1 activity, the analog exhibited an IC_50_ value of 232 ± 76 nM (**Figure S6**), approximately 23-fold less active than against *Mtb* AlaA.

**Figure 4.**
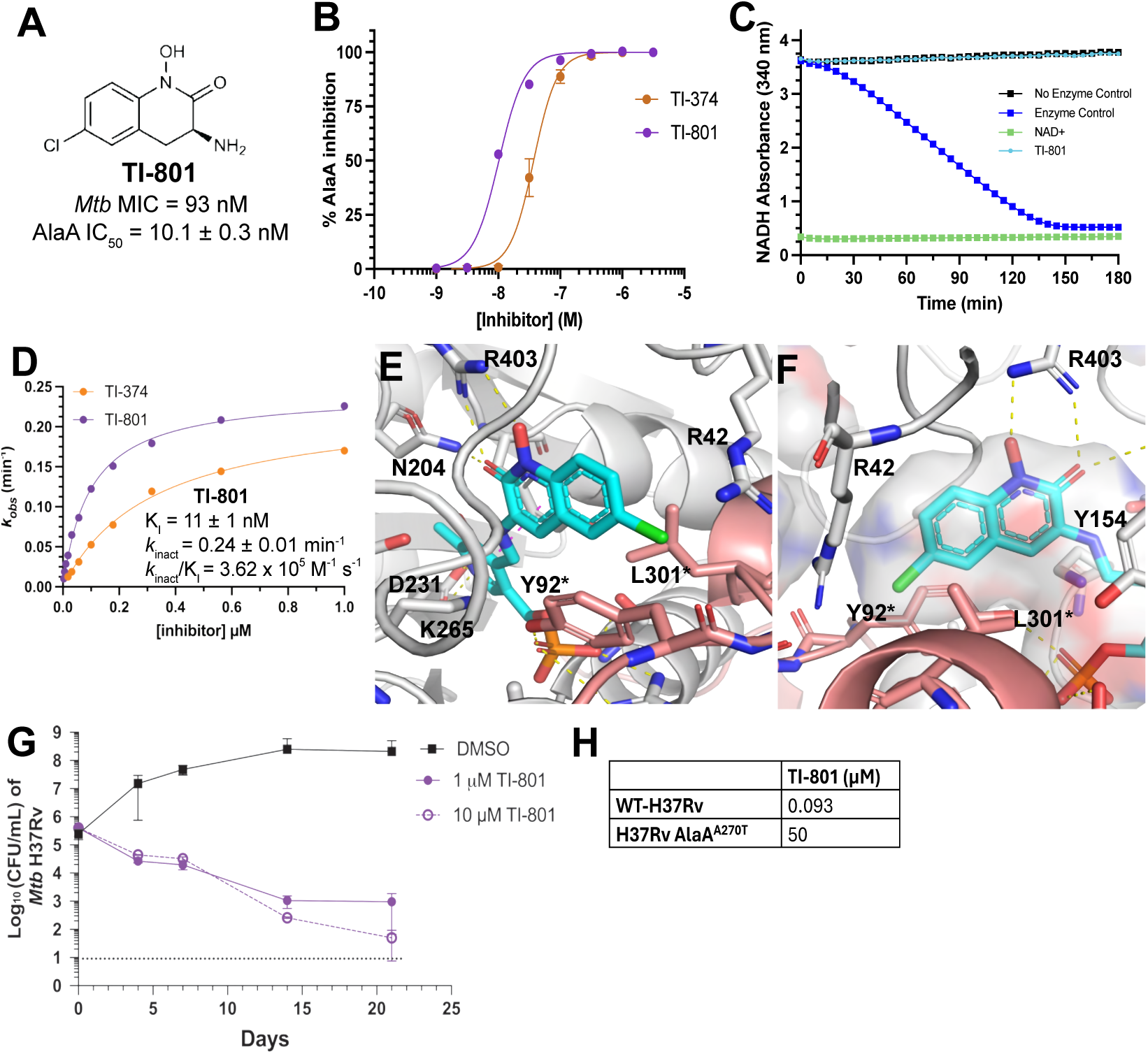
Biochemical and micrbiological evaluation of TI-801. (**A**) Structure and potency data for TI-801. (**B**) Overlaid dose-response curves for TI-374 and TI-801 against *Mtb* AlaA reported as mean ± SD, n = 3. (**C**) *Mtb* AlaA continuous assay run in jump-dilution format with TI-801 reported as mean ± SD, n = 2. (**D**) Kinetics of covalent inhibition for TI-801 against *Mtb* AlaA, overlaid with TI-374 reported as mean ± SD, n = 3. (**E** and **F**) Predicted binding pose of TI-801 (cyan sticks shown as the PLP-adduct). Residues relevant to the binding site shown as sticks and labeled. Residues with * indicates Chain B. Hydrogen bonds depicted as yellow dashed lines. Surface of ligand binding site shown in **F**. (**G**) Time-kill curve for TI-801 against *Mtb* H37Rv strain at both 10x MIC (1 µM) and 100x MIC (10 µM) reported as mean ± SD, n = 3 biological replicates. (**H**) MICs for TI-801 against *Mtb* H37Rv strain and H37Rv AlaA^A270T^ mutant strain.

Kinetic characterization by progress curve analysis (**Figure S7**) indicates that TI-801 is a potent time-dependent inactivator of AlaA, with a *K*_i_ of 11 ± 1 nM, a *k*_inact_ of 0.24 ± 0.01 min⁻¹, and a *k*_inact_/*K*_i_ of 3.6 × 10^5^ M^-1^ s^-1^ (**Figure 4D**). Compared to TI-374, this represents an approximately 3-fold improvement in binding affinity and a 4-fold improvement in overall inactivation efficiency. Notably, the *k*_inact_ values for the two compounds are similar, suggesting that the enhanced efficiency of TI-801 is driven primarily by tighter initial binding rather than a faster chemical inactivation step.

To rationalize the improved potency of TI-801, the TI-801–PLP adduct was docked into the *Mtb* AlaA homology model. The predicted pose recapitulates the binding mode proposed for TI-374, with the hydroxamate engaging Asn204 and Arg403 and the PLP moiety occupying the phosphate-binding region (**Figure 4E** and **F**). In this pose the 6-chloro substituent projects into the narrow channel formed by Leu301* and Tyr92* of the opposing monomer and is oriented toward the guanidinium of Arg42. Introduction of this substituent is accompanied by a modest improvement in AlaA affinity (Kᵢ 37 to 11 nM) and a corresponding gain in whole-cell potency (MIC 540 to 93 nM), consistent with a favorable contact between the chlorine and Arg42 in this region of the pocket. The precise nature and energetic contribution of this interaction, together with a systematic structure–activity analysis of substituents at this position will ultimately require a high-resolution co-crystal structure of TI-801 bound to Mtb AlaA.

Time-kill studies revealed dose-dependent bactericidal activity, with concentrations of 1 µM (10× MIC) and 10 µM (100× MIC) TI-801 resulting in progressive reductions in bacterial burden over 21 days, surpassing the activity of TI-374 at concentrations of 5 µM and 50 µM (**Figure 4G**). Additionally, to assess whether the A270T resistance mutation confers cross-resistance to TI-801, the compound was evaluated against *Mtb* H37Rv harboring the *Mtb* AlaA A270T mutation. Consistent with TI-374, TI-801 exhibited a reduced activity against the mutant strain, with an IC_50_ of 50 µM compared to 93 nM in WT H37Rv, representing a greater than 500-fold shift in potency (**Figure 4H**). This cross-resistance profile strongly implicates inhibition of AlaA as the primary mechanism-of-action for this inhibitor class and confirms that the A270T substitution is a shared resistance determinant across these analogs.

### Differential reversibility distinguishes AlaA and HisC1 engagement

To compare how the scaffold engages the two PLP-dependent targets, we evaluated TI-374, TI-801, and the related congener BVL3572S against both *Mtb* AlaA and HisC1 (**Figure 5A**). First, we carried out biochemical evaluation of BVL3572S for both AlaA and HisC1 to establish a potency baseline with AlaA IC_50_ = 53 ± 9 nM and HisC1 activity was 3.5-fold weaker with IC_50_ = 195 ± 73 nM (**Figure 5A** and **B**). We next assessed reversible inhibition in a jump-dilution format under matched conditions (preincubation at 10× IC_50_, 100-fold dilution to 0.1× IC_50_). Against AlaA, all three compounds demonstrated no recovery of activity following dilution, consistent with irreversible inactivation independent of the aryl substituent (**Figure 5C**). In contrast, HisC1 inhibition showed to be reversible, and the extent of recovered activity tracked with the aryl substituent and placement in which TI-374 (H) recovered 30 ± 5% of activity, TI-801 (6-Cl) 65 ± 7%, and BVL3572S (7-Br) recovered 83 ± 7% activity compared to control (**Figure 5D**). These results indicate that the kinetic mode of inhibition is governed by the enzyme context rather than by the warhead itself. The same PLP adduct chemistry is kinetically locked within the AlaA active site but is reversible, and tunable by peripheral substitution, within HisC1. This suggests the committed inhibitory step for all three compounds is therefore on AlaA, whereas HisC1 engagement is transient.

**Figure 5.**
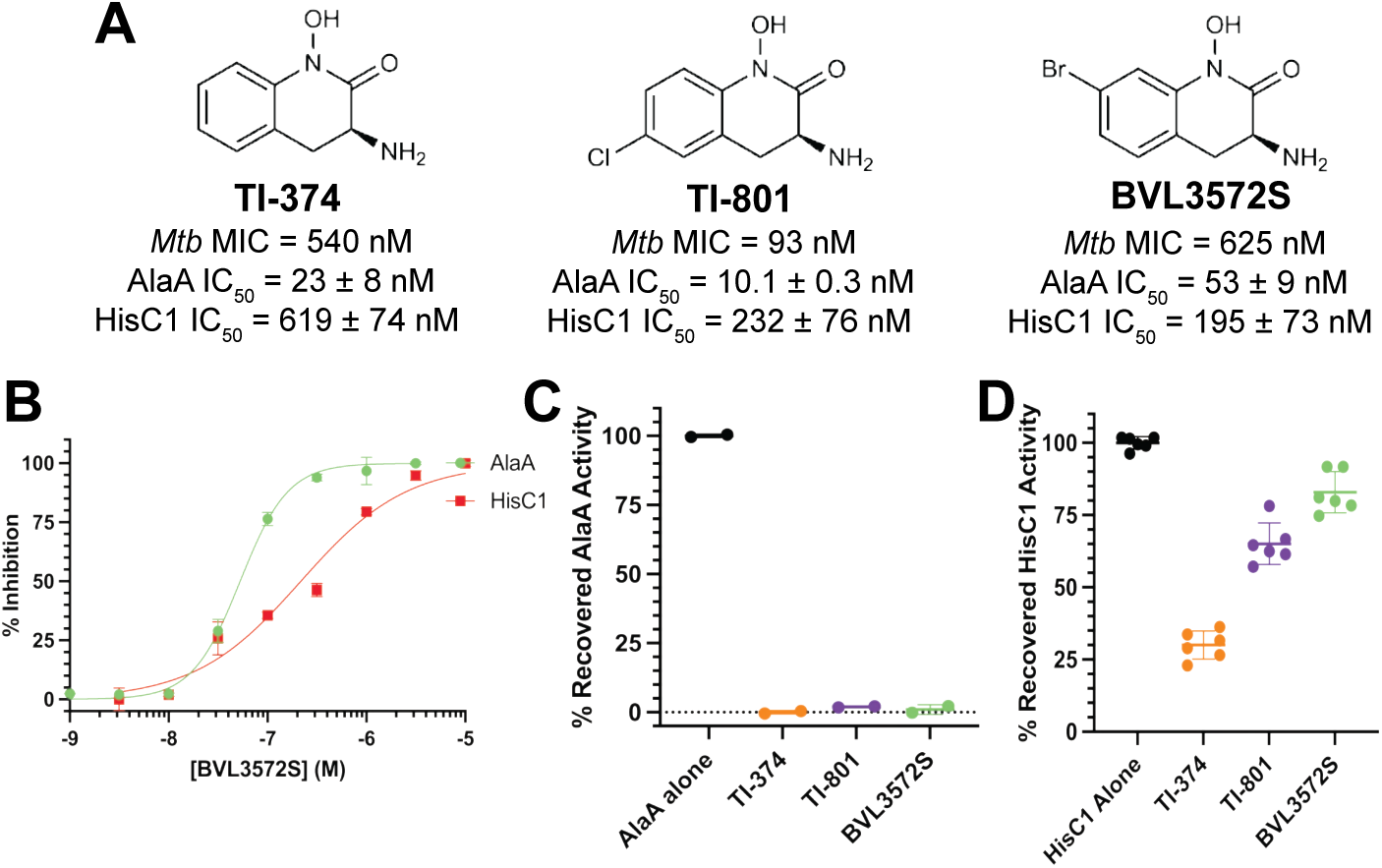
Differential reversibility against AlaA and HisC1. (**A**) Structures of analogs with *Mtb* MICs and biochemical IC_50_ values against AlaA and HisC1. (**B**) Dose-response curves for BVL3572S against both AlaA and HisC1 presented as mean ± SD, n = 3. (**C**) Recovered AlaA activity after jump-dilution for each analog compared to AlaA alone presented as mean ± SD, n = 2. (**D**) Recovered HisC1 activity after jump-dilution for each analog compared to AlaA alone presented as mean ± SD, n = 6.

For BVL3572S, interaction with HisC1 was previously characterized by Edoo et al,^32^ and our data extend the kinetic picture in two ways. First, we report quantifiable values for the molecule inhibition against both AlaA and HisC1 where these were not reported prior. Second, the full reversibility we observe at HisC1 was also observed by Edoo et al. and provides an orthogonal correlation of the short-occupancy HisC1–BVL3572S adduct reported in that work. Third, we find that BVL3572S irreversibly inactivates AlaA, a characteristic not previously reported, indicating that AlaA is the kinetically committed target across this chemotype. Finally, our data demonstrates a preliminary SAR in terms of reversible inhibition for HisC1 across the three analogs. The structural basis for the divergent adduct stability between AlaA and HisC1, such as differences in positioning of the catalytic lysine or in solvent access to the external aldimine, remains to be established and will require co-crystal structures of the inhibited enzymes.

### AlaA is indispensable in host-relevant environments despite exogenous alanine

To determine whether the alanine-dependent rescue observed *in vitro* extends to host-relevant environments, we first evaluated the effect of pharmacologic inhibition during intracellular infection. In the experiment with infected THP-1 macrophages, treatment with TI-374 (5 µM or 50 µM) or TI-801 (1 µM) supplementated with alanine did not restore intracellular growth of *Mtb* (**Figure 6A**). The same was observed in the Δ*alaA* mutant infected THP-1 macrophages, in which the knockout exhibited a pronounced defect in intracellular survival compared to WT, was not rescued by alanine supplementation (**Figure 6B**). Complementation restored bacterial growth, confirming that this phenotype is attributable to loss of AlaA. Because Mtb readily uses exogenous alanine in broth, this result is most consistent with limited delivery of supplemented alanine to intracellular bacteria, possibly reflecting uptake across host membranes or compartmentalization within the phagosome, rather than an intrinsic inability of *Mtb* to scavenge alanine. Functionally, it indicates that host-derived alanine is unlikely to bypass AlaA inhibition during infection.

**Figure 6.**
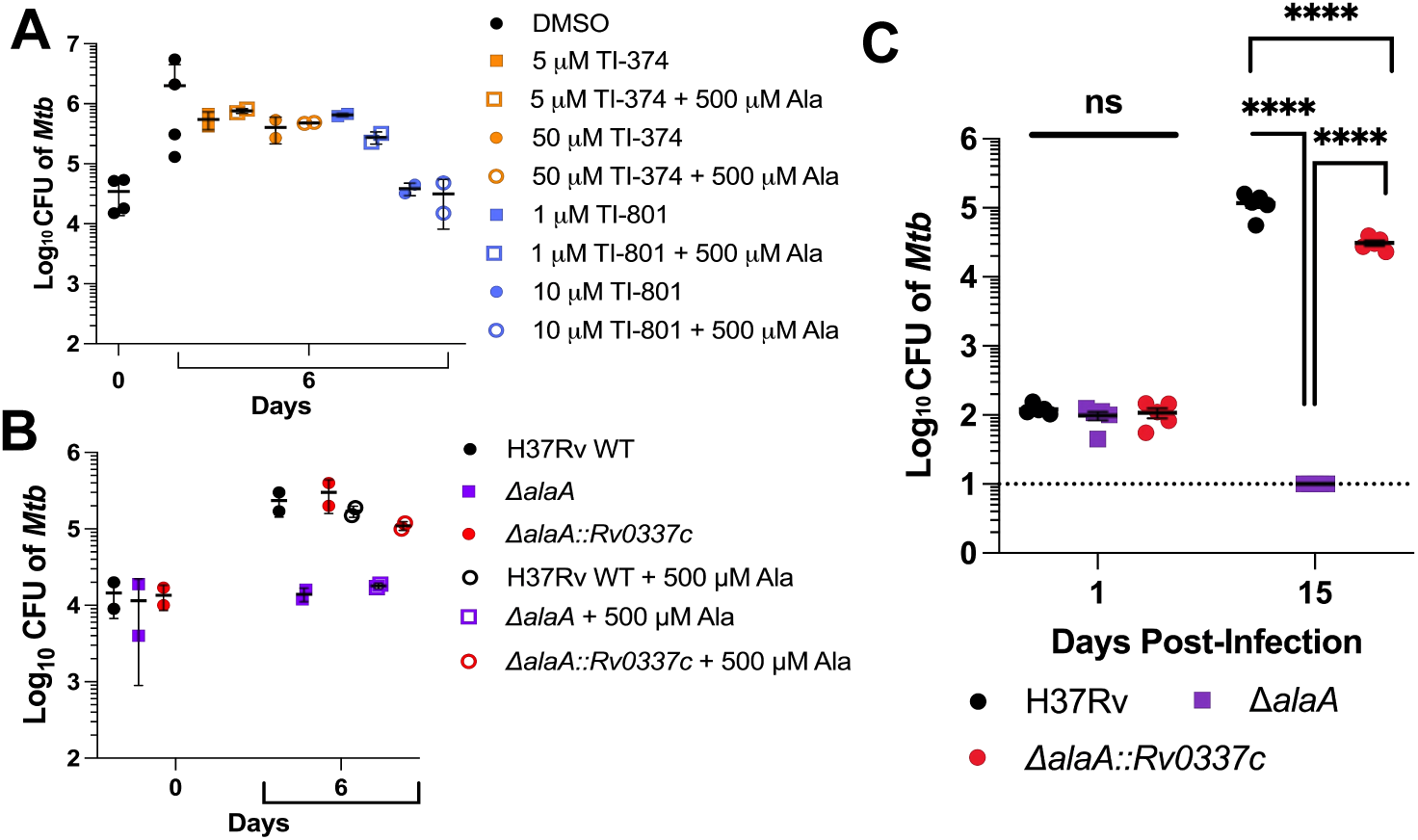
AlaA is required for growth and fitness in host-relevant environments. (**A**) Intracellular growth of Mtb in THP-1 macrophages treated with TI-374 or TI-801 at indicated concentrations, with or without alanine supplementation. Data presented as mean ± SD, n = 2 biological replicates. (**B**) Intracellular growth of WT, *ΔalaA*, and complemented (*ΔalaA::Rv0337c*) strains in THP-1 macrophages in the presence or absence of alanine. Data presented as mean ± SD, n = 2 biological replicates. (**C**) Bacterial burden in lungs of BALB/c mice infected via aerosol with WT, *ΔalaA*, or complemented strains. CFU were measured at 1- and 15-days post-infection. Data represent individual mice, with bars indicating mean ± SEM. The dotted line represents the limit of detection. Statistical analysis was performed using two-way ANOVA with Bonferroni correction for a priori pairwise comparisons of biological interest: H37Rv vs *ΔalaA*, H37Rv vs *ΔalaA::Rv0337c*, and *ΔalaA* vs. *ΔalaA::Rv0337c*. Significant differences were observed for all three pairwise comparisons at Day 15, including between WT and complemented strains (****, p<0.0001),

Our original aim was to evaluate the requirement for AlaA *in vivo* using both a genetic and pharmacologic approach. However, the TI-374 scaffold is prone to rapid Phase II metabolism leading to O-glucuronide adduct formation and clearance^33,34^ making it not yet suitable for evaluation of *in vivo* efficacy. Therefore, we utilized the genetic approach in which BALB/c mice were infected via aerosol with WT, Δ*alaA*, or complemented strains. While no significant differences in bacterial burden were observed at Day 1 post-infection, the Δ*alaA* mutant displayed marked attenuation in lung CFU at 15 days post-infection (**Figure 6C**). Notably, we only recovered a single colony per lung in two of five mice. This defect was rescued by complementation, demonstrating that AlaA is required for optimal bacterial fitness *in vivo*. Together, these results demonstrate AlaA-dependent alanine biosynthesis is critical for *Mtb* survival in host environments, where exogenous alanine is insufficient to compensate for this metabolic environment.

## Discussion

The accelerating threat of antibiotic resistance highlights the urgent need to identify novel bacterial targets and diversify the chemical space of antibacterial agents. Expanding the target landscape, particularly with compounds that act through previously unexploited mechanisms, is essential for addressing persistent and drug-resistant infections. Here, we characterize the TI-374 scaffold, previously identified from a structure-guided repurposing platform,^27^ for anti-tuberculosis activity mediated through a new mechanism of action. A central strength to this approach is that each hit is linked to a set of predicted targets, enabling focused validation efforts and efficient prioritization.

Our identification of AlaA as a pharmacologic vulnerability in *Mtb* highlights the therapeutic potential of targeting underexploited metabolic pathways, such as amino acid biosynthesis. Prior work in this field have identified other amino acid biosynthesis pathways as potential therapeutic targets^10,14^ including inhibiting biosynthesis of tryptophan,^11^ arginine,^12^ methionine,^15^ and aspartate^16^ among others. Included among these was the identification of *Mtb* AlaA, formerly misannotated at AspC,^28^ as an essential gene via transposon insertion screening.^29^ However, *Mtb* AlaA was absent from lists of essential genes identified from *in vivo* transposon screens.^30,31^ The transcriptomic response to TI-374 mirrors a canonical amino-acid-starvation program, including strong *whiB7* induction as WhiB7 directly coordinates the response to alanine starvation.^43^ This signature is consistent with AlaA engagement without, on its own, establishing the target. Given the two-replicate design, we treat these data as supportive context and base target assignment on the metabolomic, genetic, and resistance results. Although their study did not nominate AlaA as a drug target, possibly due to concerns around functional redundancy or WhiB7-mediated antagonism of aminoglycosides, our findings demonstrate that both pharmacologic and genetic inhibition of AlaA impairs *Mtb* survival in host-relevant environments, including within macrophages and the murine lung.

Whereas all prior reports probe the essentiality of *Mtb* AlaA using genetic approaches, our results are the first to bridge the data with pharmacologic validation. The data clearly support *Mtb* AlaA as the intracellular target most relevant to the antitubercular activity of TI-374 and analogs at low concentrations; however, the engagement of additional targets remains possible at higher concentrations. Indeed, the transcriptomic and metabolomic results suggest that HisC1 may also be inhibited, and the scaffold also exhibits inhibition of recombinant HisC1; however, each molecule is >20-fold more potent against *Mtb* AlaA compared to HisC1 and the scaffold exhibits a degree of reversible inhibition against HisC1. CRISPRi results suggest *hisC1* is nearly as genetically vulnerable as *alaA*,^56^ yet alanine alone was capable of suppressing TI-374 activity while histidine did not. Despite this, the transition of TI-374 from bacteriostatic at 10x MIC to bactericidal at 100x MIC is intriguing. In *Mtb* some auxotrophies are bacteriostatic due to the ability to mount an adaptive response, and the similarities of the TI-374 transcriptional response to a general starvation response is consistent with this outcome. On the other hand, some individual auxotrophs are bactericidal, such as arginine and methionine,^12,15^ as well as histidine.^22,57^ The observed shift from bacteriostatic to bactericidal activity may indicate effective engagement of HisC1 in the cell. This presents the possibility of using structure-guided design to shift the specificity of TI-374 analogs from AlaA to dual inhibition with HisC1, with a potential increase in sterilizing activity.

This study also provides evidence the scaffold engages *Mtb* AlaA through irreversible, mechanism-based inhibition, likely via formation of the covalent-PLP adduct. Moreover, by leveraging a homology model for *Mtb* AlaA, and prior knowledge of TI-374 interactions with the human KAT II,^35,58^ targeted control analogs were evaluated to probe the potential binding pose for the scaffold. This highlighted the necessity of the hydroxamic acid moiety for interactions with the Arg403 and Asn204. Additional modification placing the chlorine at the 6-position of the scaffold yielded TI-801 with significant improvement in *Mtb* potency correlated to biochemical inhibition of AlaA. The chloro-substituent highlights potential for engagement with Arg42 that may be leveraged in future design of new inhibitors. While promising in terms of improved activity, the binding pose still requires experimental structural validation to fully realize a structure-based design approach for optimization. Additionally, the scaffold is known to be a substrate for glucuronide-conjugation at the hydroxamate,^34^ making it not yet suitable for demonstrating *in vivo* activity. Prior work on TI-374 analogs as development candidates for schizophrenia suggests bioisosteric and prodrug strategies for overcoming these deficiencies.^33–35,58^ Application of these and related approaches to the design of anti-mycobacterials are being explored.

While our data demonstrate that AlaA is critical for *Mtb* growth the ultimate question is whether the bacterium can survive by acquiring alanine from the host environment. In a macrophage infection model the knockout of AlaA attenuated growth which was recovered with complement of the gene. Pharmacologically, TI-374 retained full activity even in the presence of excess exogenous alanine, suggesting that intracellular *Mtb* cannot readily access environmental alanine to overcome AlaA inhibition. Based on these results, we selected a mouse model in which the majority of bacilli in the lung reside within macrophages, providing a relevant host context for initial *in vivo* target validation. While this model does not fully replicate the architecture of the granulomatous lesions in human TB, it offers translational insight into the conditional essentiality of AlaA and supports its potential as a therapeutic target under nutrient-limited, host-relevant conditions. These experiments demonstrated a lack of survival *in vivo* using the genetic knockout strain and that alanine is not effectively scavenged from the host in the acute infection mouse model. However, further studies are warranted to evaluate essentiality within the complex microenvironment of organized granulomas, which represent a more advanced and spatially complex stage of TB pathology, where gradients of nutrients and oxygen as well as immune pressure may determine bacterial fitness requirements.

These results are complemented by the recent report of Edoo et al., who independently identified the same two PLP-dependent aminotransferases, AlaA and HisC, as targets of BVL3572S, a brominated congener of the same cyclic hydroxamic acid scaffold using genetic, biochemical, and structural approaches.^32^ Arising from independent chemical starting points and separate laboratories, the two studies converge on several points including resistance mapped to the identical A270T substitution adjacent to the catalytic lysine. Both TI-374 and BVL3572S engage HisC in addition to AlaA and retain activity against intracellular *Mtb* that is not reversed by exogenous amino acids, and both implicate a covalent PLP adduct as the basis of inhibition. This concordance substantially strengthens the assignment of AlaA as a vulnerable *Mtb* target.

The two studies also differ in emphasis and, in doing so, complement one another. Edoo et al. focused on the histidine pathway and provided a crystal structure of the BVL3572S–PLP adduct within HisC,^32^ whereas our work centers on AlaA. We define the covalent mechanism-based kinetics of AlaA inactivation and show that these compounds commit irreversibly to AlaA while engaging HisC1 only reversibly, a selectivity that extends to BVL3572S. We also establish a requirement for AlaA during infection in mice. The kinetic distinction is consistent with the partially occupied HisC adduct reported by Edoo et al. and reconciles an apparent discrepancy between the studies in that alanine supplementation alone abrogates TI-374 activity, whereas BVL3572S requires simultaneous bypass of both the alanine and histidine pathways. TI-374 is ∼27-fold more selective for AlaA over HisC1, making alanine biosynthesis the rate-limiting inhibitory action, whereas the more balanced engagement of BVL3572S necessitates relief of both pathways. The rescue requirement thus tracks the relative target engagement of each congener within a shared mechanistic framework. Finally, Edoo et al. propose that BVL3572S additionally depletes the cellular PLP pool through a futile cycle of adduct formation and release. This PLP-sequestration model remains to be tested directly; our reversibility data support the lability of the HisC1 adduct on which it partly rests but do not, on their own, distinguish cofactor sequestration from direct multi-enzyme inhibition.

In summary, this study illustrates how structure-guided drug repurposing can uncover both novel bacterial targets and tractable chemical matter. Through validation of AlaA in both intracellular and *in vivo* models, and linking its inhibition to impaired survival under host-relevant conditions, we establish AlaA as a novel vulnerability in *Mtb*, and we demonstrate that TI-374 scaffold serves as a starting point for further investigation of next-generation mechanism-based antitubercular agents. These findings broaden the therapeutic landscape for TB and underscore the potential of targeting conditionally essential metabolic enzymes in the fight against drug-resistant pathogens.

## Materials and Methods

### Bacterial Strains and Growth Conditions

*Mycobacterium tuberculosis (Mtb*) strain H37Rv (TMC 102) TMC 120, and HN878 were obtained from the Trudeau Mycobacterial Culture Collection. Strain mc^2^8251, which was grown with 24 mg/L L-pantothenate, 50 mg/L L-leucine, and 50 mg/L L-methionine, was a kind gift from Dr. William R. Jacobs, Jr. of Albert Einstein College of Medicine. Unless otherwise indicated, *Mtb* was cultured in Middlebrook 7H9 medium (BD Difco, Franklin Lakes, NJ) containing 10% v/v OADC (0.5 g/L oleic acid (Thermo Fisher Scientific, Waltham, MA); 50 g/l albumin (Fisher Scientific, Hampton, NH); 20 g/L dextrose (Fisher Scientific); 0.04 g/L catalase (Thermo Fisher Scientific) and 8 g/L sodium chloride (Fisher Scientific), 0.5% v/v glycerol (Fisher Scientific) and supplemented with 0.05% tyloxapol (Thermo Fisher Scientific). Cultures were maintained shaking at 120 rpm at 37 °C. *Escherichia coli* (*E. coli*) strains DH5⍺ (Thermo Fisher Scientific) and HST08 Stellar (TakaraBio, Kusatsu, Shiga, Japan) were used for cloning and plasmid construction and were grown in Luria-Bertani (LB) broth (Fisher Scientific) supplemented with 50 µg/mL ampicillin (Sigma) or 50 µg/mL kanamycin (Sigma).

### Drug Susceptibility Assays

An actively growing culture of *Mtb* was sub-cultured into fresh medium to optical density at 600 nm (OD_600_) 0.025 and grown at 37°C with shaking to OD_600_ 0.3–0.6. A total of 6 × 10^4^ colony forming units (CFU) of bacteria were transferred to each well of a 96-well flat bottom microtiter plate (Thermo Fisher Scientific) in 100 μL of Middlebrook 7H9 complete medium, supplemented with PF-04859989 (TI-374, Cayman Chemical, Ann Arbor, MI), starting at 10 μM and serially diluted two-fold. To prepare test compounds, 10 mM stock solutions were prepared in DMSO (Alfa Aesar, Haverhill, MA). Microtiter plates inoculated with *Mtb* were incubated standing at 37 °C for seven days. To quantify the antimicrobial effects of the test compound, 30 μL of 100 μg/mL resazurin (Acros Organics, Morris Plains, NJ) was added to each well, the plates were incubated at 37°C for an additional 24 h, after which relative fluorescent units (RFUs) were measured at excitation 540 nm/emission 590 nm with a BioTek Cytation 5 Cell Imaging Multimode Reader (Winooski, VT). Raw assay data were uploaded to the Collaborative Drug Discovery (CDD) platform, which was used to calculate Minimum Inhibitory Concentration (MIC) values and facilitate further data analysis and interpretation. The MIC was defined as the lowest concentration of a test compound resulting in more than 50% inhibition of bacterial growth, relative to DMSO negative controls.

For nutrient supplementation assays, complete Middlebrook 7H9 medium was prepared with TI-374 at 5 µM and serially diluted 2-fold. Casamino acids (Sigma) were supplemented to 2 mg/mL final concentration and alanine, histidine, serine and arginine were each supplemented to 5 mM final concentration. Glycine was supplemented to 0.59 mM, cysteine to 0.1 mM and hemin to 0.003 mM. All amino acid supplements were acquired from Sigma.

### Time-kill Assays

In a 30 mL inkwell bottle, a log-phase culture of *Mtb* was subcultured in fresh 7H9 complete medium to approximately 3 x 10^6^ CFU/mL. Compounds to be tested were added to the diluted culture at the indicated concentrations, with DMSO used as a negative inhibition control. Cultures were incubated at 37 °C with shaking. To quantify the antibacterial effects of each test compound, 100 μL aliquots of culture were removed at specified timepoints, serially diluted in Phosphate Buffered Saline (PBS) supplemented with 0.05% tyloxapol and plated on Middlebrook 7H10 solid medium (BD Difco) supplemented with 10% OADC and 0.5% glycerol to enumerate the bacteria remaining. Plates were incubated at 37 °C for 3 weeks and colonies were counted. The assay was conducted in triplicate.

### THP-1-Derived Macrophage-like Infection and Intracellular Survival Assessment

Human promonocytic THP-1 cells (ATCC, TIB-202) were maintained and differentiated in RPMI 1640 (Thermo Fisher Scientific) supplemented with 2 mM L-glutamine, 10% heat-inactivated fetal bovine serum (FBS; Thermo Fisher Scientific) and 0.05 mM 2-mercaptoethanol (Sigma, Burlington, MA; complete medium) at 37°C in a humidified with 5% CO_2_ atmosphere incubator. Cell line maintenance, passage and differentiation were as detailed previously.^59^ THP-1 cells were cultured to a density of 8 x10^5^ cells/mL, centrifuged at 500 x g for 7 min at 4°C and the cell viability was assessed using Trypan blue and counting on a hemocytometer. To differentiate THP-1 into macrophage-like cells, the cells were resuspended at a density of 2 x10^6^ cells/mL, induced with 100 nM of phorbol 12-myristate 13-acetate (PMA; Tocris Bioscience, Bristol, UK) and 100 µL was seeded per well in a 96-well tissue-culture treated plate (Corning, Tewksbury, MA). After 24 h, non-adherent cells were washed away with pre-warmed PBS. For the infection, an actively growing culture of *Mtb* was centrifuged at 3500 x g for 10 min, resuspended in cell culture medium, and used to infect cells at a multiplicity of infection (MOI) of 0.1 for 4 h (37°C, 5% CO_2_). The bacterial suspension was removed, and the cells were washed twice with 0.2 mL of pre-warmed PBS to remove extracellular bacteria. Subsequently, cells were treated with indicated concentrations of TI-374, alanine, or DMSO as the untreated control such that the solvent final concentrations did not exceed 0.1%, and incubated for up to 6 days (the medium and treatment was exchanged day 3 post-infection) at 37°C, 5% CO_2_.

Quantification of viable *Mtb* colony-forming units (CFUs) was performed at 0-, 3- and 6-days post infection. Culture media was discarded, wells were washed once with 0.2 mL pre-warmed 1 x PBS and the phagocytes were lysed by adding Triton™ X-100 to 0.1% final and incubating at 37°C for 10 minutes. The resulting lysates were serially diluted in 1 x PBS+0.05% tyloxapol and 100 µL of each dilution sample was plated onto a quadrant of the appropriate 7H10 agar plate. The plates were placed in a Ziploc® bag and incubated at 37°C with 5% CO_2_ for at least 3 weeks and colonies were counted. The experiment was performed in triplicate.

### Transcriptomics

10 mL *Mtb* H37Rv bacterial cultures were grown in 30 mL inkwell bottles at 37°C shaking at 120 rpm until an OD_600_ of 0.4 was reached, then treated with DMSO or 5 µM TI-374 in duplicate. After 4 h incubation, the cultures were harvested by centrifugation at 3500×g for 10 min, resuspended in 1 mL of Trizol reagent and transferred to 2-mL screw cap bead-beating tubes, filled with 300 μL of 0.1 mm zirconia/silica beads (BioSpec). The tubes were processed in a bead beater (BioSpec) for 45 seconds, three times with cooling on ice in between rounds for 5 minutes. Total RNA was extracted using the Direct-zol™ RNA Miniprep Kit (Zymo Research) following the manufacturer’s instructions. The resulting RNA quality and quantity were assessed by Qubit Fluorometer (Invitrogen). Sequencing library preparation was performed with the Zymo-Seq RiboFree® Total RNA Library Kit according to the manufacturer instructions.

Libraries were subsequently quantified by Qubit Fluorometer and normalized according to the Illumina Denature and Dilute Libraries Guide “Standard Normalization Method”, and finally, equimolar quantities (1.8 pM) of all libraries and 1% PhiX spike-in as control were sequenced in a 2 x 38 cycle high output run on an Illumina MiniSeq System.

FASTQ files were uploaded to Galaxy,^60^ preprocessed with fastp,^61^ and aligned to the H37Rv reference genome with Bowtie2.^62^ Reads were binned with featureCounts^63^ and differential expression was evaluated with DESeq2.^64^ The fastq files have been deposited in NCBI GEO under accession GSE267532.

### Isolation of Resistant Mutants and Whole Genome Sequencing

*Mtb* H37Rv (10^9^ CFU) was spread onto 7H10 agar plates containing 40 μM of TI-374 to isolate resistant mutants. The plates were incubated at 37°C for 4 weeks. To prepare genomic DNA, clones were grown at 37°C with shaking in complete 7H9 to reach an OD_600_ of 0.4. and genomic DNA extraction was performed using the Quick-DNA Fungal/Bacterial DNA MiniPrep Kit (Zymo Research). Libraries were prepared using the Illumina DNA Prep and sequenced using Illumina MiniSeq System 2 x 151 cycles. FASTQ files were uploaded to Galaxy^60^ and variants versus the H37Rv reference genome were determined with snippy.

### Homology Model and docking for Mtb AlaA

A high-quality atomic model was generated using established homology modeling techniques, starting from known crystal structures of closely related pyridoxal phosphate (PLP)-dependent transaminases. Based on a BLAST search from the Universal Protein Resource (UniProt)^65^ an alanine aminotransferase from *E. coli* (UniProt ID: P0A959, PDB ID: 4CVQ) was identified as a suitable template.^54^ Both *E coli* AlaA and *Mtb* AlaA (UniProt ID: P9WQ91) were aligned in UniProt and the alignment file was analyzed using ESpript^66^ and were found to share 59.16% sequence identity.

Protein sequences for both the template and target were retrieved from the UniProt. Sequence alignments were performed using the Prime STA (Structure-Based Template Alignment) method. Homology models were then constructed using the Homomultimer Model tool in Schrödinger, which builds multichain structures based on highly similar, correctly positioned templates. Specifically, chains A and B of AlaA were used as separate templates to construct the AlaA homodimer. This approach was justified by the fact that alanine aminotransferases typically form symmetric α₂ homodimers with two identical composite active sites. Structural analysis of AlaA confirms that both monomers contribute to the catalytic site, with key residues from chain B forming crucial interactions with the substrate.

Two homology models of AlaA were generated using different protocols available in the Schrödinger Maestro Suite (version 2024-2). The knowledge-based method constructs insertions and closes gaps using fragments from known structures and supports multiple model generation, making it ideal when high sequence identity is present. The energy-based method, on the other hand, applies energy minimization to construct and refine missing regions. Although it is slower and generates only a single model, it often yields greater structural accuracy. In both models, the active site was built in the same catalytically competent configuration observed in the AlaA crystal structure, featuring the covalent attachment of PLP to the conserved lysine residue.

To assess model quality, the two generated structures were aligned with the template. The energy-based model yielded an alignment score of 0.002 with an RMSD of 0.263 Å, while the knowledge-based model had an alignment score of 0.008 and an RMSD of 0.376 Å. According to Prime, lower alignment scores indicate better structural congruence; scores above 0.7 suggest poor alignment. Based on these metrics, the energy-based model was selected for further studies.

Additional refinement was performed using the Protein Preparation Wizard, which resolves structural issues, adds missing hydrogen atoms, and minimizes hydrogen and heavy atom positions to relieve close contacts. Further loop refinement was conducted on both templated and non-templated regions to explore conformational flexibility and minimize local energy. Finally, model validation was performed using a Ramachandran plot to assess backbone dihedral angles (phi/psi). The plot revealed that all residues were located within favored or allowed regions, with none in disallowed regions, indicating good stereochemical quality and overall model stability.

Molecular docking studies were carried out using the Glide module within the Schrödinger Maestro Suite (version 2024-2) to predict the binding mode of ligands within the AlaA active site. The homology model was processed using the Protein Preparation Wizard, which included assignment of bond orders, filling in missing side chains, addition of hydrogens, generation of protonation states at physiological pH using Epik, and restrained minimization using the OPLS4 force field. The docking grid was centered on the AlaA active site, defined using conserved residues identified by structural alignment with the co-crystal structure of human kynurenine aminotransferase II (hKAT II; PDB: 3UE8)^35,58^ and *E. coli* AlaA^54^ with the co-crystallized ligand serving as a geometric reference for the binding pocket. Ligands were prepared using LigPrep, generating low-energy, 3D conformations and appropriate ionization states at pH 7.4 ± 2.0. A receptor grid was generated using the Glide Receptor Grid Generation tool, centering the grid box on the PLP-binding site with default van der Waals scaling parameters (scaling factor 1, partial charge cutoff 0.25). Docking was performed using Glide in Standard Precision (SP) mode, allowing for flexible ligand sampling. The top-ranked poses were evaluated based on Glide docking scores, Glide energies, and visual inspection of key protein–ligand interactions. Docking poses were exported as PDB files and visualized using PyMOL molecular graphic program (Version 3.1.8).

### Metabolite Extraction

A 60 mL culture of *Mtb* H37Rv was grown to an OD_600_ of 0.5 in complete 7H9 media, split into 6 separate 10mL cultures and treated in triplicate with 10×MIC of TI-374 or DMSO as control for 4 hours at 37°C shaking. The cells were harvested by centrifugation (3500×g for 10 min, 4°C) and washed three times with ice-cold PBS. The pellets were resuspended in 1 mL of the extraction solvent containing acetonitrile:methanol:H_2_O (in a ratio of 2:2:1) precooled at 4°C and transferred to 2-mL screw cap bead-beating tubes, filled with 300 μL of 0.1 mm zirconia/silica beads (BioSpec). The metabolites were extracted by mechanically lysing the cell suspensions in a bead beater (BioSpec) for 45 seconds, three times with cooling on ice in between rounds for 5 minutes. Lysates were clarified by centrifugation (20,000 × g for 15 min, 4°C), the supernatant transferred to a 0.22 μm Spin-X filter tubes (Corning) and centrifuged at 5000×g for 5 min at 4°C. The sample extracts were analyzed immediately using LC–MS/MS or stored at -80°C.

### Liquid Chromatography-Mass Spectrometry

Metabolite quantitation was conducted on an Agilent 6545B LC/Q-TOF and 1290 Infinity II LC. The LC was fitted with an Agilent Poroshell 120 HILIC-Z 2.1 x 150 mm, 2.7 µm column (PN 683775-924), maintained at 15°C. Injection volume was 3 µL. The mobile phase A solvent was 20mM ammonium acetate, pH 9.3 + 5 µM medronic acid in water. Mobile phase B was pure acetonitrile. Solvents were Optima LC/MS grade from Fisher. Flow rate was 0.4 mL/min over a 24-minute run time. The gradient began at 85% acetonitrile until 1 min, decreased to 75% by 8 min, 60% by 12 min, 10% by 15 min, held to 18 min, then returned to 85% at 19.1 min and held to end of run. The MS parameters were: gas temperature 225°C, drying gas flow 9 L/min, nebulizer gas 30 psi, sheath gas temperature 375°C, capillary voltage 3000 V, nozzle voltage 500 V, fragmentor 100 V, skimmer 45 V, octopole RF peak 750 V. Purine and HP-921 were used as reference ions. Mass spectra were acquired in positive and negative modes. Initial metabolite identification was by query against a 506-compound accurate mass retention time database constructed using this method. Further tentative compound identification and quantitation was conducted by uploading raw data to MetaboAnalyst and processing the multi-modal datasets. The raw data has been deposited in Metabolomics Workbench as study ST004374.

### Construction of expression vector and expression and purification of recombinant proteins

His-AlaA was purified from *Escherichia coli* using the pET14b-MtbAlaA expression vector, which encodes the *Mtb* AlaA protein fused to a hexahistidine at the N-terminus. The *alaA* gene (Rv0337c) was amplified with DreamTaq DNA polymerase (Thermo Fisher Scientific) from *M.tb* strain H37Rv using forward primer 5’ GTGCCGCGCGGCAGCCATATGGTGGACAACGATGGCACCATTGTGG 3’ and reverse primer 5’ GCTTTGTTAGCAGCCGGATCCCTATTGCCGGTAACTGACCAGGAAG 3’. The amplified gene was purified using the Nucleospin Gel and PCR cleanup kit (TakaraBio, Machery-Nagel, Japan) and then digested with the NdeI and BamHI FastDigest restriction enzymes as per manufacturer’s instructions (Thermo Fisher Scientific). The pET14b plasmid (Sigma-Aldrich) was digested with NdeI and BamHI FastDigest restriction enzymes per manufacturer’s protocol and the *alaA* construct was subcloned into pET14b using the Snap Infusion cloning kit (Takara Bio). The assembled plasmid was transformed into HST08 Stellar competent cells (Takara Bio), as per manufacturer’s instructions, and plated onto LB plates supplemented with 100 µg/mL ampicillin (Thermo Fisher Scientific) and incubated overnight at 37°C. Colonies were selected at random and screened for the *alaA* gene using colony PCR. Positive clones were selected, propagated and plasmids were extracted and purified using the Zyppy Miniprep Kit (Zymo Research, Irvine, CA) and verified via sequencing. Expression and purification of His-AlaA was performed as previously described.^28^ Briefly, the plasmid was transformed into *E.coli* C41 (DE3) cells, and cells were grown to OD_600_ = 0.6 and expression was induced with 0.2 mM isopropyl β-d-1-thiogalactopyranoside (IPTG, MP Biomedicals) for 16 h at 37°C. Cell pellets were centrifuged at 4,670 x g at 4°C for 20 min. Supernatant was decanted and cell pellets were resuspended in lysis buffer consisting of 2 x PBS (Sigma), 10 mM imidazole (Thermo Scientific), 1 mM pyridoxal phosphate (PLP, Sigma), 5% glycerol (Fisher), 0.5 mg/mL lysozyme (Sigma), EDTA-free protease inhibitor tablets (Thermo Scientific) and 1 μg/mL DNase I (Fisher). The cells were incubated on ice for 30 minutes and then sonicated using a Model 550 Sonic Dismembranator (Fisher Scientific) at 50% power for 20 second pulses for 10 cycles, with a 30 second rest between each cycle. Lysates were clarified by centrifuging at 13,000 RPM for 30 min at 4°C. Supernatant was incubated with HisPur Superflow Nickel-NTA resin (Thermo Scientific) for 3 h at 4°C with end over end mixing. Beads were washed with lysis buffer supplemented with 226 mM NaCl and 15 mM imidazole. A second, more stringent wash step was modified and implemented, supplementing lysis buffer with 5 mM MgATP (Sigma) and 0.1 mg/mL denatured *E.coli* cell lysates and 50 mM imidazole.^67^ Protein was eluted using an imidazole step gradient of 100 mM-400 mM imidazole with 2 x PBS, 1 mM PLP and 5% glycerol. Fractions were evaluated by 12.5% SDS-PAGE analysis and AlaA containing fractions were pooled. Protein concentration was determined by Bradford assay (BioRad, Hercules, CA), using bovine serum albumin (BSA, Sigma) as the standard. Protein was stored at 4°C for short-term evaluation (up to two weeks) and aliquoted, flash frozen and stored at -80°C for extended storage.

His-HisC1 was purified from *Escherichia coli* using the pET28a(+)-MtbHisC1 expression vector, which encodes the *Mtb* HisC1 protein fused to a hexahistidine at the N-terminus, synthesized by GenScript (Piscataway, NJ) using the gene sequence from *Mtb* H37Rv Rv1600. His-HisC1 was expressed and purified as described above for His-AlaA, except that *E.coli* BL21(DE3) cells were used for expression.

### AlaA aminotransferase activity assay

AlaA aminotransferase activity was assessed using a coupled lactate dehydrogenase (LDH) assay, adapted from a previously described protocol with minor modifications.^28^ Prior to inhibition assays, each freshly purified protein batch was characterized for specific activity to ensure inter-batch consistency. Protein concentration was determined spectrophotometrically using a molar extinction coefficient of ε₂₈₀ = 60,850 M⁻¹cm⁻¹ and a molecular weight of 48,172 Da, and a fixed enzyme activity of 0.40 mU per well was maintained across all experiments, with the enzyme concentration adjusted between 0.3 and 1.0 µM depending on the specific activity of each batch.

Reactions were carried out in 384-well small-volume black plates (Corning #3542, NY, USA) in a final volume of 30 µL per well, with an estimated optical pathlength of 0.29 cm. Each reaction contained 10 mM α-ketoglutarate (Sigma), 20 mM L-alanine (Sigma), 1 mM NADH (Sigma), 1 U/mL L-lactate dehydrogenase (Roche), and 10 µM pyridoxal 5′-phosphate (PLP) in 100 mM Tris-HCl (pH 7.4). Reactions were assembled on ice and initiated by the addition of enzyme. NADH oxidation was monitored continuously at 340 nm every 5 minutes over 6 hours at 37°C using a Synergy Neo2 multimode plate reader (BioTek).

To evaluate inhibitory activity, compounds were dissolved in water to a stock concentration of 5 mM and serially diluted in water using half-log dilution steps to generate an eight-point concentration-response curve. All compounds were tested in triplicate, and the resulting data were fitted to a four-parameter logistic model (log[inhibitor] vs. normalized response, variable slope) using GraphPad Prism (version 10.6.1, GraphPad Software, San Diego, CA, USA) to determine IC_50_ values. The following controls were included in each experiment: freshly prepared AlaA, NAD⁺ control, LDH control, and no-enzyme control.

### HisC1 aminotransferase activity assay

The HisC1 aminotransferase activity assay was adapted from a previously described protocol with minor modifications.^68^ Briefly, reactions were carried out in 96-well UV-transparent microplates (Corning #3679) in a final volume of 20 µL per well. Enzymatic activity was measured using 0.125 µM HisC1 in 0.13 M triethanolamine (pH 8.5; Sigma) buffer containing 3.4 mM α-ketoglutarate. Reactions were started by adding 1.7 mM histidinol phosphate. Reactions were incubated for 45 minutes at 37°C. The reaction was stopped by adding 180 µL of 1.43 M NaOH and incubating 20 minutes at 45°C and read at 280 nm.

To evaluate inhibitor activity, compounds were dissolved in DMSO and serially diluted using half-log dilution steps to generate an eight-point concentration-response curve. All compounds were evaluated in triplicate, and the resulting data were fitted to a four-parameter logistic model (log[inhibitor] vs. normalized response, variable slope) using GraphPad Prism to determine IC_50_ values.

### Jump Dilution and Dialysis assays

To distinguish between reversible and irreversible modes of inhibition, jump-dilution and dialysis assays were performed. For the jump-dilution assay, AlaA was prepared at 100× its working concentration and pre-incubated either alone (control) or with TI-374 at 10× IC_50_ for 1 hour at room temperature. Subsequently, 1 µL from each tube was rapidly diluted 100-fold into 99 µL of assay master mix, yielding a final enzyme concentration of 1× and a residual inhibitor concentration of 0.1× IC_50_. For the dialysis assay, 1 µM AlaA was incubated with 100 µM TI-374 or assay buffer alone for 1 hour at room temperature, after which pre-dialysis activity was immediately assessed by transferring 10 µL of each sample into 20 µL of master mix (final volume 30 µL). The remaining solutions were transferred into Slide-A-Lyzer dialysis cassettes (Thermo Fisher Scientific, cat. no. 66383) and dialyzed against 1× PBS supplemented with 150 µM PLP at 4°C for 24 hours with gentle stirring, after which enzyme activity was re-assessed under identical conditions. PLP was included in the dialysis buffer to maintain protein conformational stability, as cofactor loss during dialysis can independently reduce enzyme activity and confound result interpretation. In both assays, the master mix contained 20 mM L-alanine, 10 mM α-ketoglutarate, 10 µM PLP, 1 mM NADH, and 10 U/mL LDH in 100 mM Tris-HCl (pH 7.4), and NADH oxidation was monitored continuously at 340 nm at 37°C for 3 hours as described above. The following controls were included in each experiment: freshly prepared AlaA, NAD⁺ control, LDH control, and no-enzyme control. Freshly prepared AlaA was set as the 100% activity reference, and all data were normalized accordingly and plotted using GraphPad Prism (version 10.6.1, GraphPad Software, San Diego, CA, USA).

For the HisC1 jump dilution assay, HisC1 was prepared at 100× its working concentration and pre-incubated either alone as a control, or with TI-374 or TI-801 at 10× IC_50_ for 1 hour at room temperature. Subsequently, 1 µL from each tube was rapidly diluted 100-fold into 99 µL of assay master mix, yielding a final enzyme concentration of 1× and a residual inhibitor concentration of 0.1× IC_50_. The master mix contained 3.4 mM α-ketoglutarate and 1.7 mM L-histidinol phosphate in 130 mM triethanolamine buffer (pH 8.5). Reactions were incubated at 37°C for 45 minutes, then quenched with 180 µL of 1.43 M NaOH and heated to 45°C for 20 minutes to generate the enolized imidazole acetol phosphate product, which was detected by absorbance at 280 nm as described in the HisC1 enzymatic activity section. A no-enzyme control was included in each experiment, and all data were normalized accordingly and plotted using GraphPad Prism (version 10.6.1, GraphPad Software, San Diego, CA, USA).

### Irreversible inhibition kinetics assay

To characterize the mechanism-based irreversible inhibition of AlaA, time-dependent inhibition assays were performed under the same conditions described above for IC_50_ determination. Progress curves were monitored continuously at 340 nm over 6 hours at 37°C across a range of inhibitor concentrations. Each progress curve was individually fitted to an exponential decay **equation 1**:

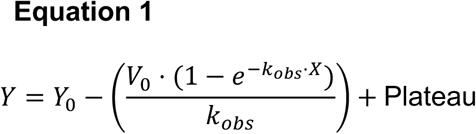

where *Y* is the observed signal at time *X*, *Y₀* is the initial product concentration (usually 0), *V₀* is the initial rate of product formation, *k*_obs_ is the first-order inactivation rate constant at each inhibitor concentration, and Plateau represents the steady-state signal value. This yielded an observed inactivation rate constant (*k*_obs_) for each inhibitor concentration tested.

The resulting *k*_obs_ values were plotted as a function of inhibitor concentration and globally fitted to a hyperbolic **equation 2** accounting for substrate competition with L-alanine^52^:

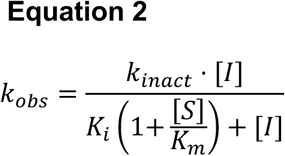

where *k*_inact_ is the maximum rate of inactivation, *K*_i_ is the apparent inhibitor concentration at half-maximal inactivation rate, [S] is the fixed substrate concentration (20 mM L-alanine), and *K*_m_ is the Michaelis constant for L-alanine. The *K*_m_ value for L-alanine was taken from the literature as 2.58 mM.^28^ The inactivation efficiency was expressed as the *k*_inact_ /*K*_i_ ratio (M⁻¹s⁻¹), which represents the second-order rate constant for enzyme inactivation and serves as the primary parameter for comparing irreversible inhibitors. All curve fitting was performed using GraphPad Prism (version 10.6.1, GraphPad Software, San Diego, CA, USA).

### Construction of strains for in vivo validation

*Mtb* was cultured in complete 7H9 medium to an OD_600_ = 0.8–1.0. Ten milliliters of the culture were transferred to a 15 mL conical tube and centrifuged at 3,500 × g for 10 minutes at room temperature. The resulting cell pellet was washed twice with 10 mL of mycobacteriophage (MP) buffer, resuspended in 500 µL of the same buffer and split equally into two new sterile 15 mL conical tubes. One tube received 1 mL of high-titer Rv0337c knockout transducing phage lysate (∼10⁹ PFU/mL; generously provided by Dr. William Jacobs Jr., Albert Einstein College of Medicine), while the control tube received 1 mL of MP buffer. Both mixtures were incubated overnight at 37°C, a non-permissive temperature for lytic phage replication. Following incubation, the cells were pelleted by centrifugation, resuspended in 300 µL of complete 7H9 medium, and plated (100 µL and the remainder) onto 7H10 agar containing 75 µg/mL hygromycin and 5 mM alanine. Plates were incubated at 37°C for 3–4 weeks to allow colony formation. Transduction frequencies were calculated by dividing the number of hygromycin-resistant colonies obtained, minus the number of spontaneous drug-resistant colonies from control cells (no phage), by the total number of viable cells. Positive *Mtb* H37Rv Δ*alaA* mutant was confirmed by PCR and the strain maintained in complete 7H9 medium supplemented with 2.5 mM alanine and 75 μg/mL hygromycin.

The pMV361-alaA plasmid was constructed using Gibson assembly. Briefly, the pMV361 vector was linearized by digestion with BamHI. The alaA coding sequence was PCR-amplified from genomic DNA of *Mtb* H37Rv strain using primers containing 20-bp overlaps homologous to the BamHI-digested ends of the pMV361 vector (Ala comp forward: 5’ CAATGGCCAAGACAATTGCGGTGGACAACGATGGCACCAT 3’ and Ala comp reverse: 5’ AGCTTCGAATTCTGCAGCTGCTATTGCCGGTAACTGACCA 3’). The linearized vector and amplified insert were subsequently assembled using Gibson assembly according to the manufacturer’s protocol. The complemented strain was generated by electroporating the integrative plasmid pMV361-*alaA* into *Mtb* H37Rv Δ*alaA* cells. Briefly, electrocompetent cells were prepared by harvesting 40 mL of a mid-log phase culture (OD_600_ = 0.8 - 1.0) and washing the pellet three times with an equal volume of sterile, room temperature 10% glycerol. After the final wash, the cell pellet was resuspended in sterile 10% glycerol at 1/10 of the original culture volume. Approximately 1 to 2 µg of plasmid DNA was mixed with 200 µL of electrocompetent cells, incubated at room temperature for 10 minutes, transferred to a 0.2 cm electroporation cuvette, and electroporated using a Gene Pulser Xcell (Bio-Rad) set to 2.5 kV, 25 µF, and 1,000 Ω. Following electroporation, 1 mL of Middlebrook 7H9 complete medium was added to each cuvette as recovery medium, and the cells were transferred to a sterile 15 mL snap-cap tube. The cultures were incubated at 37 °C for 24 hours, then plated on 7H10 agar supplemented with 75 µg/mL hygromycin and kanamycin 50 µg/mL. Plates were incubated at 37 °C until colonies became visible (3 - 4 weeks). The resulting colonies were screened by PCR for plasmid integration, and the confirmed complemented H37Rv Δ*alaA*::*Rv0337c* strain was maintained in 7H9 supplemented with 75 μg/mL hygromycin and 50 μg/mL kanamycin.

### Evaluation in a murine infection model

Six-to-eight week-old BALB/c female mice were obtained from Jackson Laboratories (Bar Harbor, ME) and housed according to the approved Trudeau Institute Institutional Animal Care and Use Committee (IACUC) protocol. *Mtb* strains (constructed as described above) were grown at 37°C to mid-exponential phase in 7H9 medium with the appropriate supplementation. Three separate groups of mice were aerosol infected with a low dose (∼100 bacilli) of *Mtb* H37Rv Δ*alaA, Mtb* H37Rv Δ*alaA* complemented strain (Δ*alaA::Rv0337c*) and *Mtb* H37Rv wild-type strain. Infection dose was verified by plating the inoculum and the whole-lung homogenates onto Middlebrook 7H11 plates (with appropriate supplementation depending on strain) at 24 h post-infection. The remaining mice were harvested 15 days post-infection. Lungs were homogenized in sterile 0.85% saline, serially diluted in saline and plated on Middlebrook 7H11 plates containing the appropriate supplementation and selection depending on strain. Plates were incubated at 37°C for 3-4 weeks and CFUs were recorded.

## Synthesis

### General Materials and Methods

Common organic solvents and reagents (>95%) were obtained from commercial suppliers and used as received. Purchased commercially available (L)-2-nitro-phenylalanine (AA Blocks LLC, Catalog # AA002BOS), (D)-2-nitro-phenylalanine (1Click Chemistry, Catalog # 4C73739), 2-bromo-4-chloro-1-nitrobenzene (A2BChem, LLC., Catalog # AB48807), and (*R*)-2-[(*tert*-Butoxycarbonyl)amino]-3-iodopropionic acid methyl ester (98%) (ChemScene, LLC, Catalog # CS-M3115) and were used without further purification. Chloroform-*d* (CDCl_3_, 99.9% D), methanol-*d_4_* (CD_3_OD, 99.9% D), water-*d_2_* (D_2_O, 99.9% D), and dimethylsulfoxide-*d_6_* (DMSO-*d*_6_, 99.9% D) were obtained from Cambridge Isotope Laboratories, Inc. and were used without further purification.

Flash Chromatography was carried out on a Smart Flash EPCLC W-Prep 2XY system using prepacked universal non-bonded silica gel (normal phase) or octadecylsilyl groups (ODS) bonded silica gel (reverse phase) cartridges by Yamazen.

^1^H NMR spectra were obtained on Bruker DRX-2 500-MHz (BBFO Z-gradient probe) or Bruker NEO-2 500-MHz (BBFO Z-gradient probe) or Bruker Avance-III 800-MHz (QCI Z-gradient cryoprobe) spectrometers. Tetramethylsilane (TMS, 0 ppm) or residual solvent (CHCl_3_/CDCl_3_, CD_3_OD/CHD_2_OD, D_2_O/HOD, and DMSO-*d*_5_/DMSO-*d*_6_) was used as internal standard. Chemical shifts (δ) are reported as parts per million (ppm) and the coupling constants (*J*) in hertz (Hz) are reported as s = singlet, d = doublet, t = triplet, q = quartet, dd = doublet of doublet, td = triplet of a doublet, dt = doublet of a triplet, m = multiplet. ^13^C NMR spectra were obtained on Bruker NEO-2 500-MHz (126 MHz ^13^C) or Bruker Avance-III 800-MHz (201 MHz ^13^C) spectrometers. Spectra were processed either using MestReNova software (v12.0).

APCI-MS (atmospheric pressure chemical ionization source, APCI) was obtained on an Advion Express Ion CMS-L Compact Mass Spectrometer either in a positive- or a negative-ion mode. HRMS for final products was recorded on an Orbitrap exploris-480 mass spectrometer purchased from Thermofisher scientific.

High performance liquid chromatography (HPLC) was obtained on an Agilent 1260 Infinity II chromatograph using an InfinityLab Poroshell 120 EC-C18 4.6 x 100 mm, 2.7 μm column. UV detection wavelength = 254 nm; flow-rate = 1.0 mL/min; gradient = 5-95% acetonitrile in water over 7 min and 3 min hold time at 95% acetonitrile in water. Both acetonitrile and water contain 0.1% v/v trifluoroacetic acid.

### Synthesis of TI-374 and analogs

The synthesis of **TI-374, TI-656, TI-772** and lactam **TI-654** was achieved through a series of steps outlined in **Scheme S1** and the synthetic procedures below. Synthesis of **TI-801**, an analogue of **TI-374** was achieved, was achieved as outlined in **Scheme S2** and the synthetic procedures below.

### Methyl (S)-2-((tert-butoxycarbonyl)amino)-3-(2-nitrophenyl)propanoate (**1a**)^69^

In a 100 mL round-bottom flask, 2-nitro-L-phenylalanine (1.0 g, 4.76 mmol) and dry methanol (20 mL) were added and stirred at 0°C. Thionyl chloride (0.42 mL, 5.71 mmol, 1.2 equiv) was added, and the mixture was heated to 45°C and stirred for 6 h. The solvents were evaporated, and the resulting methyl ester hydrochloride salt of 2-nitro phenylalanine was obtained in quantitative yield and used without further purification. The crude methyl ester hydrochloride salt of 2-nitro phenylalanine (1.20 g, 3.8 mmol) was dissolved in a 1:1 tetrahydrofuran-water mixture (20 mL) and treated with sodium bicarbonate (0.96 g, 2.4 equiv). A solution of di-*tert*-butyl dicarbonate (1.25 g, 1.2 equiv) in tetrahydrofuran (2 mL) was added dropwise at 0°C. The mixture was heated to room temperature and stirred for 12 h. The solvents were evaporated, extraction with ethyl acetate (3 x 50 mL) and water. Combined organic layers were dried with anhydrous sodium sulfate, filtered, and evaporated to obtain crude **1a** as a white-to-light brown solid (1.2 g, 3.8 mmol, 80% over 2 steps). ^1^H NMR (500 MHz, CDCl_3_) δ 7.95 (d, *J* = 8.1 Hz, 1H), 7.54 (t, *J* = 7.2 Hz, 1H), 7.45 – 7.30 (m, 2H), 5.20 (d, *J* = 8.6 Hz, 1H), 4.68 (td, *J* = 8.5, 5.8 Hz, 1H), 3.72 (s, 3H), 3.52 (dd, *J* = 13.6, 5.6 Hz, 1H), 3.24 (dd, *J* = 13.6, 8.5 Hz, 1H), 1.35 (s, 9H) ppm. APCI-MS (m/z): [M+H]^+^ calcd. for C_15_H_21_N_2_O_6_: 325.6, found: 325.1.

### Methyl (*R*)-2-((*tert*-butoxycarbonyl)amino)-3-(2-nitrophenyl)propanoate (**1b**)

Synthesized according to the same procedure as **1a** using 2-nitro-D-phenylalanine (1.0 g, 4.76 mmol), to yield **1b** as white solid (1.2 g, 3.7 mmol, 78% over two steps. ^1^H NMR (500 MHz, CDCl_3_) δ 7.95 (d, *J* = 8.1 Hz, 1H), 7.54 (t, *J* = 7.2 Hz, 1H), 7.45 – 7.30 (m, 2H), 5.20 (d, *J* = 8.6 Hz, 1H), 4.68 (td, *J* = 8.5, 5.8 Hz, 1H), 3.72 (s, 3H), 3.52 (dd, *J* = 13.6, 5.6 Hz, 1H), 3.24 (dd, *J* = 13.6, 8.5 Hz, 1H), 1.35 (s, 9H) ppm. APCI-MS (m/z): [M+H]^+^ calcd. for C_15_H_21_N_2_O_6_: 325.6, found: 325.1.

### Tert-butyl (S)-(1-hydroxy-2-oxo-1,2,3,4-tetrahydroquinolin-3-yl)carbamate (**2a**)^70^

Tin (II) chloride (948 mg, 5.0 mmol, 5.0 equiv) and sodium acetate trihydrate (1.36 g, 10.0 mmol, 10.0 equiv) were added to a solution of intermediate **1a** (324 mg, 1.0 mmol, 1.0 equiv) in 1:1 tetrahydrofuran-methanol mixture (60 mL) at 0°C. Stirred the reaction mixture, gradually warming to room temperature over 2 hours and continued stirring overnight. Concentrated the reaction mixture and the residue was extracted with ethyl acetate (2 x 20 mL) and water. Separated the organic layer, washed with water, dried with anhydrous sodium sulfate, filtered, and concentrated. Purified the crude product by silica gel chromatography to obtain the desired product **2a** as an off-white solid (120 mg, 0.43 mmol, 43%). ^1^H NMR (800 MHz, CDCl_3_) δ 8.19 (s, 1H), 7.38 – 7.31 (m, 2H), 7.22 (d, *J* = 7.5 Hz, 1H), 7.10 (td, *J* = 7.2, 1.7 Hz, 1H), 5.45 (s, 1H), 4.55 (s, 1H), 3.43 (d, *J* = 15.0 Hz, 1H), 2.90 (t, *J* = 14.5 Hz, 1H), 1.50 (s, 9H) ppm. APCI-MS (m/z): [M+H]^+^ calcd. for C_14_H_19_N_2_O_4_: 279.1, found: 279.1.

### Tert-butyl (R)-(1-hydroxy-2-oxo-1,2,3,4-tetrahydroquinolin-3-yl)carbamate (**2b**)

Synthesized using same procedure as **2a** from **1b** (324 mg, 1.0 mmol, 1.0 equiv), according to the general to isolate **2b** as off-white solid (111 mg, 0.40 mmol 40%). ^1^H NMR (800 MHz, CDCl_3_) δ 8.19 (s, 1H), 7.38 – 7.31 (m, 2H), 7.22 (d, *J* = 7.5 Hz, 1H), 7.10 (td, *J* = 7.2, 1.7 Hz, 1H), 5.45 (s, 1H), 4.55 (s, 1H), 3.43 (d, *J* = 15.0 Hz, 1H), 2.90 (t, *J* = 14.5 Hz, 1H), 1.50 (s, 9H) ppm. APCI-MS (m/z): [M+H]^+^ calcd. for C_14_H_19_N_2_O_4_: 279.1, found: 279.1.

### *Tert*-butyl (*S*)-(1-methoxy-2-oxo-1,2,3,4-tetrahydroquinolin-3-yl)carbamate (**3a**).^35^

A solution of **2a** (0.28 g, 1.0 mmol, 1 equiv) in acetone (5 mL) was treated with with potassium carbonate (1.2 g, 8.84 mmol, 8.8 equiv) for 5 minutes. Iodomethane (3.0 g, 21.1 mmol, 21 equiv) was added and refluxed the mixture on a water bath for 3 hours. Then the solvent and excess methyl iodide was evaporated with rotavap followed by addition of water to the residue to dissolve the potassium salts. The suspension was filtered and the precipitate was washed with a small amount of water, and recrystallized from ethanol to obtain the product **3a** as a white solid (100 mg, 34%). ^1^H NMR (500 MHz, CDCl_3_) δ 7.29 – 7.23 (m, 1H), 7.17 (d, *J* = 7.8 Hz, 2H), 7.05 – 7.00 (m, 1H), 5.63 (s, 1H), 4.36 – 4.27 (m, 1H), 3.88 (s, 3H), 3.38 (dd, *J* = 15.3, 6.0 Hz, 1H), 2.79 (t, *J* = 14.7 Hz, 1H), 1.43 (s, 9H). APCI-MS (m/z): [M+H]^+^ calcd. for C_15_H_21_N_2_O_4_: 293.6, found: 293.2.

### Tert-butyl (S)-(2-oxo-1,2,3,4-tetrahydroquinolin-3-yl)carbamate **4a.**^71^

Zinc powder (1.31 g, 20 mmol, 20 equiv) and ammonium chloride (1.1 g, 20 mmol, 20 equiv) were added to a solution of compound **1a** (324 mg, 1.0 mmol, 1.0 equiv) in methanol (10 mL) at room temperature and stirred the reaction mixture for 2 hours. Filtered off the zinc powder and removed the solvents in vacuo. Added 20 mL of saturated sodium bicarbonate to the resulting precipitate and extracted the aqueous phase with ethyl acetate (2 x 20 mL). Dried the organic phase over anhydrous sodium sulfate, filtered, and removed the solvent under reduced pressure. Purified the crude residue by silica gel column chromatography (5-30% ethyl acetate in hexanes containing 1% triethylamine) to obtain pure compound **4a** as off-white solid (202 mg, 0.77 mmol, 77%). ^1^H NMR (800 MHz, CDCl_3_) δ 7.52 (s, 1H), 7.22 (d, *J* = 7.1 Hz, 2H), 7.04 (t, *J* = 7.5 Hz, 1H), 6.77 (d, *J* = 7.9 Hz, 1H), 5.62 (s, 1H), 4.37 (t, *J* = 9.4 Hz, 1H), 3.58 – 3.44 (m, 1H), 2.86 (t, *J* = 14.7 Hz, 1H), 1.50 (s, 9H) ppm. APCI-MS (m/z): [M+H]^+^ calcd. for C_14_H_19_N_2_O_3_: 263.1, found: 263.1.

## Synthesis of compounds TI-374, TI-654, TI-656 and TI-772.^70^

### (S)-3-Amino-1-hydroxy-3,4-dihydroquinolin-2(1H)-one trifluoroacetate (**TI-374**).^35^

Intermediate **2a** (70 mg, 0.25 mmol, 1.0 equiv) was treated with a solution of trifluoroacetic acid (4.0 mL) and stirred the reaction mixture at room temperature overnight. Concentrated the mixture under reduced pressure, washed with diethyl ether (2 x 1 mL). The crude product was purified the crude product by reverse-phase flash column chromatography (0 – 5 % acetonitrile in water) to obtain pure **TI-374** (34 mg, 0.19 mmol 76%). ^1^H NMR (800 MHz, D_2_O) δ 7.44 – 7.34 (m, 2H), 7.30 (d, *J* = 9.2 Hz, 1H), 7.18 (t, *J* = 7.4 Hz, 1H), 4.38 (dd, *J* = 14.7, 6.6 Hz, 1H), 3.31 (dd, *J* = 15.0, 6.5 Hz, 1H), 3.23 (t, *J* = 14.8 Hz, 1H) ppm. ^13^C NMR (201 MHz, D_2_O) δ 163.9, 139.2, 130.0, 129.5, 126.5, 121.1, 115.0, 50.2, 30.0 ppm. HRMS (m/z): [M+H-C_2_F_3_O_2_]^+^ calcd. for C_9_H_11_N_2_O_2_: 179.0821, found 179.0818. APCI-MS (m/z): [M+H-C_2_F_3_O_2_]^+^ calcd. for C_9_H_11_N_2_O_2_: 179.1, found 179.1. HPLC retention time 4.514 min; HPLC purity >99.9%.

### (R)-3-Amino-1-hydroxy-3,4-dihydroquinolin-2(1H)-one trifluoroacetate (**TI-656**)

Synthesized using the same procedure as **TI-374** with intermediate **2b** (70 mg, 0.25 mmol, 1.0 equiv) using trifluoroacetic acid (4 mL) to produce **TI-656** as white solid (36 mg, 0.20 mmol, 80%). ^1^H NMR (800 MHz, D_2_O) δ 7.40 (dt, *J* = 16.9, 8.2 Hz, 2H), 7.32 (d, *J* = 7.6 Hz, 1H), 7.19 (t, *J* = 7.4 Hz, 1H), 4.40 (dd, *J* = 14.7, 6.5 Hz, 1H), 3.32 (dd, *J* = 14.9, 6.6 Hz, 1H), 3.24 (t, *J* = 14.8 Hz, 1H) ppm. ^13^C NMR (201 MHz, D_2_O) δ 163.9, 139.2, 130.0, 129.6, 126.5, 121.2, 115.1, 50.2, 30.0 ppm. HRMS (m/z): [M+H-C_2_F_3_O_2_]^+^ calcd. for C_9_H_11_N_2_O_2_: 179.0821, found: 179.0817. APCI-MS (m/z): [M+H-C_2_F_3_O_2_]^+^ calcd. for C_9_H_11_N_2_O_2_: 179.1, found 179.1. HPLC retention time 4.495 min; HPLC purity >99.9%.

### (S)-3-Amino-1-methoxy-3,4-dihydroquinolin-2(1H)-one hydrochloride (**TI-772**).^35^

Synthesized using the same procedure as **TI-374** with intermediate **3a** (73 mg, 0.25 mmol, 1.0 equiv) using hydrogen chloride in diethyl ether (4 mL) to isolate **TI-772** as an off-white solid (10 mg, 0.053 mmol, 21%). ^1^H NMR (500 MHz, D_2_O) δ 7.41 – 7.34 (m, 1H), 7.29 (t, *J* = 8.1 Hz, 2H), 7.16 (t, *J* = 7.5 Hz, 1H), 4.37 (dd, *J* = 14.5, 6.6 Hz, 1H), 3.85 (s, 3H), 3.26 (dd, *J* = 15.0, 6.7 Hz, 1H), 3.18 (t, *J* = 14.7 Hz, 1H). ^13^C NMR (126 MHz, D_2_O) δ 162.1, 135.4, 128.7, 128.5, 125.4, 120.2, 113.3, 62.8, 48.7, 28.4 ppm. HRMS (m/z): [M+H-Cl]^+^ calcd. for C_10_H_13_N_2_O_2_: 193.0977, found: 193.0975. APCI-MS (m/z): [M+H-Cl]^+^ calcd. for C_10_H_13_N_2_O_2_: 193.4, found: 193.1.

HPLC retention time 5.129 min; HPLC purity 99.4%.

### (S)-3-Amino-3,4-dihydroquinolin-2(1H)-one hydrochloride (**TI-654**).^72^

Synthesized using the same procedure as **TI-374** with intermediate **4a** (65 mg, 0.25 mmol, 1.0 equiv) using hydrogen chloride in diethyl ether (4 mL), to isolate **TI-654** as an off-white solid (34 mg, 0.22 mmol, 85%). ^1^H NMR (800 MHz, CD_3_OD) δ 7.26 – 7.16 (m, 2H), 7.02 (t, *J* = 6.8 Hz, 1H), 6.90 (d, *J* = 9.0 Hz, 1H), 3.91 (dd, *J* = 14.4, 6.5 Hz, 1H), 3.19 (dd, *J* = 15.0, 6.5 Hz, 1H), 3.02 (t, *J* = 14.7 Hz, 1H) ppm. ^13^C NMR (201 MHz, CD_3_OD) δ 170.6, 138.3, 129.4, 129.2, 124.5, 122.9, 116.7, 50.3, 32.6 ppm. HRMS (m/z): [M+H-Cl]^+^ calcd. for C_9_H_11_N_2_O: 163.0871, found: 163.0869. APCI-MS (m/z): [M+H-Cl]^+^ calcd. for C_9_H_11_N_2_O: 163.0, found: 163.1. HPLC retention time 4.421 min; HPLC purity 97.7%.

### Methyl (S)-2-((tert-butoxycarbonyl)amino)-3-(5-chloro-2-nitrophenyl)propanoate **7.**^73^

Zinc powder (127 mg, 1.94 mmol, 1.1 equiv) was added to a 10 mL round bottom flask and dried under vacuum at 80°C for 30 minutes. Suspended dry zinc powder in *N*,*N*-dimethylformamide (3 mL) and added iodine (43 mg, 0.17 mmol, 0.05 equiv). The resulting deep-red solution was stirred until decolorization, followed by the addition of methyl (*R*)-2-((tert-butoxycarbonyl)amino)-3-iodopropanoate **6** (579 mg, 1.76 mmol, 1 equiv) and stirred for 30 minutes. In a separate 10 mL round bottom flask, mixture palladium (II) acetate (4 mg, 0.017 mmol, 0.01 equiv) and 2-dicyclohexylphosphino-2’,4’,6’-triisopropylbiphenyl (X-Phos, 25 mg, 0.053 mmol, 0.03 equiv) in *N*,*N*-dimethylformamide (3 mL), stirred for 5 minutes, and added 2-bromo-4-chloro-1-nitrobenzene **5** (416 mg, 1.76 mmol, 1 equiv). The zincate solution was added and stirred at room temperature for 12 h. The reaction mixture was diluted with ethyl acetate and quenched with ice-cold water. The organic layer was separated, dried with anhydrous sodium sulfate, filtered, and concentrated in a rotary evaporator. The crude product was purified by silica gel chromatography (5 – 25% ethyl acetate in hexanes) to obtain intermediate **7** as a pale brown solid (210 mg, 0.58 mmol, 33% yield). ^1^H NMR (500 MHz, CDCl_3_) δ 7.94 (d, *J* = 8.6 Hz, 1H), 7.39 – 7.35 (m, 2H), 5.19 (d, *J* = 8.2 Hz, 1H), 4.68 (s, 1H), 3.75 (s, 3H), 3.58 – 3.53 (m, 1H), 3.16 – 3.08 (m, 1H), 1.36 (s, 9H) ppm. APCI-MS (m/z): [M+Cl]^+^ calcd. for C_15_H_19_Cl_2_N_2_O_6_: 393.0, found: 393.0.

### tert-Butyl (S)-(1-((tert-butoxycarbonyl)oxy)-6-chloro-2-oxo-1,2,3,4-tetrahydroquinolin-3-yl)carbamate (**8**).^70^

Added tin (II) chloride (948 mg, 5.0 mmol, 5.0 equiv) and sodium acetate trihydrate (1.36 g, 10.0 mmol, 10.0 equiv) to a solution of intermediate **7** (358 mg, 1.0 mmol, 1.0 equiv) in 1:1 tetrahydrofuran-methanol mixture (60 mL) at 0°C. Stirred the reaction mixture, gradually warming to room temperature over 6 hours. Added triethylamine (1 g, 1.4 mL, 10.0 mmol, 10.0 equiv) and di-*tert*-butyl dicarbonate (655 mg, 3.0 mmol, 3.0 equiv) to the mixture and stirred overnight at room temperature. The reaction mixture was extracted with ethyl acetate (2 x 50 mL) and water. Separated the organic layer, washed with water, dried with anhydrous sodium sulfate, filtered, and concentrated. Purified the crude product by flash chromatography on silica gel, eluting with 0-10% EtOAc/hexane. Obtained the desired product **8** (243 mg, 0.59 mmol, 59 %) as a gummy solid, which was immediately used for the next step. ^1^H NMR (500 MHz, CDCl_3_) δ 7.20 – 7.26 (m, 3H), 5.53 (s, 1H), 4.52 - 4.47 (m, 1H), 3.40 – 3.32 (m, 1H), 2.96 - 2.90 (m, 1H), 1.54 (s, 9H), 1.45 (s, 9H). APCI-MS (m/z): [M+H]^+^ calcd. for C_19_H_26_ClN_2_O_6_: 413.1, found: 413.7.

### (*S*)-3-Amino-6-chloro-1-hydroxy-3,4-dihydroquinolin-2(1*H*)-one hydrochloride (**TI-801**)^70^

Synthesized by *N*- and *O*-Boc deprotection of Intermediate **8** (103 mg, 0.25 mmol) using hydrogen chloride in diethyl ether (4 mL) to obtain **TI-801** as white solid (29 mg, 0.12 mmol, 47%). ^1^H NMR (500 MHz, D_2_O) δ 7.48 – 7.20 (m, 3H), 4.42 (dd, *J* = 14.3, 6.8 Hz, 1H), 3.37 – 3.10 (m, 2H) ppm. ^13^C NMR (126 MHz, D_2_O) δ 162.3, 136.6, 129.4, 128.3, 127.8, 121.4, 115.0, 48.4, 28.2 ppm. HRMS (m/z): [M+H-Cl]^+^ calcd. for C_9_H_10_ClN_2_O_2_: 213.0431, found: 213.0426. APCI-MS (m/z): [M+H-Cl]^+^ calcd. for C_9_H_10_ClN_2_O_2_: 213.0, found: 213.4. HPLC retention time 5.102 min; HPLC purity >99.9%.

## Supporting information

Compound characterization data

## Acknowledgements

The authors gratefully acknowledge funding support from the Potts Memorial Foundation. Additional funding was provided by the Trudeau Institute. We thank Dr. William R. Jacobs Jr. for providing the knockout phage used to create the *alaA* deletion strain and *Mtb* strain mc^2^8251. We also acknowledge support from the Purdue College of Pharmacy including a gift from Chip and Jane Rutledge to the College of Pharmacy (D.P.F.). Research results were also obtained with support from the National Institute of Allergy and Infectious Diseases award numbers R01AI175024 (D.P.F.) and R01AI195815 (M.C. and D.P.F.).

## Supporting Information

Compound characterization NMR spectra, mass spectra and HPLC traces are all provided in the supporting information file.

---

**Figure S1.**
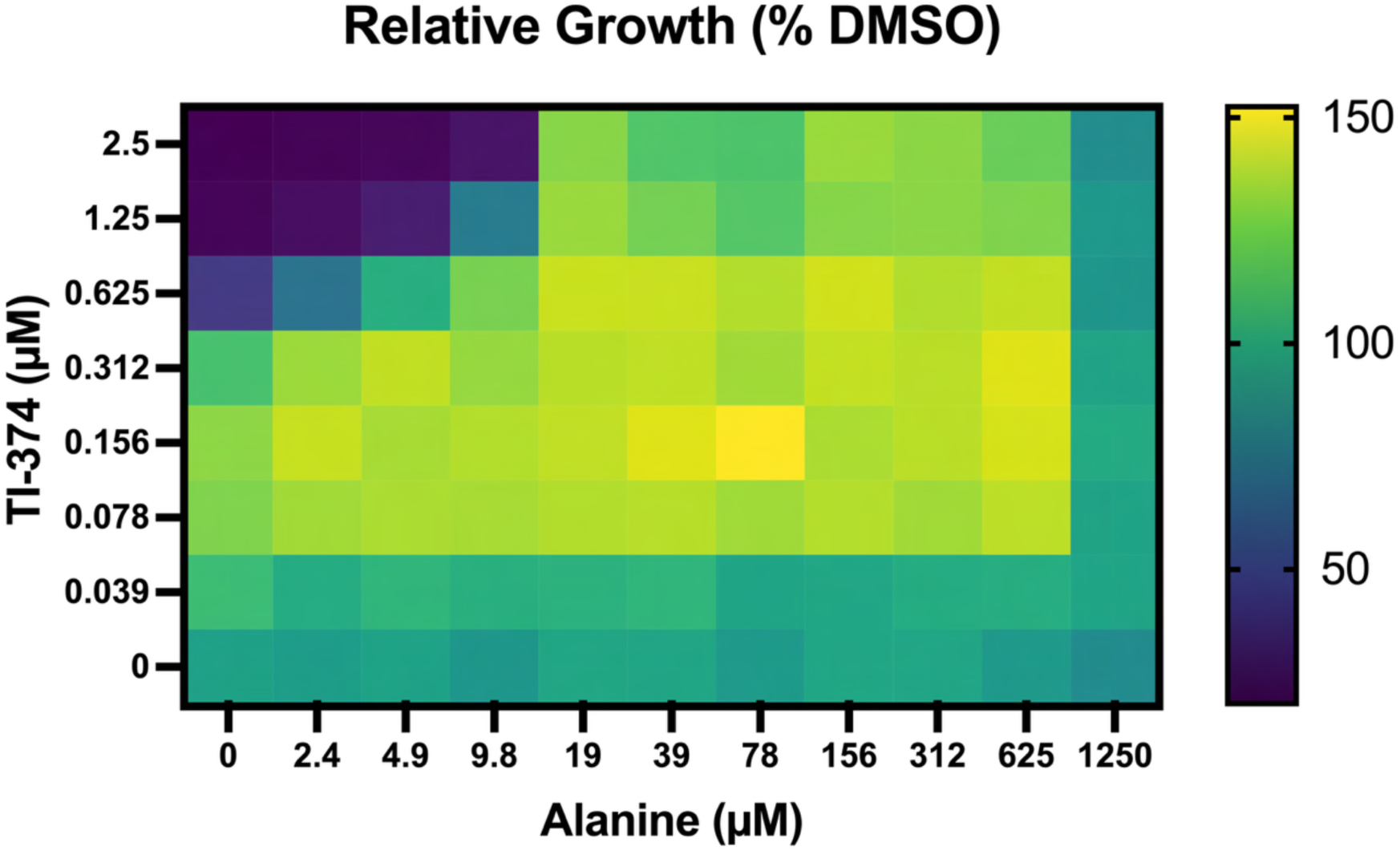
Checkerboard assay for Mtb growth with TI-374 combined alanine supplementation.

**Figure S2.**
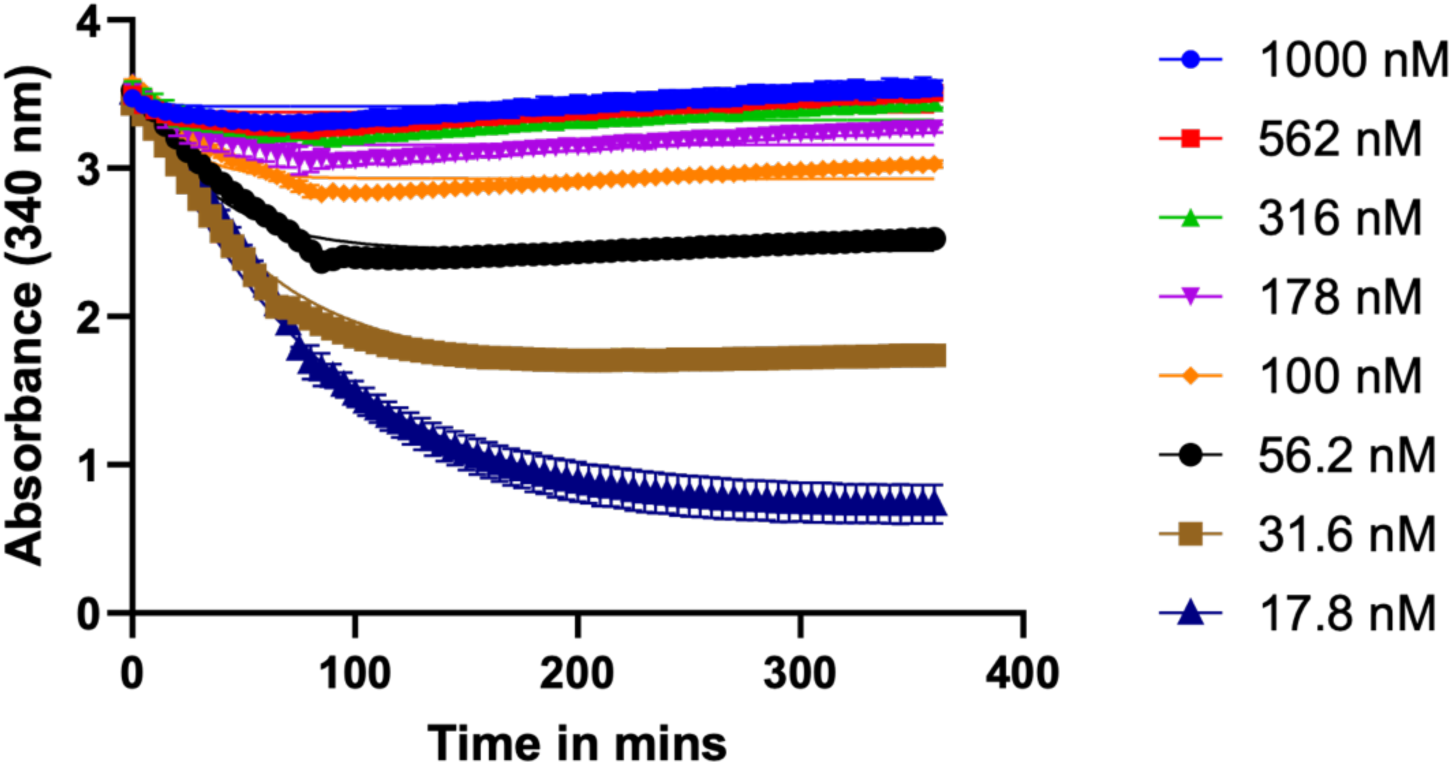
Progress curves for *Mtb* AlaA kinetic inhibition assay with TI-374.

**Figure S3.**
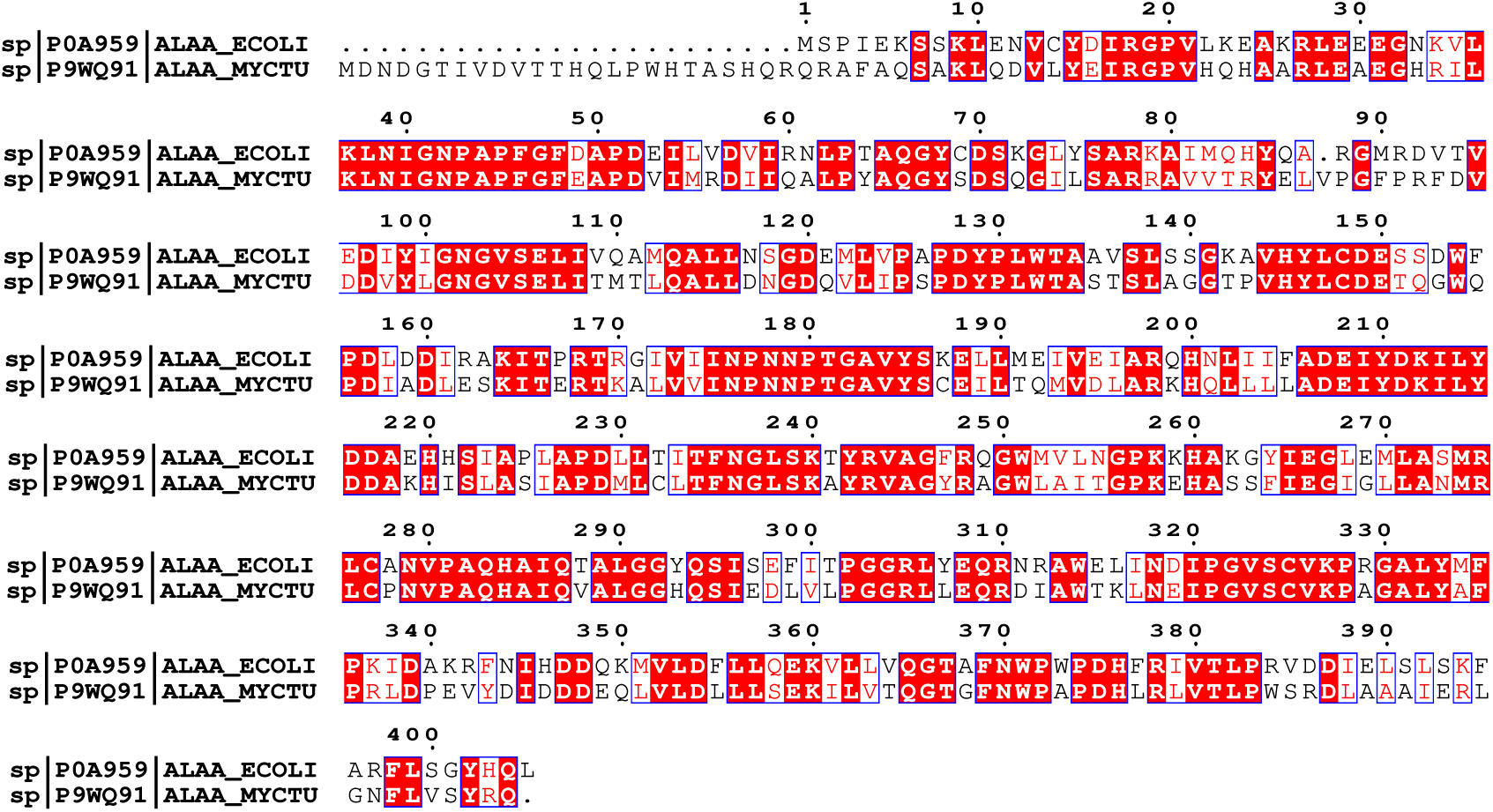
Squence alignment for *E. coli* and *Mtb* AlaA genes. *E. coli* AlaA (UniProt ID P0A959) and *Mtb* AlaA (UniProt ID P9WQ91) were aligned in Uniprot and image created using ESpript. Black font = non-similar amino acids. Red font in blue box = similar amino acids. Red highlight with white font = identical amino acids.

**Figure S4.**
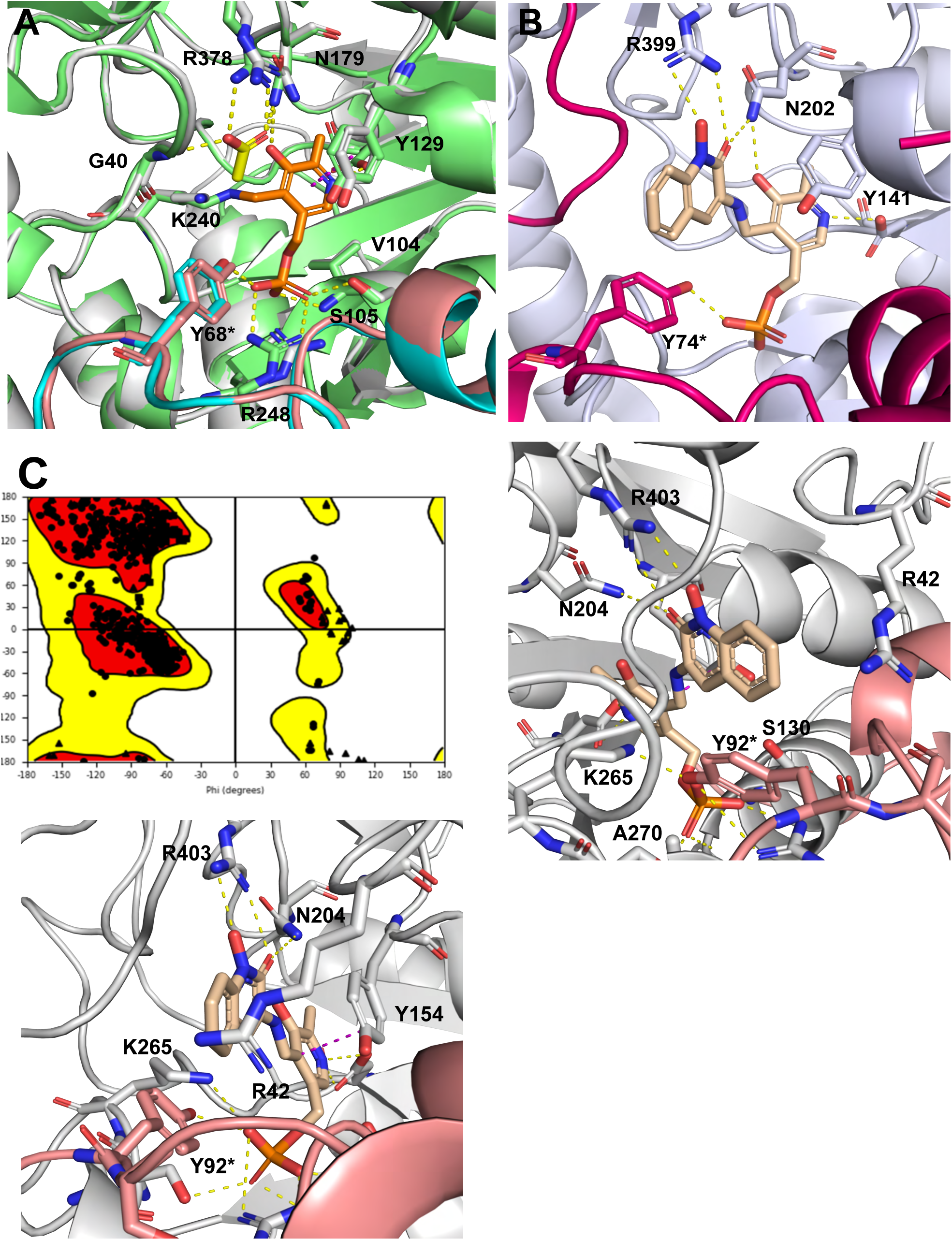
Comparison of *Mtb* AlaA homology model with previously reported structures and analysis of proposed TI-374 interactions. (**A**) *Ec*AlaA (PDB: 4CVQ, mint and cyan cartoons) with acetate (yellow sticks) and PLP (orange sticks) bound overlaid with *Mtb* AlaA homology model (gray and salmon cartoon). Identical residues in the active sites shown as sticks and labeled according to the *Ec*AlaA numbers with * indicating chain B. Hydrogen-bonds to ligand in *Ec*AlaA depicted with yellow-dashes. (**B**) Human KAT II (lavender and magenta cartoon) bound to TI-374-PLP adduct (gold sticks) (PDB: 3UE8). (**C**) Ramachandran plot for *Mtb* AlaA homology model. (**D** and **E**) *Mtb* AlaA homology model (gray and salmon cartoons) docked with TI-374-PLP adduct (gold sticks). Residues in active site shown as sticks and labeled, with * indicating Chain B, and hydrogen-bonds depicted by yellow dashed lines.

**Figure S5.**
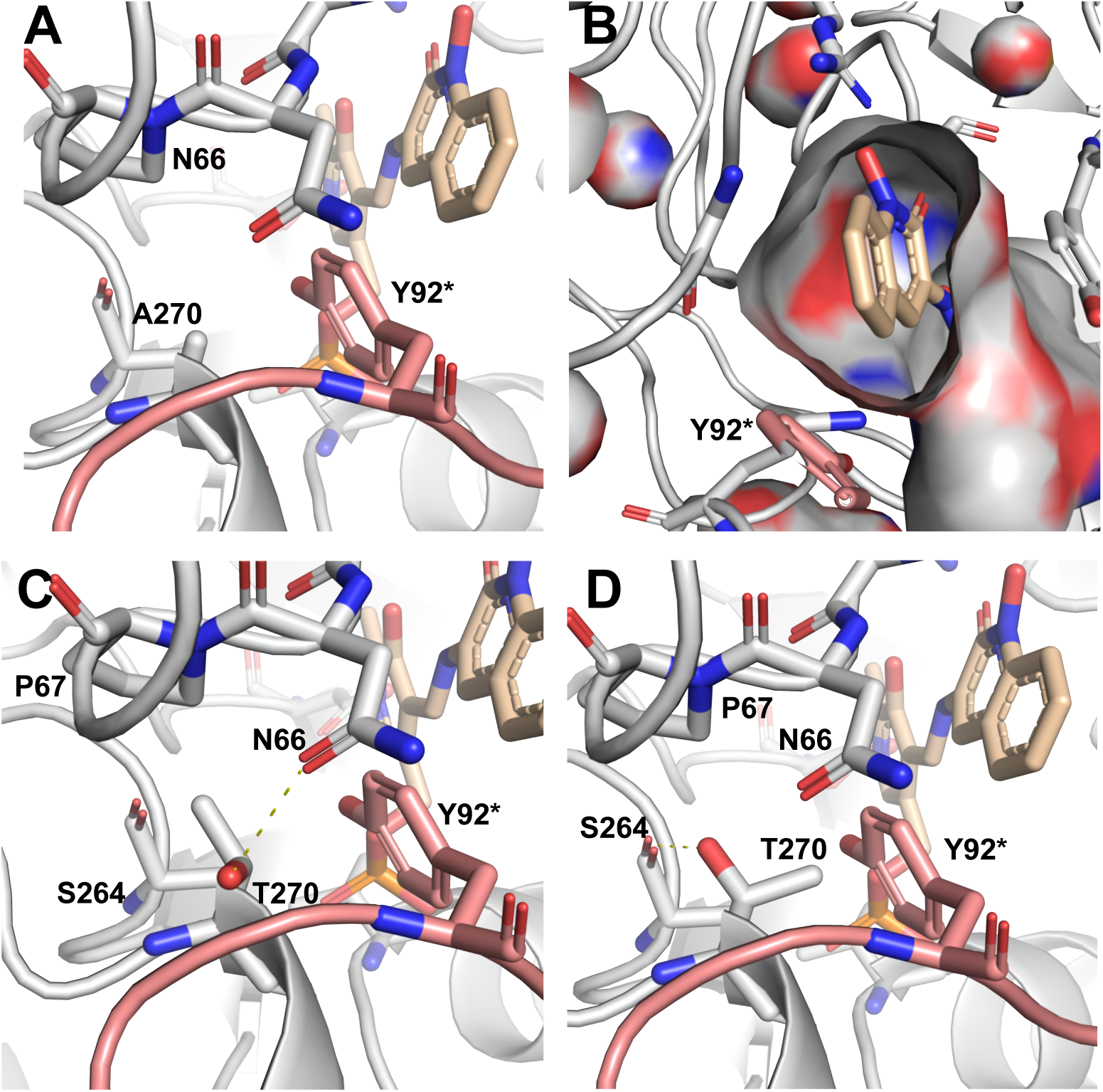
Analysis of A270T mutation on inhibition of *Mtb* AlaA (gray cartoon) by TI-374-PLP-adduct (gold sticks). Residues shown as sticks and labeled, with * indicating chain B residue. (**A**) Ala270 location shown behind Tyr92*. (**B**) Binding site surface representation with TI-374-PLP-adduct. (**C** and **D**) Mutated T270 residue in each predicted rotamer.

**Figure S6.**
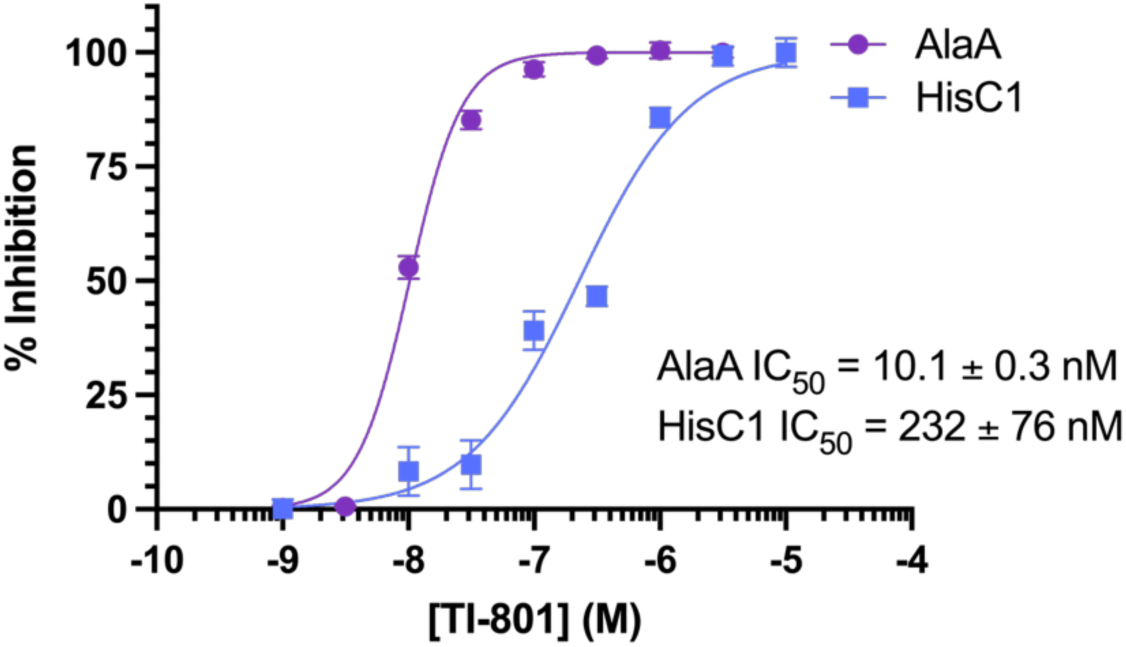
Dose response curves for TI-801 against AlaA and HisC.

**Figure S7.**
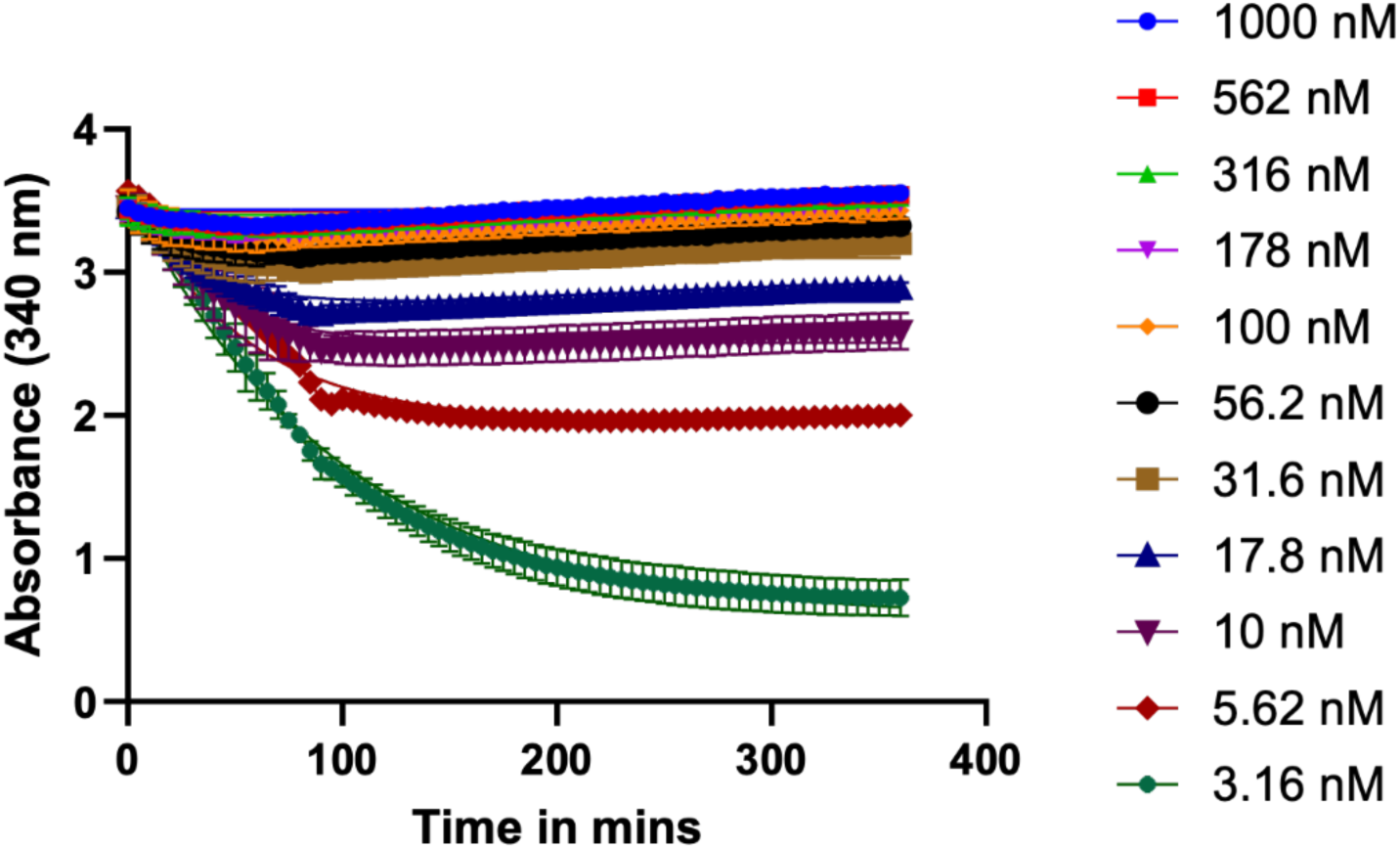
Progress curves for *Mtb* AlaA kinetic inhibition assay with TI-801.

**Table S1.**
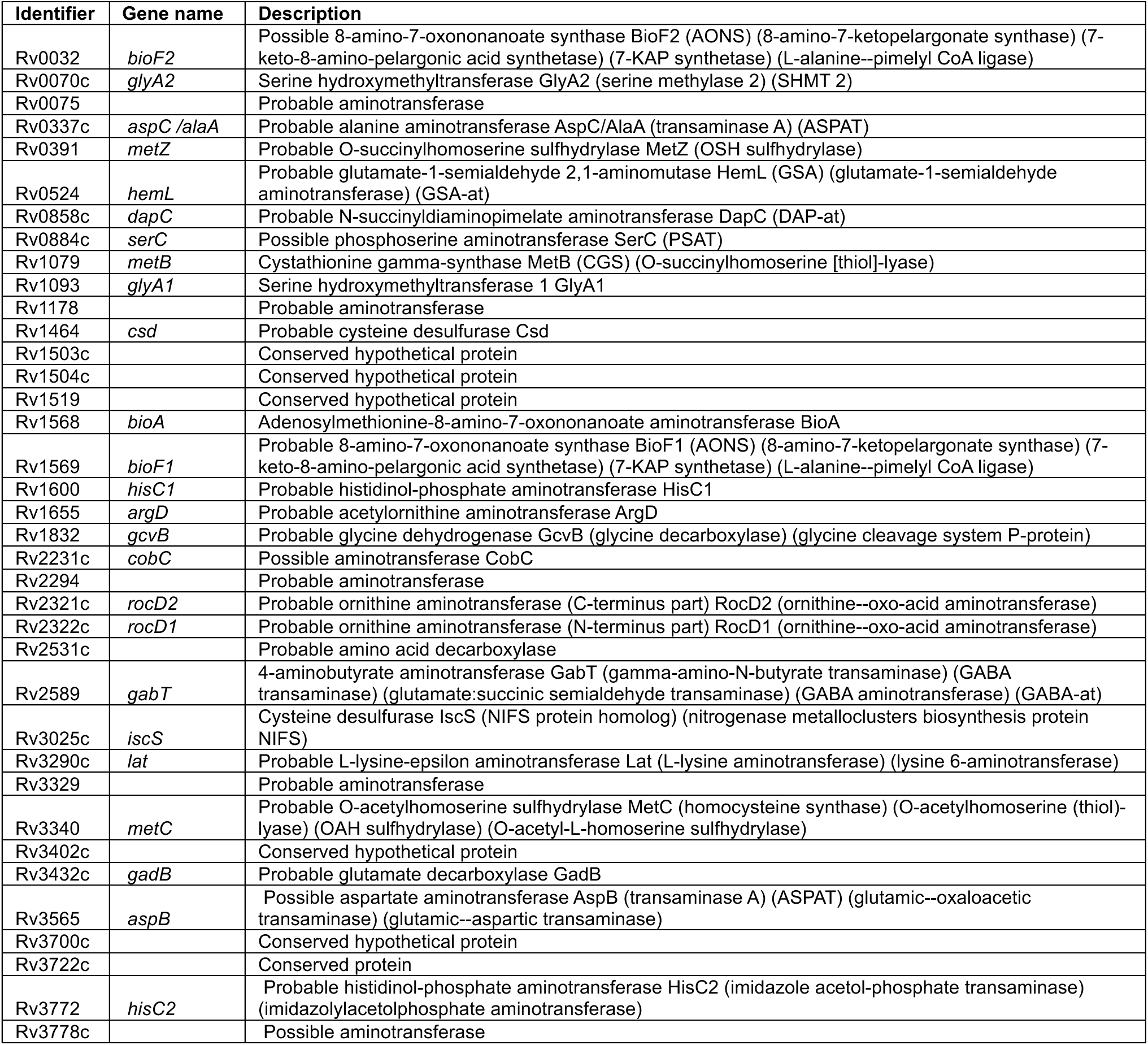
Structural domains belonging to the superfamily of PLP-dependent transferases.

**Table S2.** MIC for TI-374 against *Mtb* H37Rv supplemented with metals^a^.

| **Table S2. MIC for TI-374 against *Mtb* H37Rv supplemented with metals<sup>a</sup>** |  |  |  |  |  |  |
| --- | --- | --- | --- | --- | --- | --- |
| Treatment | TI-374 alone | + FeSO<sub>4</sub> | + ZnSO<sub>4</sub> | + MgSO<sub>4</sub> | + CaSO<sub>4</sub> | + CuSO<sub>4</sub> |
| MIC ($\mu\text{M}$) | 0.62 | 0.62 | 0.62 | 0.62 | 0.62 | 0.62 |

<sup>a</sup> TI-374 was tested in dose response with supplementation of each metal-sulfate complex. FeSO<sub>4</sub> supplemented at 50 $\mu\text{M}$ , others at 100 $\mu\text{M}$ .

**Scheme S1.**
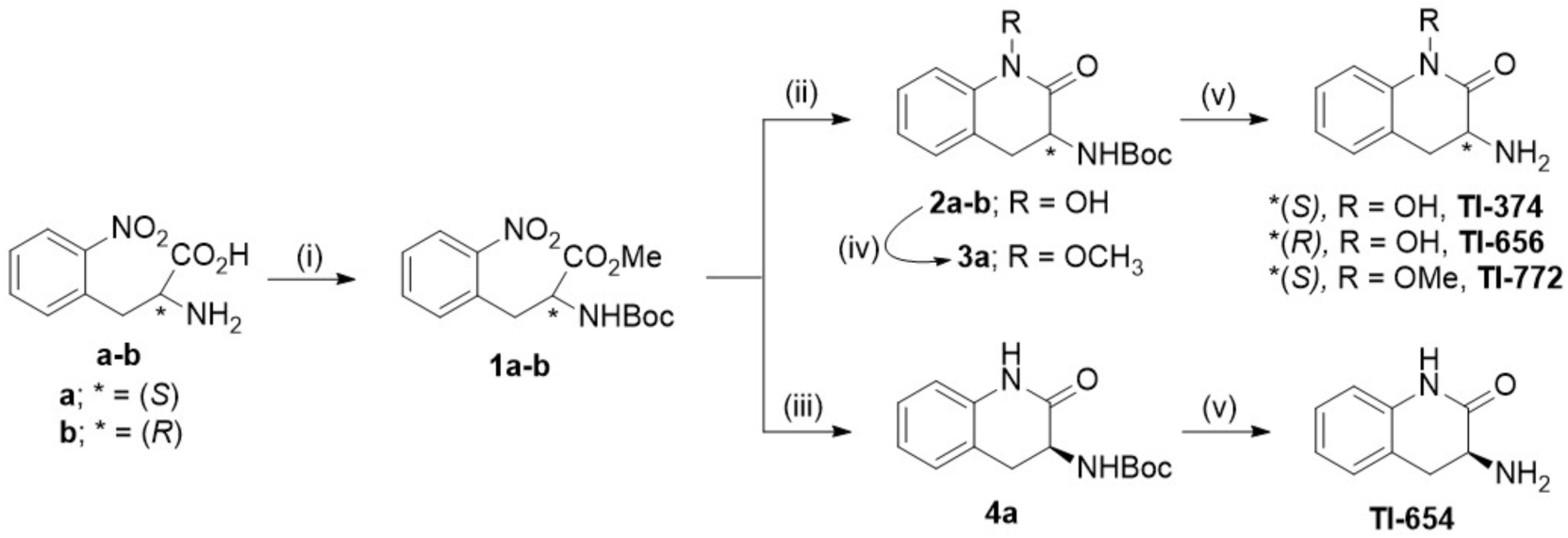
Synthesis of TI-374, TI-656, TI-772 and TI-654. Reagents and conditions: (i) SOCl_2_ (1.2 equiv), MeOH, 45°C, 6 h, and (Boc)_2_O (2 equiv), NaHCO_3_ (2 equiv), 1:1 THF-H_2_O, r.t., 12 h; (ii) SnCl_2_ (5 equiv), NaOAc•3H_2_O (10 equiv), 1:1 THF-MeOH, r.t., 12 h; (iii) Zn (20 equiv), NH_4_Cl (20 equiv), THF-MeOH, r.t., 2 h; (iv) CH_3_I ( 21 equiv), K_2_CO_3_ (8.8 equiv), acetone, 70°C, 3 h; (v) 2M HCl in diethyl ether, r.t., 12 h.

**Scheme S2.**
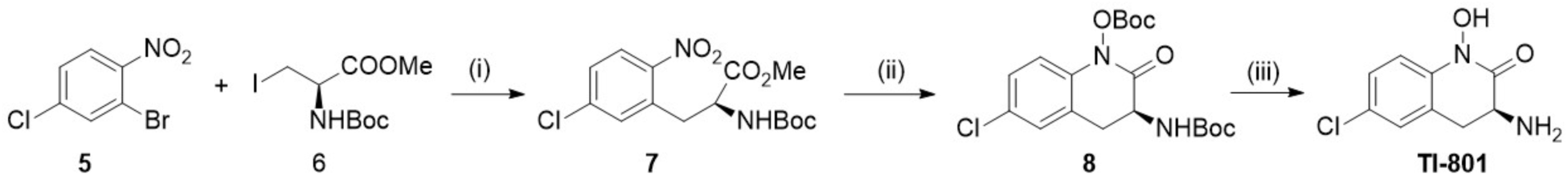
Synthesis of TI-801. Reagents and conditions: (i) Zn (1.1 equiv), I_2_ (5 mol%), Pd (OAc)_2_ (1mol %), Xphos (3 mol%) DMF, r.t., 12 h; (ii) SnCl_2_ (5 equiv), NaOAc•3H_2_O (10 equiv), 1:1 THF-MeOH, r.t., 6 h then Et_3_N (10.0 equiv), Boc_2_O (3 equiv), r.t., 12 h (iii) HCl (2M) in diethyl ether, r.t., 12 h.

