## Supplementary material for "Induced alanine auxotrophy as a therapeutic strategy against *Mycobacterium tuberculosis*": Compound characterization data

**Supporting Information File - 1**

$^{1}H$ NMR, $^{13}C$ NMR, APCI-MS and HPLC data for final compounds **TI-374**, **TI-656**, **TI-772, TI-654 and TI-801**

### $^1H$ and $^{13}C$ NMR spectra of compound – TI-374

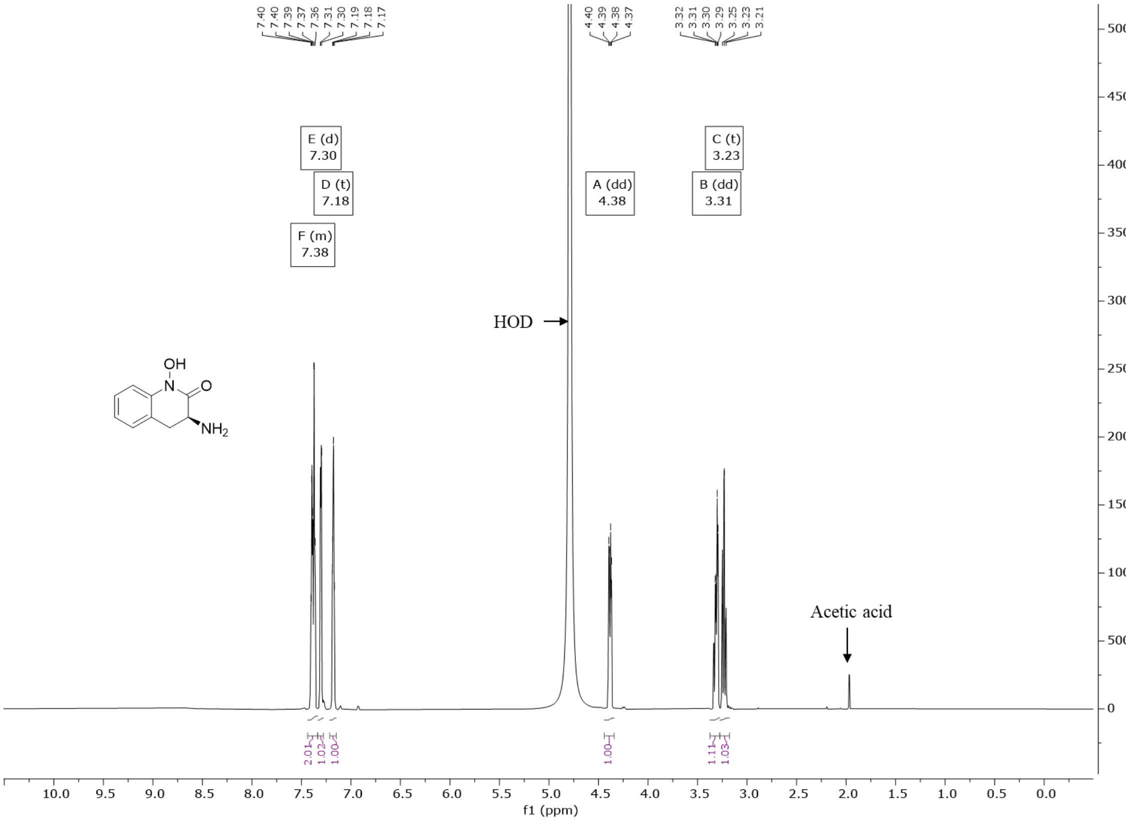


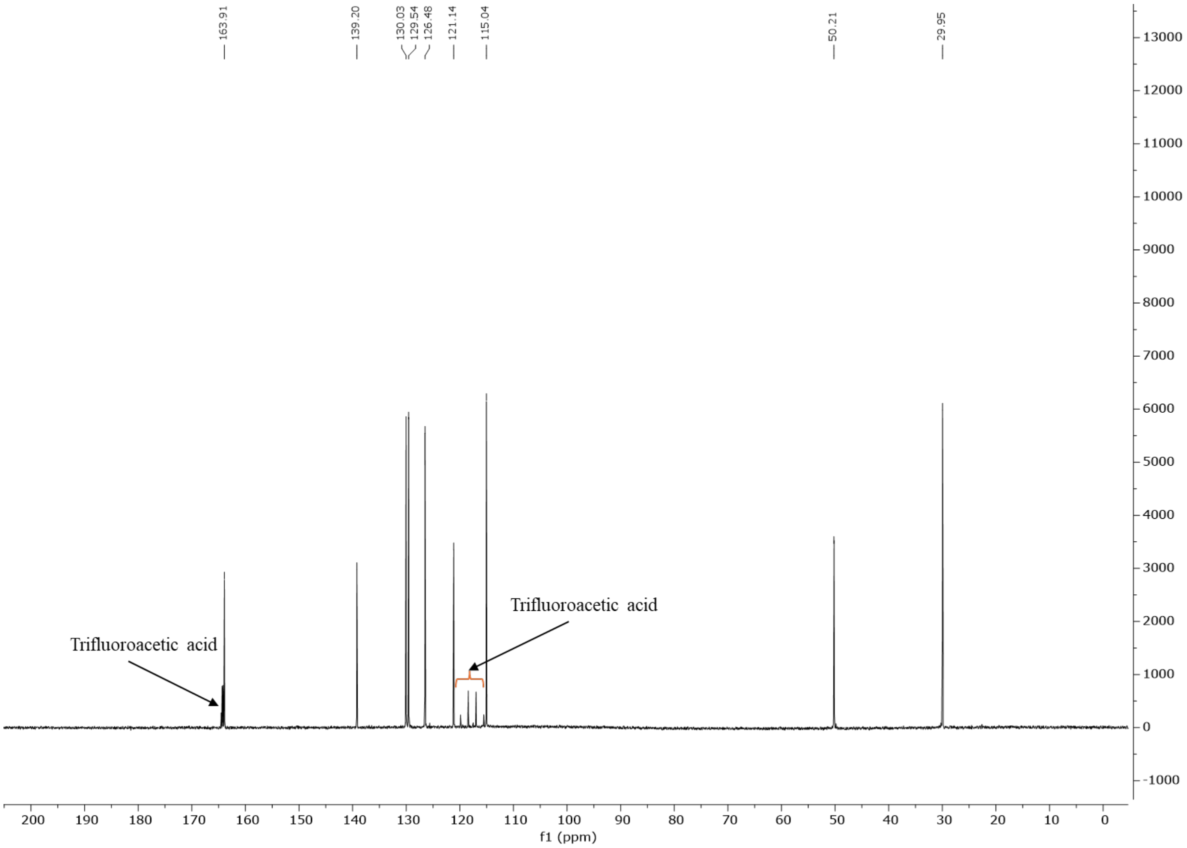

**ESI-MS spectrum and HPLC profile of compound – TI-374**

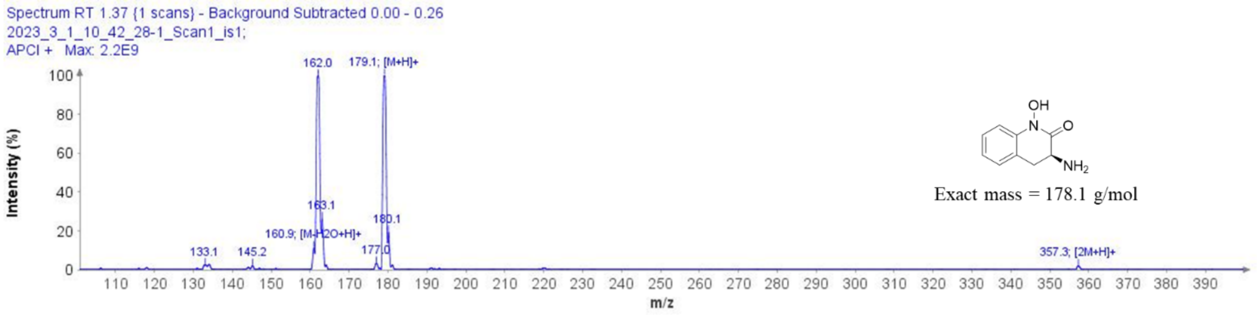


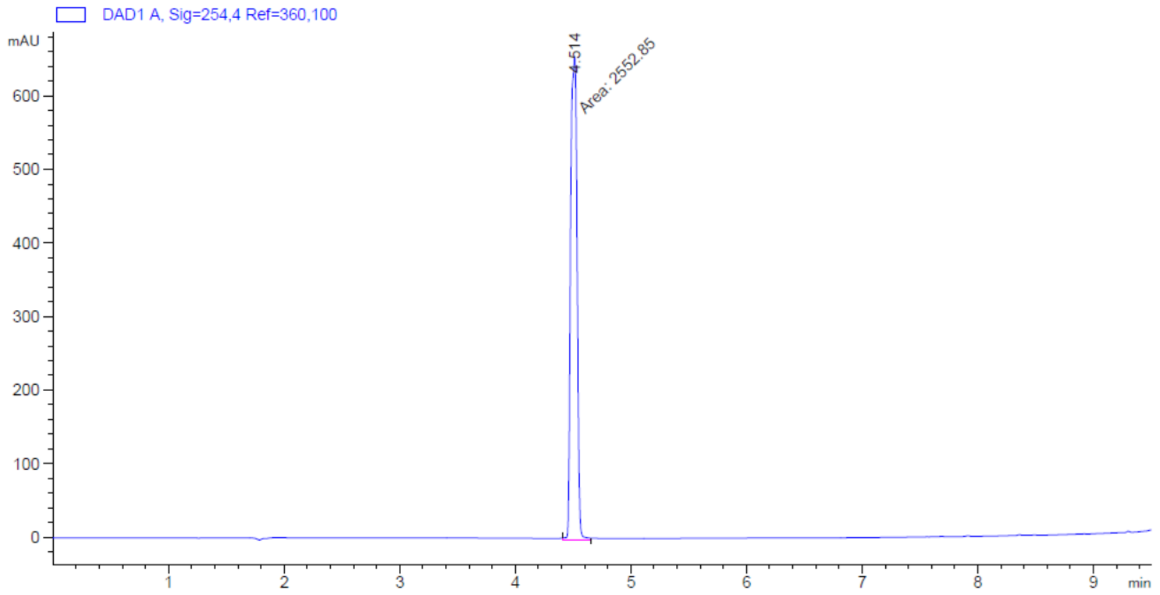


```
Signal 1: DAD1 A, Sig=254,4 Ref=360,100

Peak RetTime Type  Width     Area       Height     Area
  #   [min]        [min]   [mAU*s]      [mAU]       %
----|-------|----|-------|----------|----------|--------|
   1   4.514 MM    0.0647 2552.85156  657.86725 100.0000

Totals :                  2552.85156  657.86725
```

### $^{1}H$ and $^{13}C$ NMR spectra of compound – TI-656

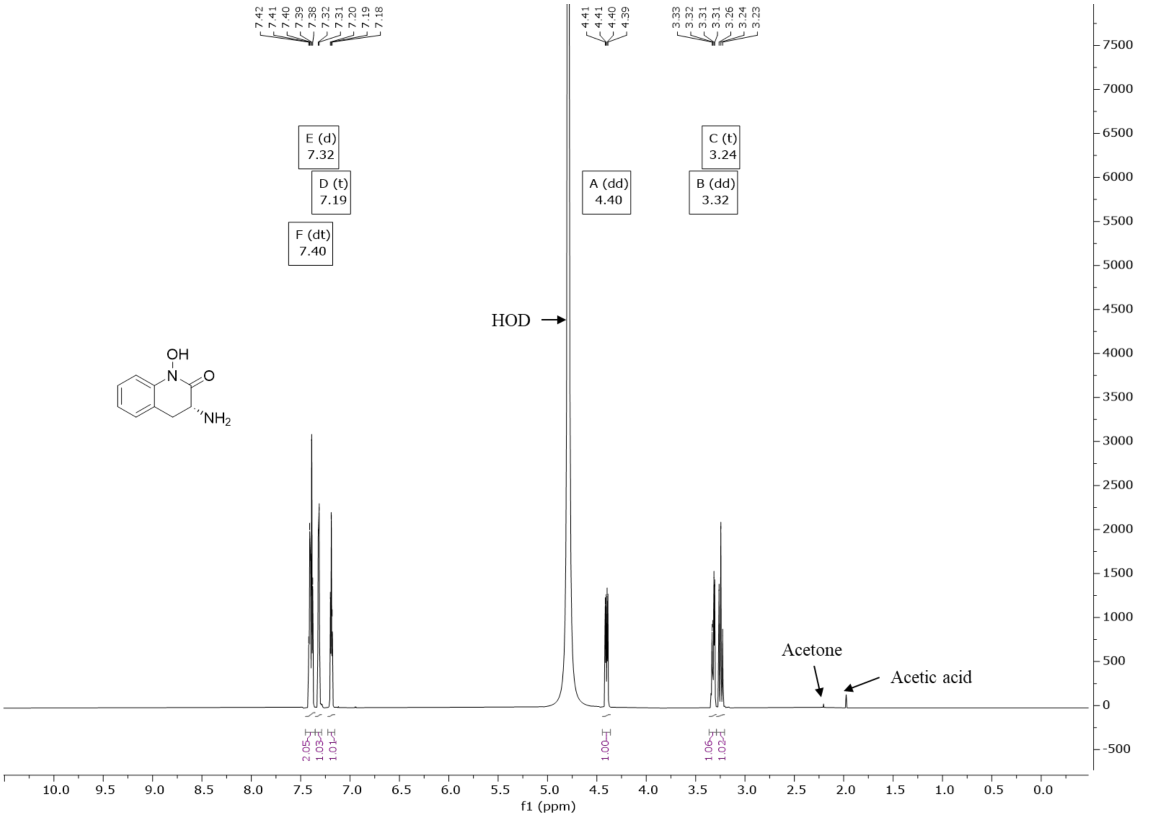


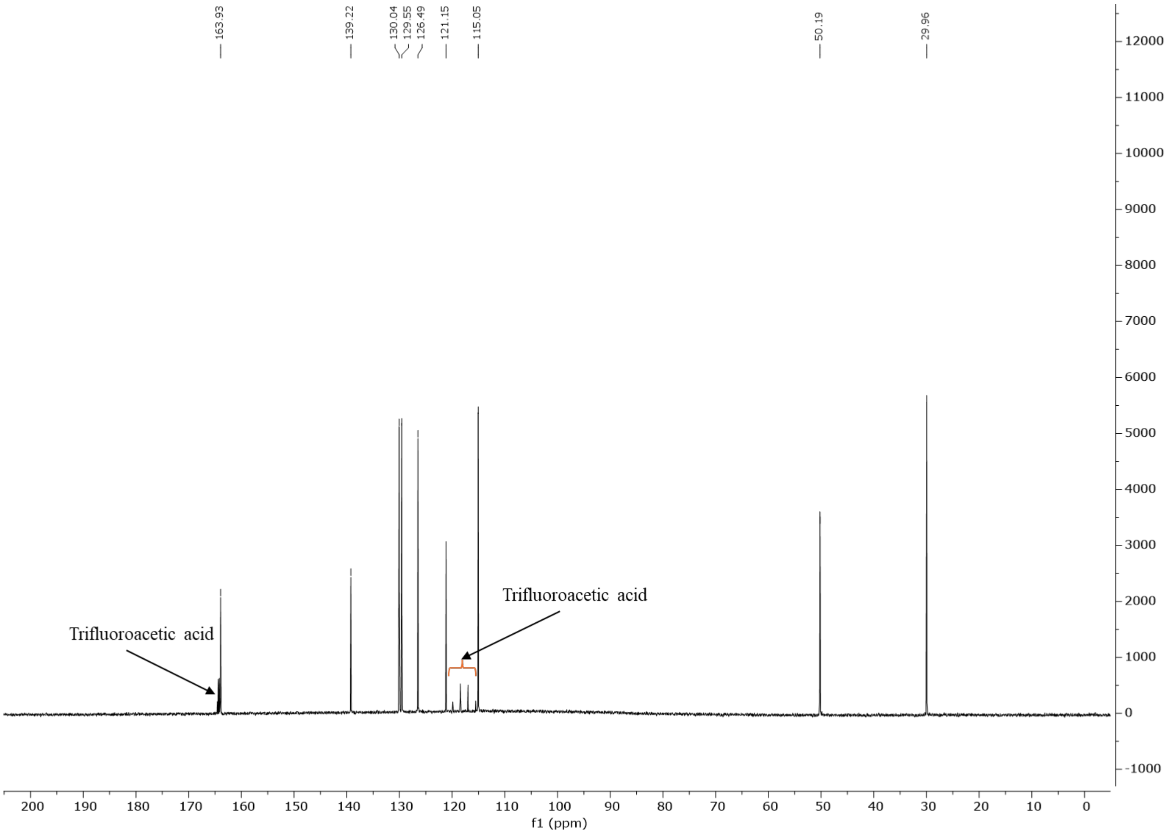

### ESI-MS spectrum and HPLC profile of compound – TI-656

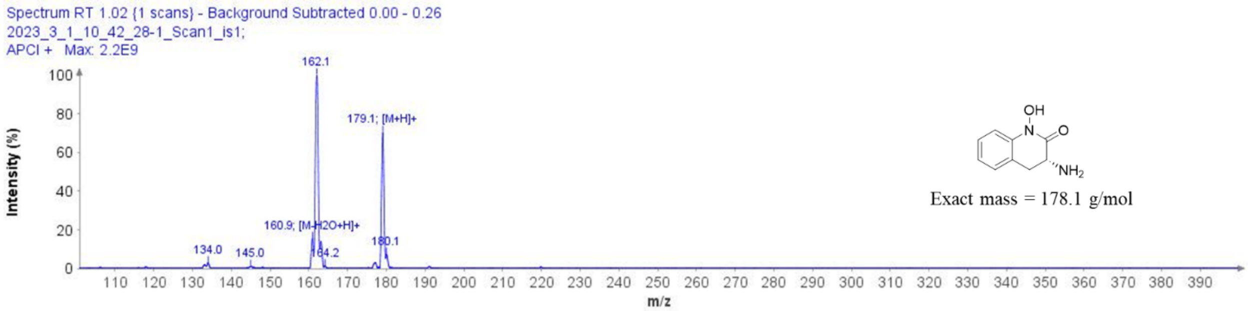


Exact mass = 178.1 g/mol

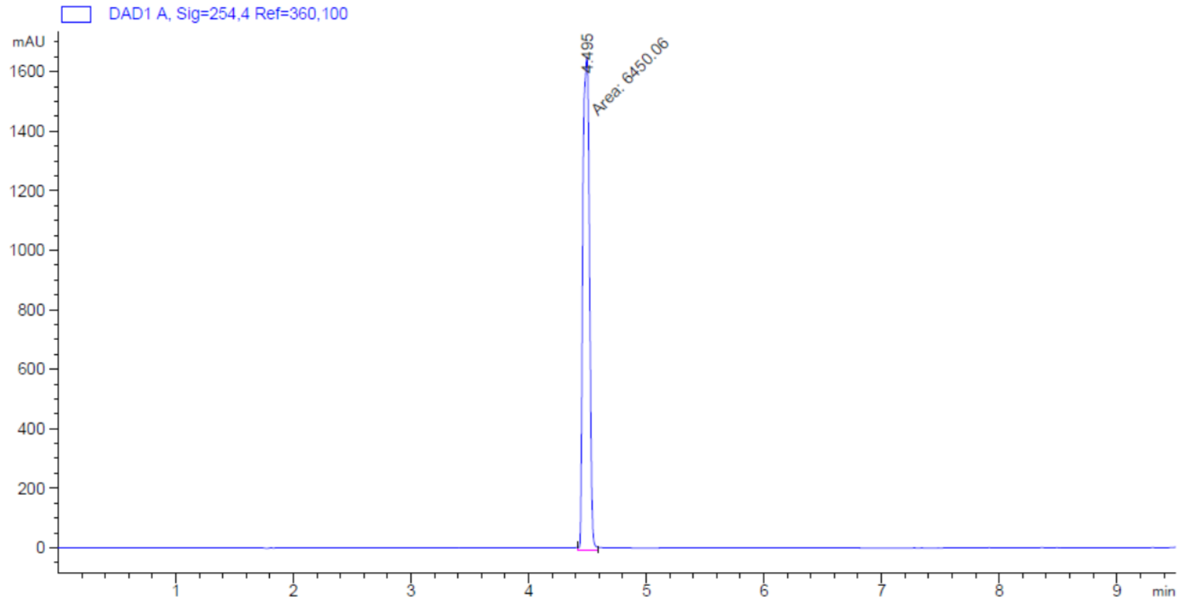


Signal 1: DAD1 A, Sig=254,4 Ref=360,100

| Peak # | RetTime [min] | Type | Width [min] | Area [mAU*s] | Height [mAU] | Area % |
|---|---|---|---|---|---|---|
| 1 | 4.495 | MM | 0.0647 | 6450.06445 | 1660.51746 | 100.0000 |

Totals : 6450.06445 1660.51746

$^{1}H$ and $^{13}C$ NMR spectra of compound – TI-772

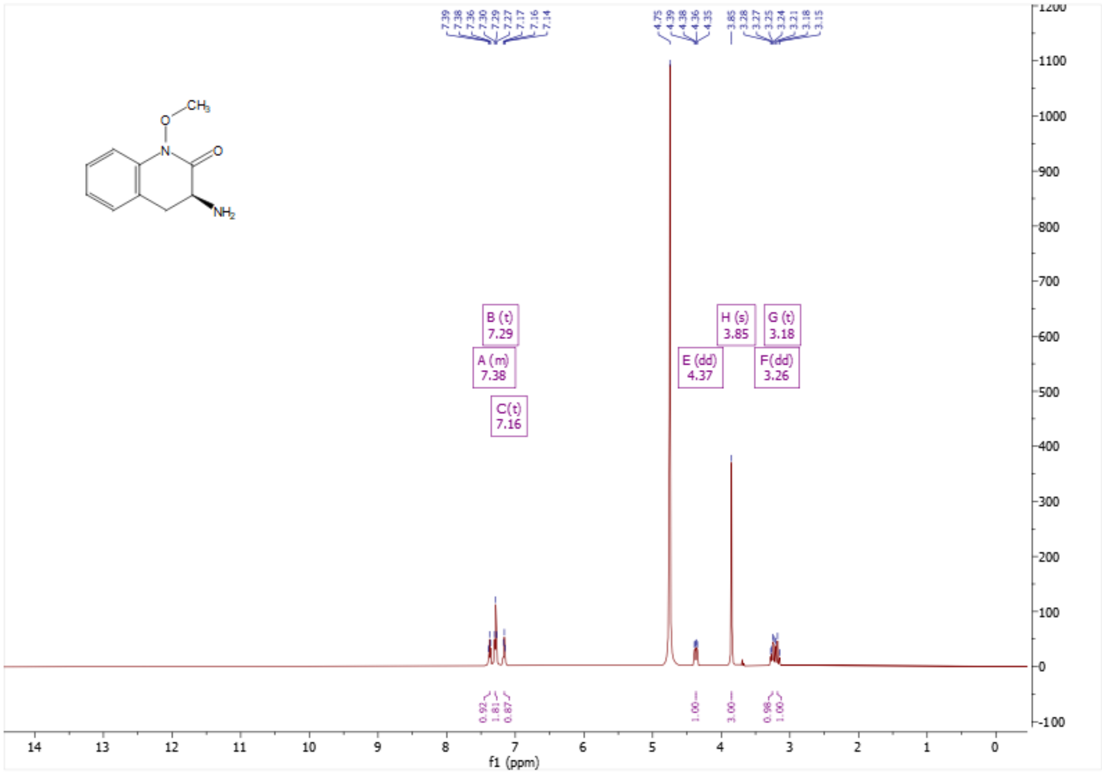

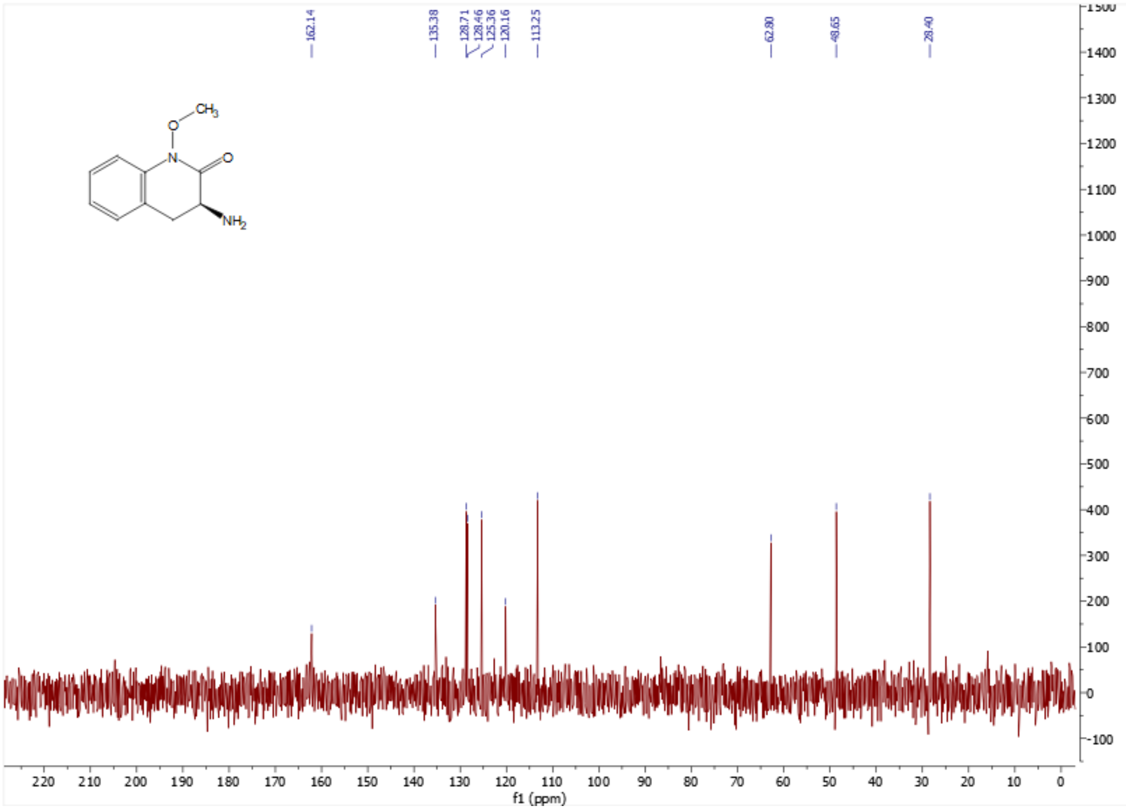

#### ESI-MS spectrum and HPLC profile of compound – TI-772

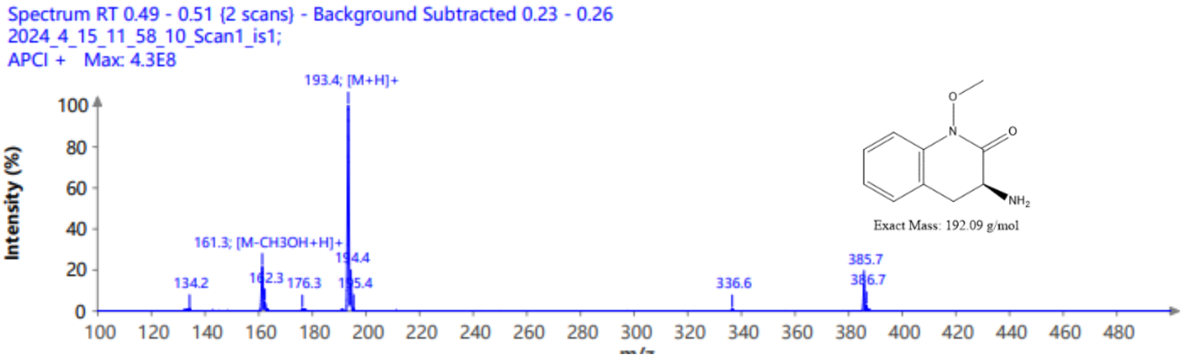


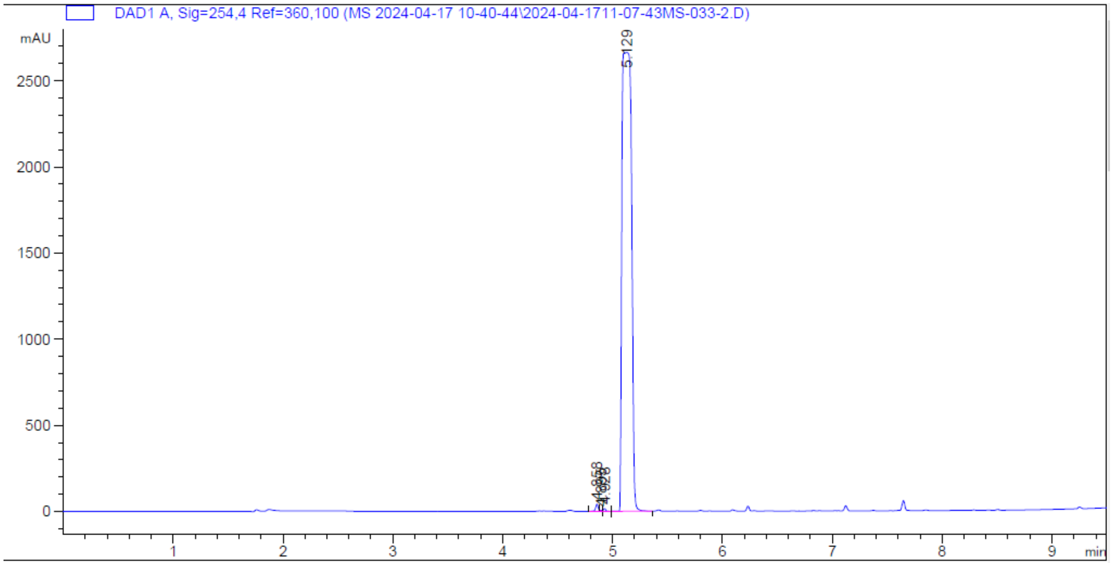


Signal 1: DAD1 A, Sig=254,4 Ref=360,100

| Peak # | RetTime [min] | Type | Width [min] | Area [mAU*s] | Height [mAU] | Area % |
|---|---|---|---|---|---|---|
| 1 | 4.858 | BV R | 0.0255 | 70.35204 | 42.17065 | 0.4389 |
| 2 | 4.895 | VV E | 0.0179 | 6.18696 | 5.20030 | 0.0386 |
| 3 | 4.928 | VB | 0.0241 | 23.55146 | 15.17008 | 0.1469 |
| 4 | 5.129 | VV R | 0.0898 | 1.59290e4 | 2663.24414 | 99.3756 |
| Totals : | | | | 1.60291e4 | 2725.78517 | |

### $^{1}H$ and $^{13}C$ NMR spectra of compound – TI-654

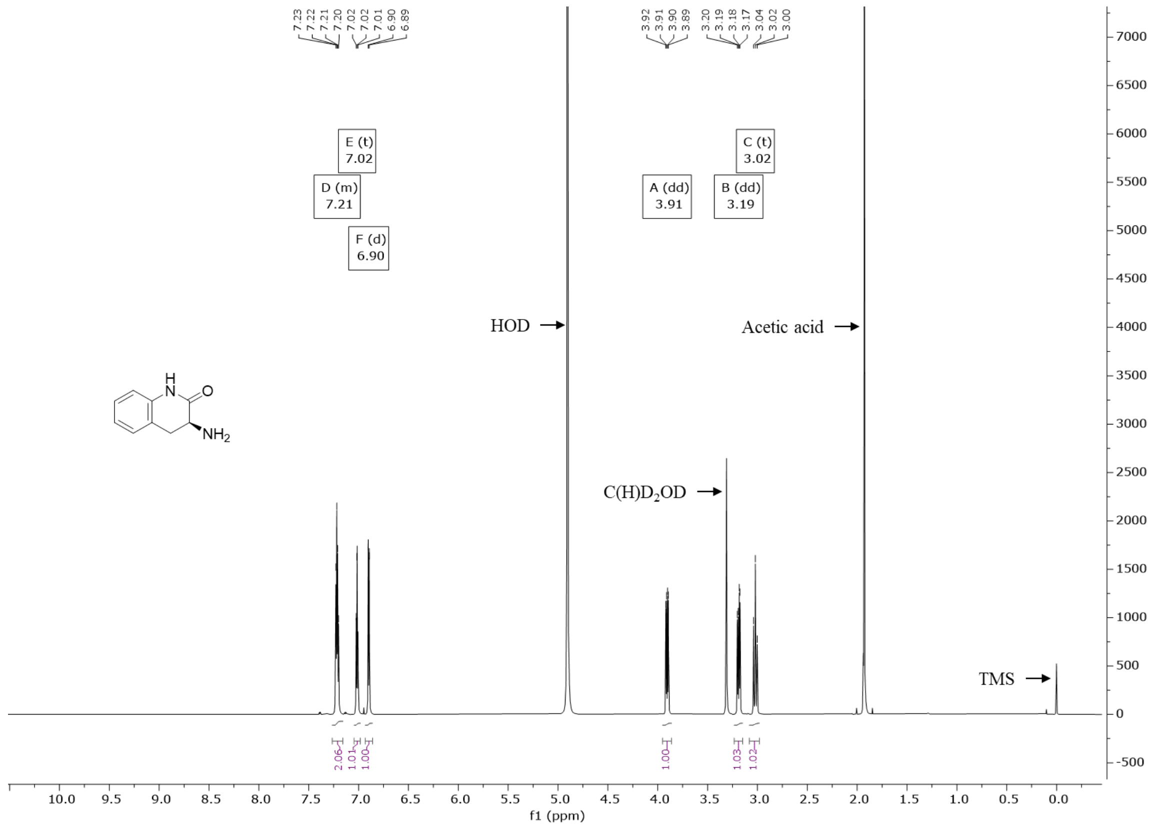


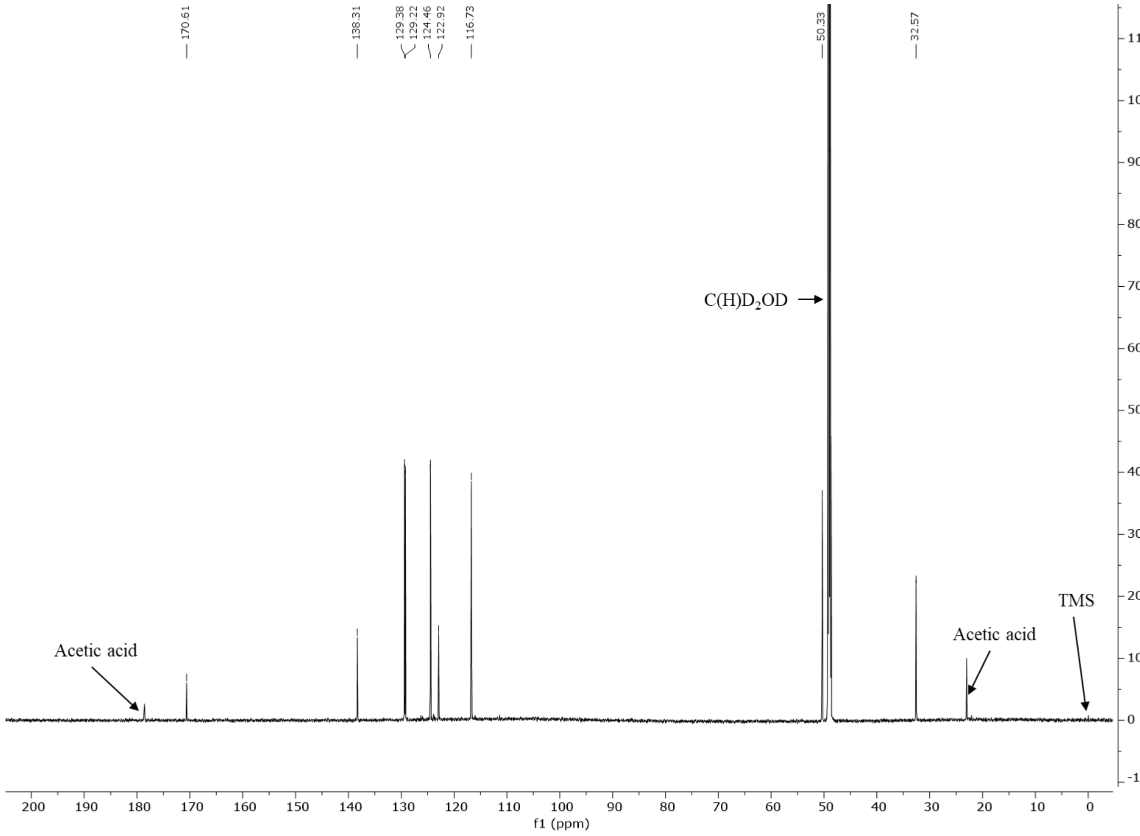

### ESI-MS spectrum and HPLC profile of compound – TI-654

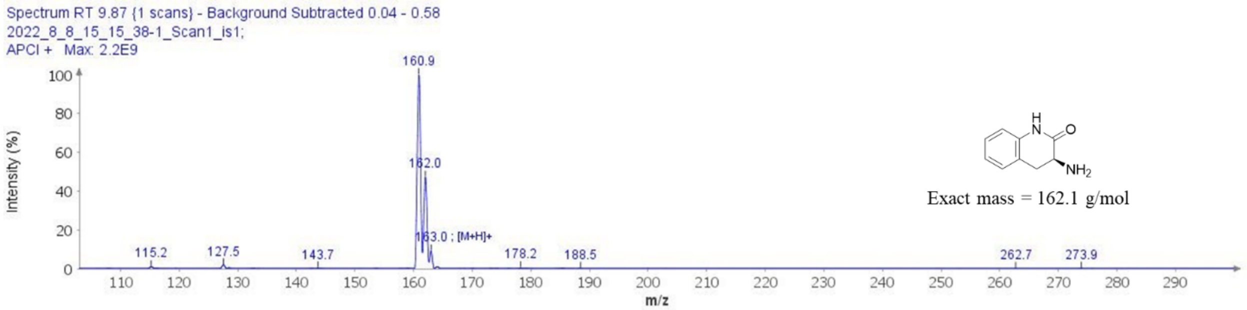


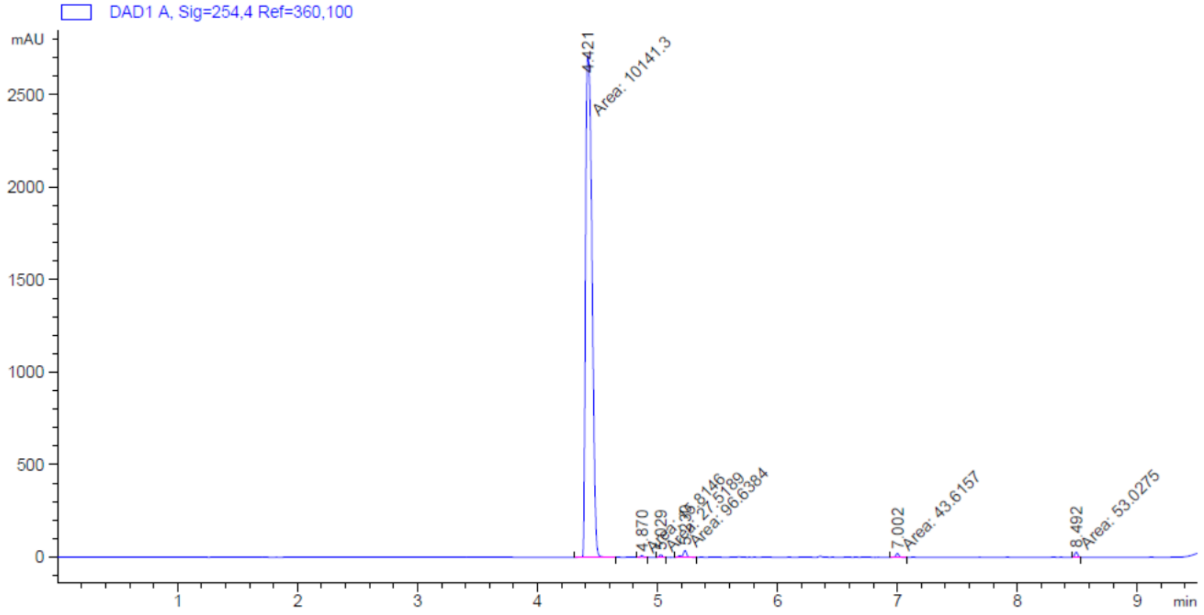


Signal 1: DAD1 A, Sig=254,4 Ref=360,100

| Peak # | RetTime [min] | Type | Width [min] | Area [mAU*s] | Height [mAU] | Area % |
|---|---|---|---|---|---|---|
| 1 | 4.421 | MM | 0.0622 | 1.01413e4 | 2715.67969 | 97.7200 |
| 2 | 4.870 | MM | 0.0308 | 15.81463 | 8.56742 | 0.1524 |
| 3 | 5.029 | MM | 0.0312 | 27.51895 | 14.67853 | 0.2652 |
| 4 | 5.230 | MM | 0.0420 | 96.63844 | 38.34515 | 0.9312 |
| 5 | 7.002 | MM | 0.0327 | 43.61565 | 22.24426 | 0.4203 |
| 6 | 8.492 | MM | 0.0310 | 53.02751 | 28.54146 | 0.5110 |
| Totals : | | | | 1.03780e4 | 2828.05651 | |

### [1]H and [13]C NMR spectra of compound – TI-801

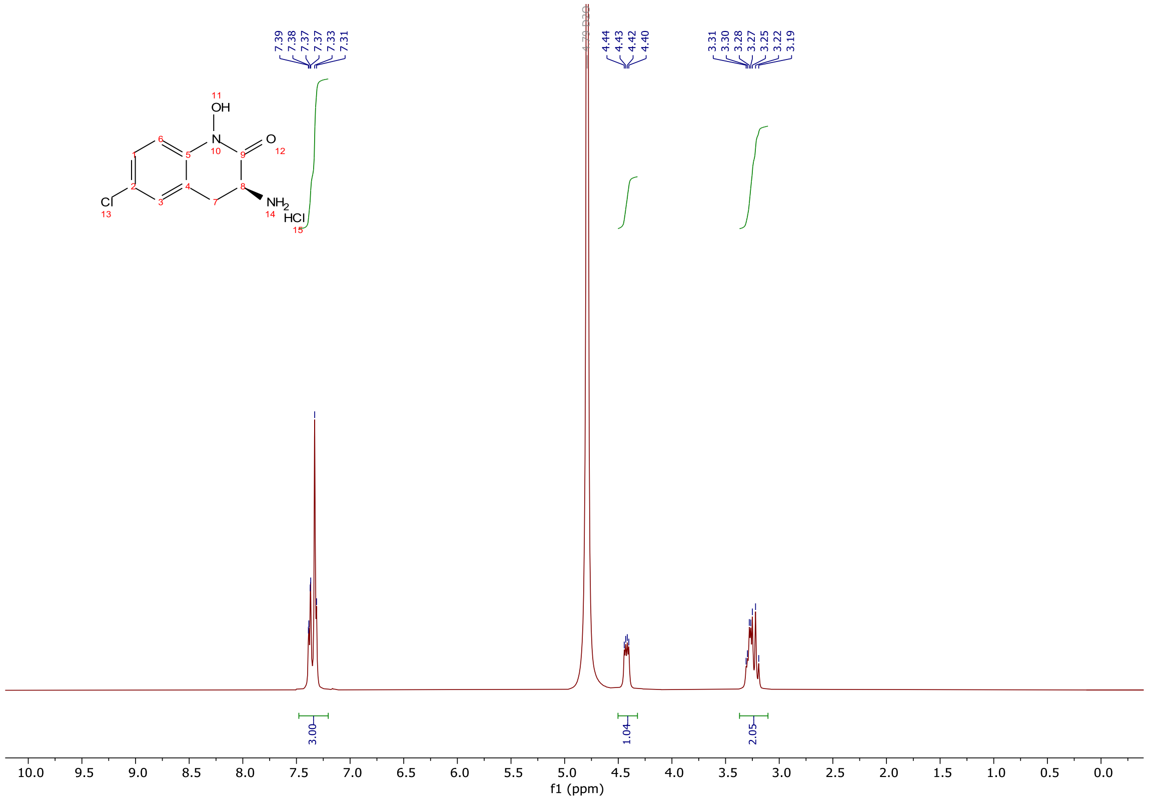

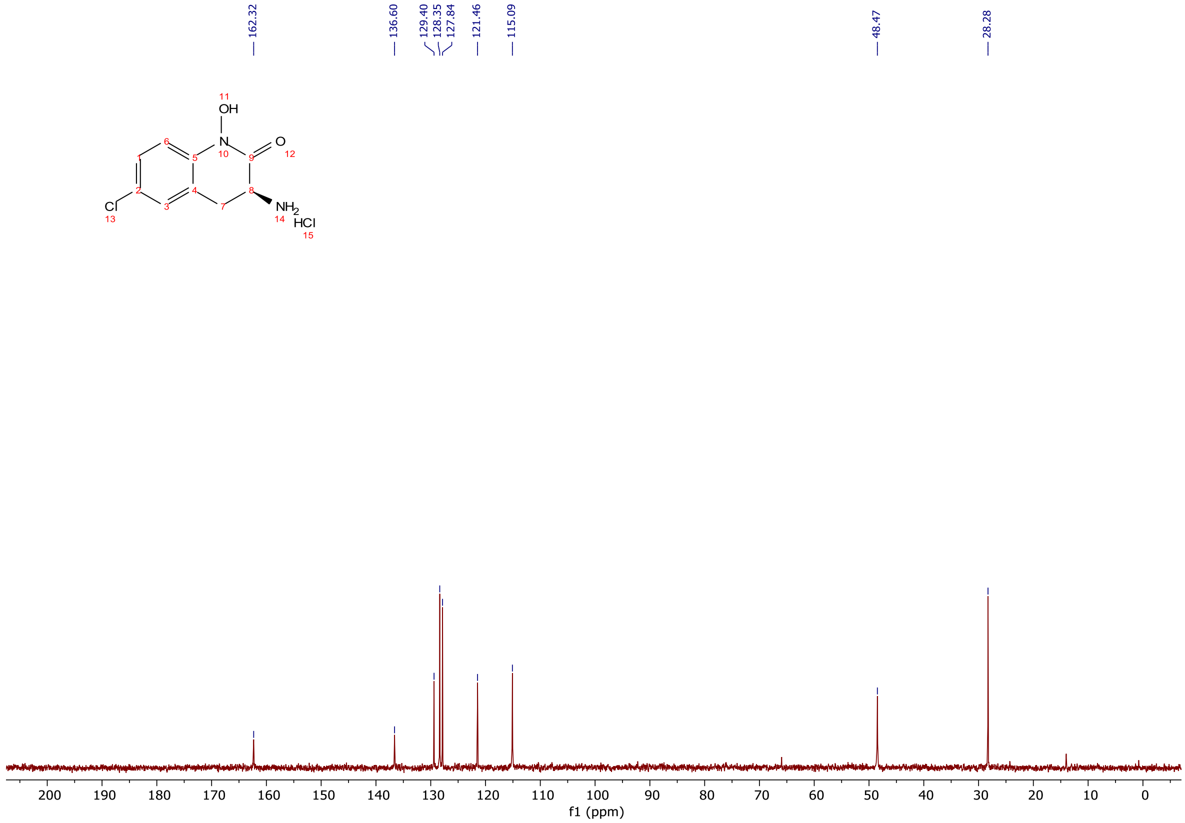

**ESI-MS spectrum and HPLC profile of compound – TI-801**

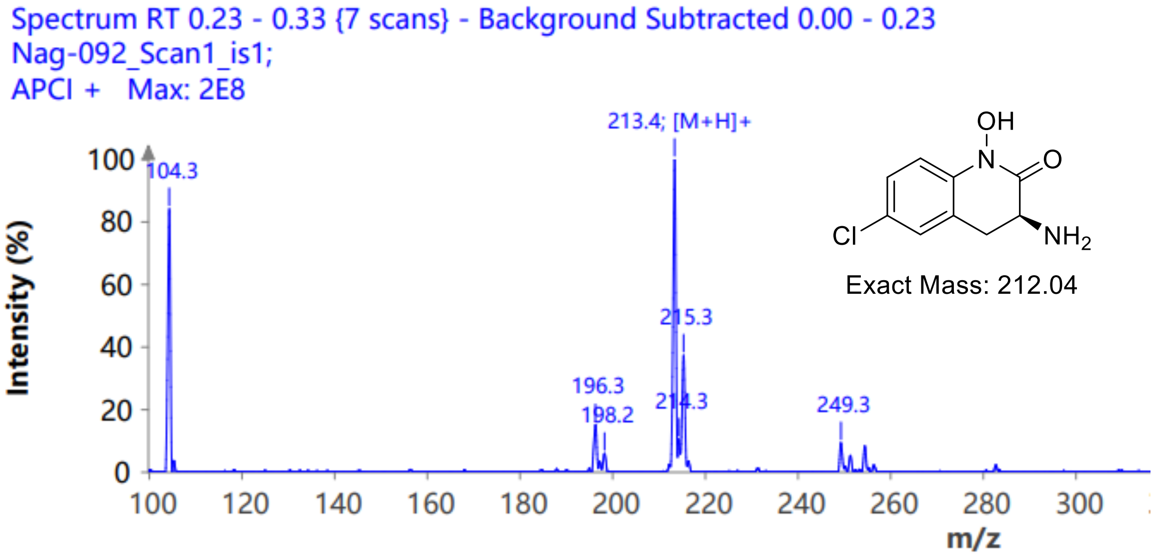


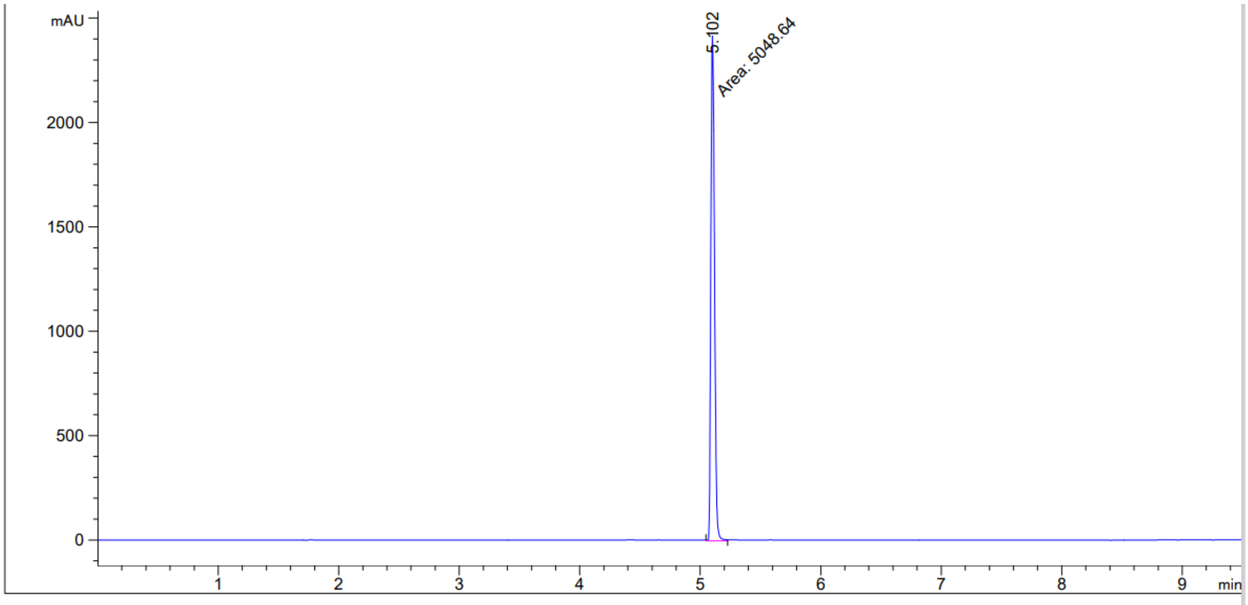


Signal 1: DAD1 A, Sig=254,4 Ref=360,100
Signal has been modified after loading from rawdata file!

| Peak # | RetTime [min] | Type | Width [min] | Area [mAU*s] | Height [mAU] | Area % |
|---|---|---|---|---|---|---|
| 1 | 5.102 | MM | 0.0347 | 5048.63965 | 2423.40991 | 100.0000 |
